# Competition between stochastic neuropeptide signals calibrates the rate of satiation

**DOI:** 10.1101/2023.07.11.548551

**Authors:** Stephen X. Zhang, Angela Kim, Joseph C. Madara, Paula K. Zhu, Lauren F. Christenson, Andrew Lutas, Peter N. Kalugin, Yihan Jin, Akash Pal, Lin Tian, Bradford B. Lowell, Mark L. Andermann

## Abstract

We investigated how transmission of hunger- and satiety-promoting neuropeptides, NPY and αMSH, is integrated at the level of intracellular signaling to control feeding. Receptors for these peptides use the second messenger cAMP, but the messenger’s spatiotemporal dynamics and role in energy balance are controversial. We show that AgRP axon stimulation in the paraventricular hypothalamus evokes probabilistic and spatially restricted NPY release that triggers stochastic cAMP decrements in downstream MC4R-expressing neurons (PVH^MC4R^). Meanwhile, POMC axon stimulation triggers stochastic, αMSH-dependent cAMP increments. NPY and αMSH competitively control cAMP, as reflected by hunger-state-dependent differences in the amplitude and persistence of cAMP transients evoked by each peptide. During feeding bouts, elevated αMSH release and suppressed NPY release cooperatively sustain elevated cAMP in PVH^MC4R^ neurons, thereby potentiating feeding-related excitatory inputs and promoting satiation across minutes. Our findings highlight how state-dependent integration of opposing, quantal peptidergic events by a common biochemical target calibrates energy intake.

## Introduction

Arcuate hypothalamic neurons integrate hormonal^1–4^, interoceptive^5–7^, and external cues^8–12^ and use neuropeptides to regulate food intake, energy expenditure, and body weight^13–18^. In particular, AgRP neurons release hunger-promoting agouti-related peptide (AgRP) and neuropeptide Y (NPY)^19–24^, while POMC neurons release the satiety-promoting α-Melanocyte-stimulating hormone (αMSH) among other peptides^13,25–28^ (Figure 1A). These peptides bind G-protein coupled receptors (GPCRs) that signal by increasing^29^ or decreasing^30^ cAMP concentration. Mutations in these receptor-to-cAMP cascades are associated with obesity in both humans and experimental models^31–40^. One important target of these peptides is the satiety-promoting MC4R-expressing neurons in the paraventricular nucleus of the hypothalamus (PVH^MC4R^)^4,13,14,24,41–46^. How these hunger and satiety peptides are integrated at the level of cAMP, and how cAMP, in turn, controls spiking of PVH^MC4R^ neurons remain largely unknown. One major obstacle has been the inability to directly monitor cAMP levels in individual PVH^MC4R^ neurons *in vivo*, because cAMP dynamics are invisible using conventional electrophysiological or calcium recordings. In the absence of direct cAMP measurements, past studies – which have largely relied on indirect measurements and pharmacology – have generated conflicting results regarding the involvement of cAMP in regulating PVH^MC4R^ neuron activity and hunger^47–54^. There is a critical need to assess neuropeptide release and intracellular signaling dynamics in the awake brain, where neurons are surrounded by many competing peptides whose concentrations vary across behavioral states.

**Figure 1.**
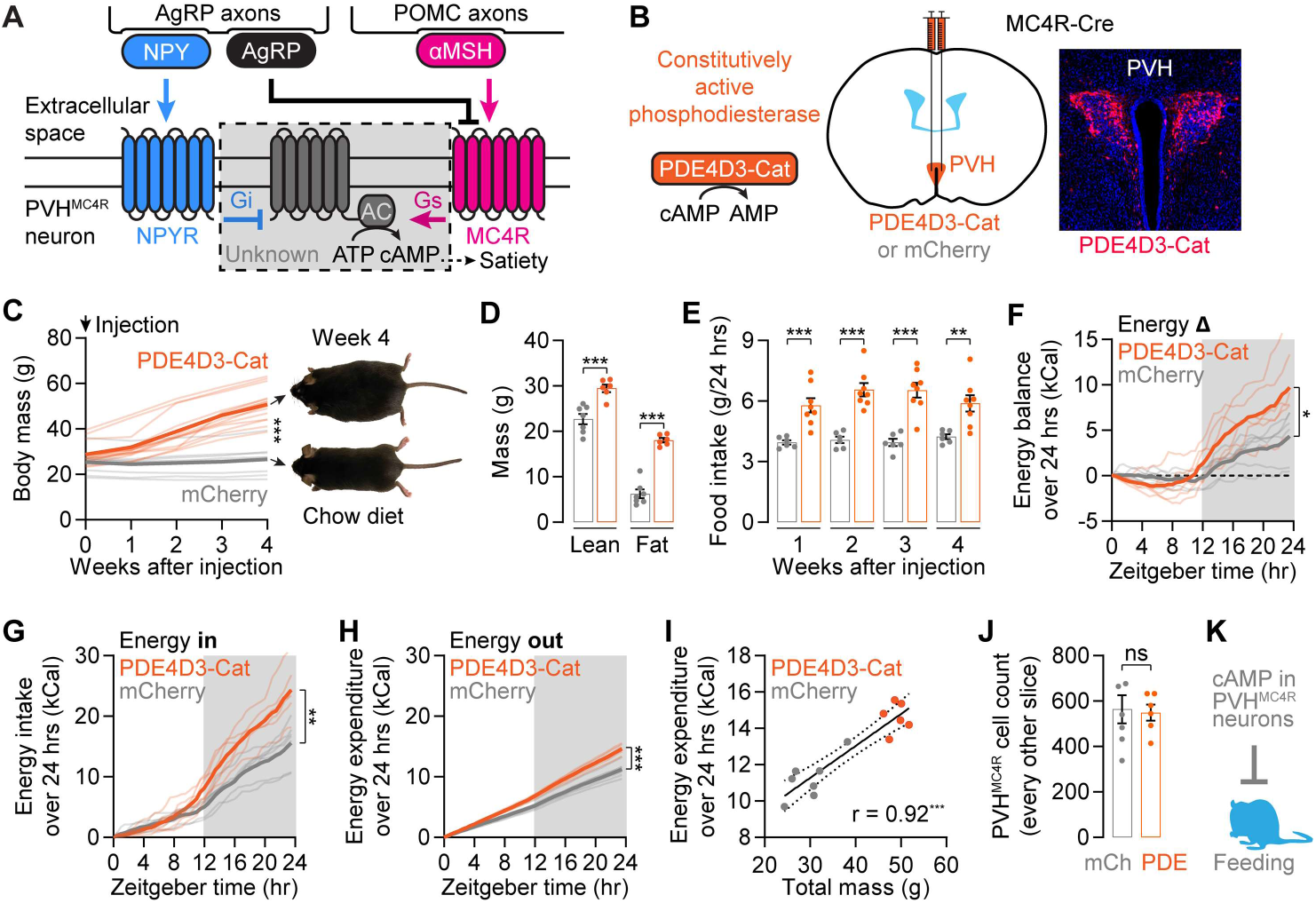
Accelerating cAMP degradation in PVH^MC4R^ neurons results in hyperphagia-induced obesity. **(A)** Model: Bidirectional regulation of cAMP signaling in satiety-promoting PVH^MC4R^ neurons by hunger and satiety peptides. **(B-C)** AAV expression of PDE4D3-Cat, a constitutive cAMP PDE, in PVH^MC4R^ neurons (B) results in drastic weight gain when compared to mCherry controls (C; n = 9-15 mice, t-test). **(D)** Mice that express PDE4D3-Cat have elevated lean mass and fat mass (n = 6-7 mice, one-way ANOVA). **(E)** PDE4D3-Cat expression causes elevated 24-hr food intake (n = 6-8 mice, one-way ANOVA). **(F-H)** Over 24 hours, mice that express PDE4D3-Cat show a net energy surplus (F) that is driven by a large increase in energy intake (G) that is partially offset by elevated energy expenditure (H). n = 6-7 mice, t-test. **(I)** Correlation between energy expenditure and total mass in PDE4D3-Cat-expressing and control mice (n = 6-7 mice, Pearson correlation). **(J)** PDE4D3-Cat expression does not affect PVH^MC4R^ cell counts (n = 6-7 mice, t-test). Model: cAMP signaling in PVH^MC4R^ neurons is important for reducing feeding. n.s. not significant, *p < 0.05, **p < 0.01, ***p < 0.001 for all figures. See Table S1 for statistical details and sex-specific characterizations.

To overcome these technical challenges, we have recently assembled a suite of tools to stimulate and track endogenous neuropeptide release from AgRP and POMC neurons, as well as to measure and manipulate cAMP levels in individual PVH^MC4R^ neurons, in awake mice across seconds to hours^55^. Neuropeptides are found in dense-core vesicles that sparsely populate chemical synapses^56^. Once released, these neuropeptides could potentially diffuse across long distances to signal downstream neurons^56–59^. However, many questions regarding this process remain, particularly in the awake brain. How frequently are the neuropeptides released, and how far do they travel? What are the timescale and state-dependence of peptidergic effects on downstream neurons? Do multiple neuropeptides act on the same intracellular signals? How do changes in intracellular signaling impact neural activity and behavior?

In this paper, we investigate the integration and competition of peptide-evoked biochemical signals in awake mice, and ultimately describe how this biochemical computation leads to gradual changes in synaptic strength and feeding behavior (Figure 1A). Specifically, we show that NPY release from AgRP axons and αMSH release from POMC neurons decrease and increase cAMP in PVH^MC4R^ neurons, respectively. These peptides are released stochastically, and each release event triggers spatially localized subcellular changes in cAMP levels. For both peptides, cAMP changes were larger and more persistent when levels of opposing peptides were low: NPY-mediated signaling was enhanced in fasted mice (when αMSH tone is low), while αMSH-mediated signaling was enhanced in fed mice (when NPY tone is low). Repeated food consumption resolves this competition by driving αMSH release while inhibiting NPY release, resulting in a steady state increase in cAMP that gradually promotes feeding-evoked responses in PVH^MC4R^ neurons over minutes and thereby suppresses appetite. Based on these findings, we propose a model of biochemical computation through which opposing, probabilistic neuropeptide signals are integrated over time to regulate behavior, and suggest ways in which this model could guide future anti-obesity strategies.

## Results

### Blocking cAMP signaling in PVH^MC4R^ neurons causes profound obesity through hyperphagia but not decreased energy expenditure

Previous biochemical and ultrastructural characterizations of feeding-related neuropeptide receptors show converging signaling through cAMP^4,29,30,36,41,60^ (Figure 1A). However, knocking out the relevant G proteins in PVH^MC4R^ neurons from birth has little to no effect on feeding^53,61^, and alternative signaling mechanisms have been suggested^41,47,54,62^. To assess the consequence of increased cAMP clearance on energy balance, we expressed a constitutively active phosphodiesterase (PDE), PDE4D3-Cat^55^, in PVH^MC4R^ neurons in adult mice (Figure 1B). Injection of AAV expressing PDE4D3-Cat at eight weeks of age caused mice to increase body weight up to 80% compared to littermates within four weeks (Figures 1C and S1A, Table S1; standard chow diet). MRI body-composition scanning showed disproportionately increased fat mass in these mice (Figures 1D and S1B), confirming that increasing cAMP clearance in PVH^MC4R^ neurons results in obesity. In addition to obesity, PDE4D3-Cat-expressing mice are also hyperphagic (i.e., overeating) (Figure 1E) and have increased body length (Figure S1C). All three phenotypes are hallmarks of global MC4R knockout mice^41–44^, and because they can be mimicked by accelerating cAMP clearance, our results support the idea that Gαs-coupled MC4R signals via cAMP production^4,29^.

In addition to hyperphagia, decreased energy expenditure could also contribute to the obesity phenotype^63,64^. To investigate the relative contributions of these two causes, we used an indirect calorimetry system to measure energy intake and expenditure independently (CalR^65^) (Figures S1D-S1J). Four weeks after surgery, PDE4D3-Cat-expressing mice accumulated a ∼5 kCal energy surplus per day compared to littermates, consistent with weight gain (Figure 1F). However, instead of positive contributions by both increased energy intake and decreased expenditure, the net energy gain was derived from elevated food intake (∼10 kCal/day) that was insufficiently offset by elevated energy expenditure (∼5 kCal/day) (Figures 1G-1H). The excess energy intake was primarily driven by elevated dark-phase feeding (Figures S1D-E). The elevated energy expenditure, which is calculated from O_2_ and CO_2_ measurements (Figures S1F-S1H), was evident throughout the day (Figures S1I-S1J). Additional analyses indicated that the elevated energy expenditure is well-explained by the elevation in body mass associated with increased food intake (Figures 1I and S1K-S1N). These results indicate that PDE4D3-Cat expression causes obesity primarily through hyperphagia, not decreased energy expenditure. None of the energy-balance changes above were due to loss of PVH^MC4R^ neurons (Figure 1J) or impaired cell health (see electrophysiology recordings in Figure 7). Together with previous studies of PVH^MC4R^ neurons^42–44^, these results support the conclusion that cAMP signaling in PVH^MC4R^ neurons is critical for suppressing feeding. We then set out to characterize how endogenous release of peptides regulates cAMP in these cells *in vivo* (Figure 1K).

### Photostimulating POMC axons triggers all-or-none cAMP increments in a state-dependent manner

αMSH released from satiety-promoting POMC neurons has been proposed to activate MC4R receptors that, in turn, elevate cAMP production in PVH^MC4R^ neurons^4,13,24,26,36,41^ (Figure 2A). However, direct *in vivo* observation of endogenous peptide–induced changes in cAMP or in downstream signaling is lacking. To assess the spatiotemporal dynamics of peptide transmission, we performed two-photon imaging of a cAMP sensor, cADDis^66^, in individual PVH^MC4R^ neurons in awake, head-fixed mice through a GRIN lens^55,67^, together with optogenetic stimulation of POMC axons in the PVH^68^ (Figure 2B). In fasted mice, single-trial photostimulation trains (8 s, 30 Hz, shaded red, Figure 2C) could trigger large increments in cAMP that persisted for tens of seconds in the somas of individual PVH^MC4R^ neurons. However, these large increments occurred only rarely, and on different trials for different PVH^MC4R^ neurons (Figure 2C). The occurrence of a cAMP increment is an all-or-none event with a bimodal distribution of amplitudes that occurs in approximately 15% of trials (Figure 2D; few cAMP decrements were seen during POMC axon stimulation). Because single-trial amplitudes of cAMP increments were bimodally distributed (Figure 2D inset), we used a classifier (see Methods) for automated labeling of trials as ‘hits’ involving large increases in cAMP, or ‘misses’ lacking changes in cAMP.

**Figure 2.**
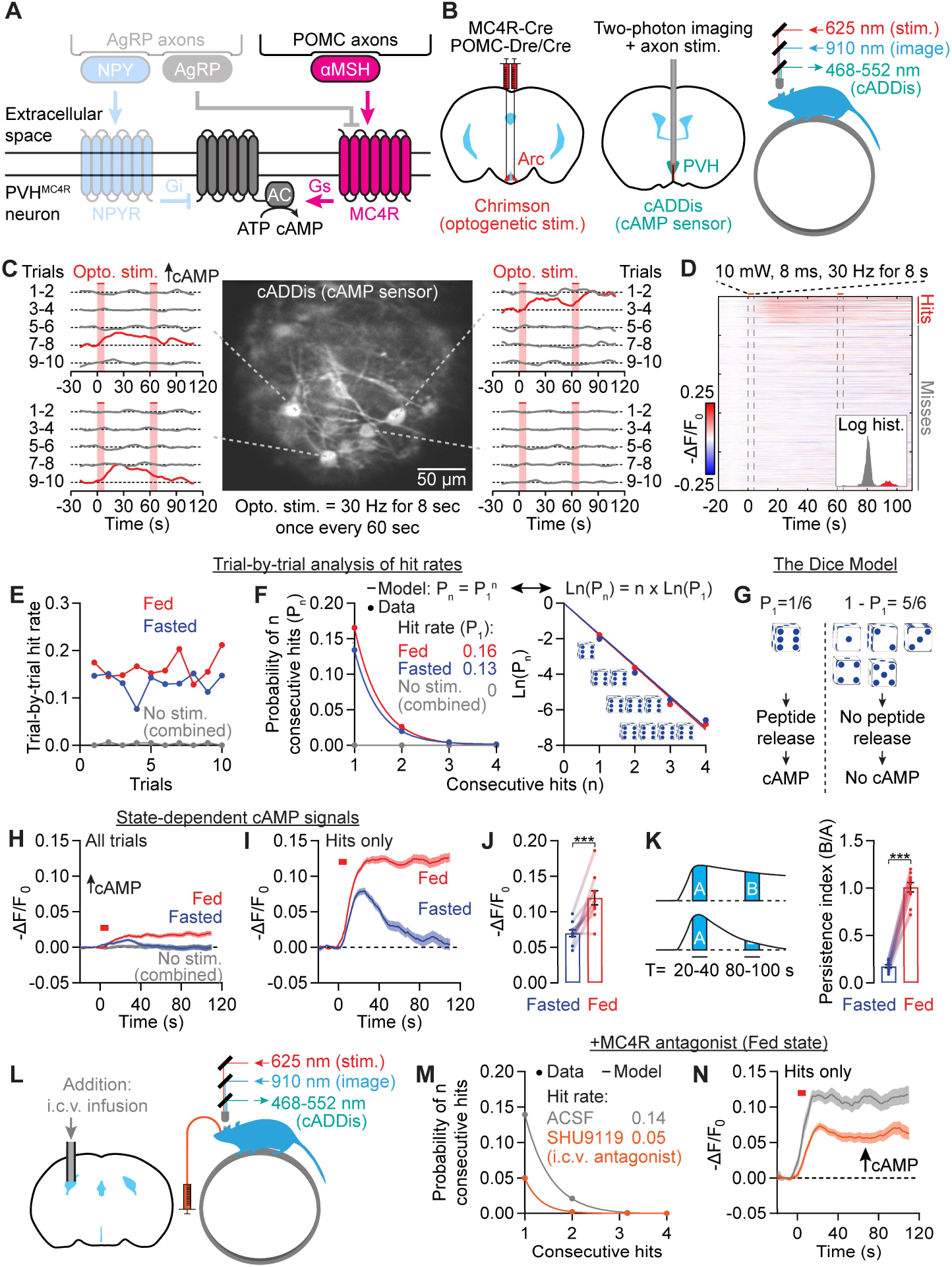
Photostimulation of POMC axons triggers all-or-none, state-dependent cAMP increments in PVH^MC4R^ neurons through αMSH release. **(A)** Model: POMC axon stimulation releases neuropeptide αMSH that subsequently binds to MC4R receptors to activate cAMP production. **(B)** Steps for optogenetic Chrimson stimulation of POMC axons in the PVH in awake, head-fixed mice while measuring cAMP activity in PVH^MC4R^ neurons using the sensor cADDis via a GRIN lens. **(C)** Example cAMP traces during 10 trials of 8 s, 30 Hz stimulation (red shade) with 60 s inter-trial interval show cAMP transients (red traces) on different trials for different cells. Data show a fasted session. Since cADDis fluorescence decreases with higher cAMP, the y-axes are flipped for all cADDis traces (-ΔF/F_0_) to make the plots intuitive. Upward arrows indicate increases in cAMP. **(D)** Summary plot of single-trial cADDis traces shows an all-or-none distribution. Traces are sorted by peak fluorescence changes from 20-40s after stimulation onset. Inset: distribution of peak intensities, color-coded red for hits and gray for misses, with x-axis on a log scale. n = 3407 trials from 4 mice. **(E-F)** Hit rates show no clear trends of increase or decrease (E), and odds of obtaining 2-4 hits in a row are well modeled by a power function (P_n_ = [P_1_]^n^), indicating a fixed hit rate per trial (F). Right side of F: same plot but with y-axis on a log scale. No-stimulation control trials show almost no hits. n = 1571 fasted and 1836 fed trials from 4 mice. **(G)** In the Dice Model, each trial has a fixed chance of triggering a cAMP transient, similar to a die roll stopping on a given number. **(H-I)** Across all trials (H) and for hit trials only (I), cAMP increments are greater in amplitude and more persistent in the fed state than the fasted state (n = 1571 fasted and 1836 fed trials from 4 mice). **(J)** cAMP transient amplitudes are elevated in the fed state (n = 10 FOVs from 4 mice, paired t-test). **(K)** Using a persistence index defined as the ratio of late (80-100 s after stimulation onset) vs. early (20-40 s) areas under the curve, cAMP transients are more persistent in the fed state (n = 10 FOVs from 4 mice, t-test). **(L)** We used intracerebroventricular (i.c.v.) infusion via a 22-gauge cannula to pre-infuse an MC4R antagonist before recording cAMP transients in PVH^MC4R^ during stimulation of POMC axons (see B). **(M-N)** I.c.v. pre-infusion of 1 nmol SHU9119 reduced both hit rate (M) and cAMP transient amplitude (N) in fed mice. n = 931 ACSF and 1757 SHU9119 trials from 2 mice.

The proportion of hit trials, or hit rate, did not monotonically decrease or increase over the course of a session (10 trials, Figure 2E). This argues against neuropeptide vesicle depletion (which should diminish hit rates across trials)^69^ or gradual, use-dependent synaptic facilitation (which should increase hit rates across trials)^70–72^ as causes of the variable occurrence of cAMP increments. Instead, cAMP increments in individual neurons appear to be stochastic, fixed-probability events (potentially due to stochastic exocytosis of individual dense-core vesicles from single POMC axons^73^). The probability that a stimulation train produces a hit could be modeled as rolling a six-sided die (Figure 2F). In such a model, if the probability of rolling a hit (P1) is known, the chance of observing two, three, or n consecutive hits is simply the product of the probabilities (P12, P13, and P1n, respectively). In our data, the probability of observing several consecutive cAMP increments fits such a model (Pn = P1n, also expressed as Ln(Pn) = n * Ln(P1), where Ln is the natural logarithm function; Figure 2F). We therefore used this dice model (Figure 2G) as a framework to analyze stochastic cAMP transients for the rest of this study. We used stimulation-locked windows to classify cAMP hits in order to decrease false-positives (see Methods). As such, we did not assess spontaneous changes in cAMP (e.g., during no-stimulation sessions [gray traces] in Figures 2E-2F).

We compared cAMP responses to POMC axon stimulation in *ad libitum* fed (‘fed’) and chronically fasted (‘fasted’) mice. Although the frequencies of cAMP transients did not differ between fed and fasted states (Figures 2E-2F, S2A), cAMP increments were much more prolonged in the fed state (Figures 2H-2I and S2B). This difference was evident regardless of whether we averaged across all trials (Figure 2H) or only hit trials (Figure 2I). POMC axon-evoked cAMP increments showed ∼80% larger peak amplitudes (Figure 2J) and longer durations (>5 min vs ∼100 s; Figure S2B) in the fed state than in the fasted state. Accordingly, persistence (calculated as the ratio of the mean cAMP response magnitudes from 20-40 s and from 80-100 s after stimulus onset) was five-fold greater in fed mice (Figure 2K). Together, these results demonstrate satiety state-dependent enhancement in the efficacy of second messenger signaling downstream of POMC axon activation.

Although the above results are consistent with αMSH-MC4R mediated cAMP increments (Figure 2A), POMC neurons also release other neuropeptides and small-molecule neurotransmitters that do not act on MC4R^27,28,73,74^. We therefore tested whether intracerebroventricular (i.c.v.) pre-infusion of the MC4R antagonist SHU9119^75^ affects POMC stimulation–evoked cAMP increments (Figures S2C-S2D). Using two-photon fluorescence lifetime imaging microscopy (2p-FLIM), a photobleaching-insensitive imaging technique that is well-suited for tracking sensor fluorescence across slow timescales^76^, we validated the setup by verifying that i.c.v. infusion of αMSH (1 nmol in 2 μl, see Methods) induced a robust increase in cAMP in PVH^MC4R^ neurons over ∼8 min (Figures S2E-S2G). When we pre-infused SHU9119 (1 nmol in 2 μl) in fed mice (Figure 2L), it blocked ∼60% of POMC stimulation-induced cAMP increments (Figures 2M and S2H) and reduced the amplitudes of the remaining hits by 50% (Figures 2N and S2I-S2K). SHU9119 also blocks MC3R signaling^75^. However, to our knowledge, αMSH is the only neuropeptide released from POMC neurons that binds MC4R at high affinity^27,77^, and therefore likely mediates a majority of POMC-to-PVH^MC4R^ cAMP signaling. Taken together, these results support a model of peptidergic transmission in which stimulation of POMC axons triggers MC4R-dependent, stochastic cAMP increments in PVH^MC4R^ neurons with signaling kinetics that vary substantially across fasted and fed states.

### Photostimulating AgRP axons triggers all-or-none, state-dependent cAMP decrements

Hunger-promoting AgRP neurons also project to PVH, where they release the neuropeptides NPY and AgRP to suppress cAMP (Figure 3A)^13,20,24,30,41,75,78,79^. In contrast to POMC axon-evoked cAMP increments, photostimulation of AgRP axons (8 s, 30 Hz) evoked cAMP decrements in PVH^MC4R^ neuron somas (Figures 3B-3C). These decrements were also stochastic, all-or-none events (Figure 3D) that occurred in 25% of trials (Figures 3C-3D). Similar to POMC stimulation-evoked cAMP increments, the probability of occurrence of these AgRP-evoked cAMP decrements was also well-modeled by the dice model (Figure 3E), and the hit rate was not strongly modulated by hunger state (Figure S3A). However, cAMP decrements also showed state dependence, as they were greater in amplitude and more persistent in fasted than in fed mice (Figures 3F-3I and S3B).

**Figure 3.**
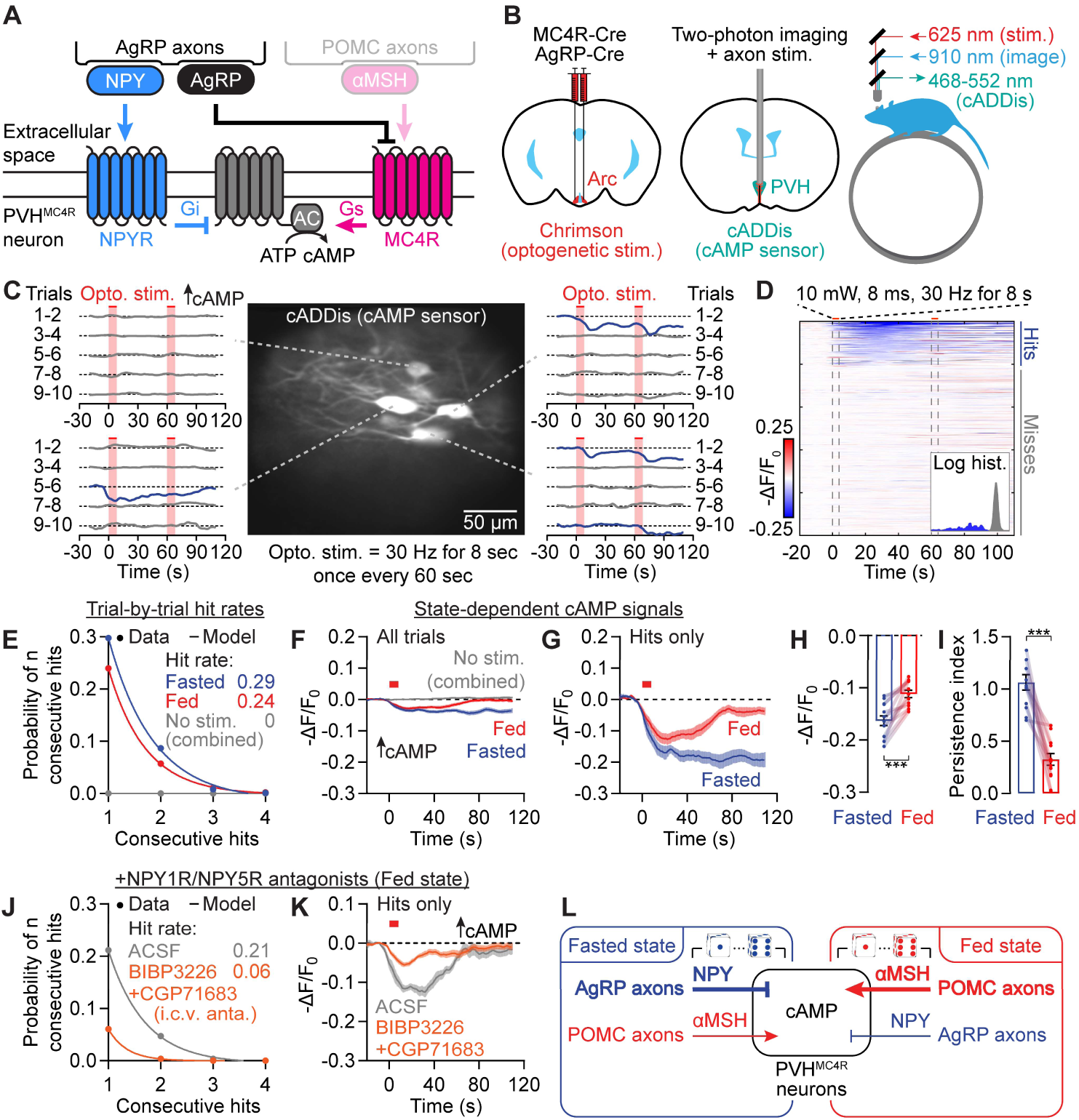
Photostimulation of AgRP axons triggers all-or-none, state-dependent cAMP decrements in PVH^MC4R^ neurons through NPY release. **(A)** Model: AgRP axon stimulation could decrease cAMP production by releasing neuropeptide NPY that binds to NPY receptors and/or by releasing AgRP that prevents MC4R receptor agonism. **(B)** Setup for optogenetic Chrimson stimulation of AgRP axons in the PVH in awake, head-fixed mice while measuring cAMP activity in PVH^MC4R^ neurons using the sensor cADDis via a GRIN lens. **(C)** Example cAMP traces during 10 trials of photostimulation (8 s, 30 Hz, 60 s inter-trial interval; red shade) from a fed session show cAMP decrements (blue traces) that occurred on different trials for different cells. Traces show –ΔF/F_0_. **(D)** Summary plot of single-trial cADDis traces across cells shows an all-or-none distribution. Traces are sorted by peak fluorescence decrements in the 20-40 s window. Inset shows distribution of peak intensities, color-coded blue for hits and gray for misses, with the x-axis on a log scale. n = 2283 trials from 4 mice. **(E)** Odds of obtaining 2-4 hits (decrements) in a row are well modeled by the dice model (P_n_ = [P_1_]^n^). n = 1190 fasted and 1093 fed trials from 4 mice. **(F-G)** In all trials (F) and in hit trials only (G), cAMP decrements are greater in amplitude and more persistent in the fasted state (n = 1190 fasted and 1093 fed trials from 4 mice). No-stimulation control trials show almost no hits. **(H-I)** cAMP decrements show greater amplitude (H) and persistence (I) in fasted than in fed mice (n = 13 FOVs from 4 mice, paired t-test). **(J-K)** I.c.v. pre-infusion of 10 nmol BIBP3226 (NPY1R antagonist) and 10 nmol CGP71683 (NPY5R antagonist) reduced both the hit rate (J) and amplitude (K) of cAMP decrements in fed mice. n = 680 ACSF and 909 antagonist trials from 2 mice. **(L)** Model: NPY release from AgRP axons and αMSH release from POMC axons respectively trigger stochastic and state-dependent cAMP increments and decrements in PVH^MC4R^ neurons.

Out of all the neuropeptides and small-molecule neurotransmitters that AgRP neurons could release (e.g., GABA, AgRP, NPY), NPY mediates a significant proportion of the hunger-promoting effects, especially the slow effects that last many minutes^20^. Therefore, we tested whether NPY is the main mediator of AgRP-to-PVH^MC4R^ cAMP signaling (Figure S3C-S3D; see Figures S3E-S3F for infusion validation). We pre-infused antagonists for two NPY receptors (10 nmol BIBP3226^80^ for NPY1R and 10 nmol CGP71683^81^ for NPY5R, 4 μl total, i.c.v.) that are known to be important for mediating the hunger-promoting effects of NPY^80–84^. In fed mice, these antagonists blocked ∼75% of hits and reduced hit amplitudes by ∼60% (Figures 3J-K and S3G-S3J). These findings show that NPY mediates ∼90% of the influence of AgRP axons on cAMP signaling in PVH^MC4R^ neurons. While release of AgRP peptide, an inverse agonist of the MC4R^85^, might also play a more minor or redundant role, our experiments below focused on the release dynamics and impact of NPY.

Together, the above *in vivo* findings show that αMSH transmission from POMC axons and NPY transmission from AgRP axons mediate the majority of evoked increments and decrements in cAMP, respectively (Figure 3L). Further, the amplitude and persistence of cAMP transients during AgRP or POMC stimulation were state-dependent. Below, we investigated the mechanisms of stochasticity in cAMP signaling (Figure 4), the causes of state-dependent differences in persistence of cAMP transients (Figure 5), and the implications of cAMP stochasticity and persistence for PVH^MC4R^ neuron activity and feeding (Figures 6-7).

**Figure 4.**
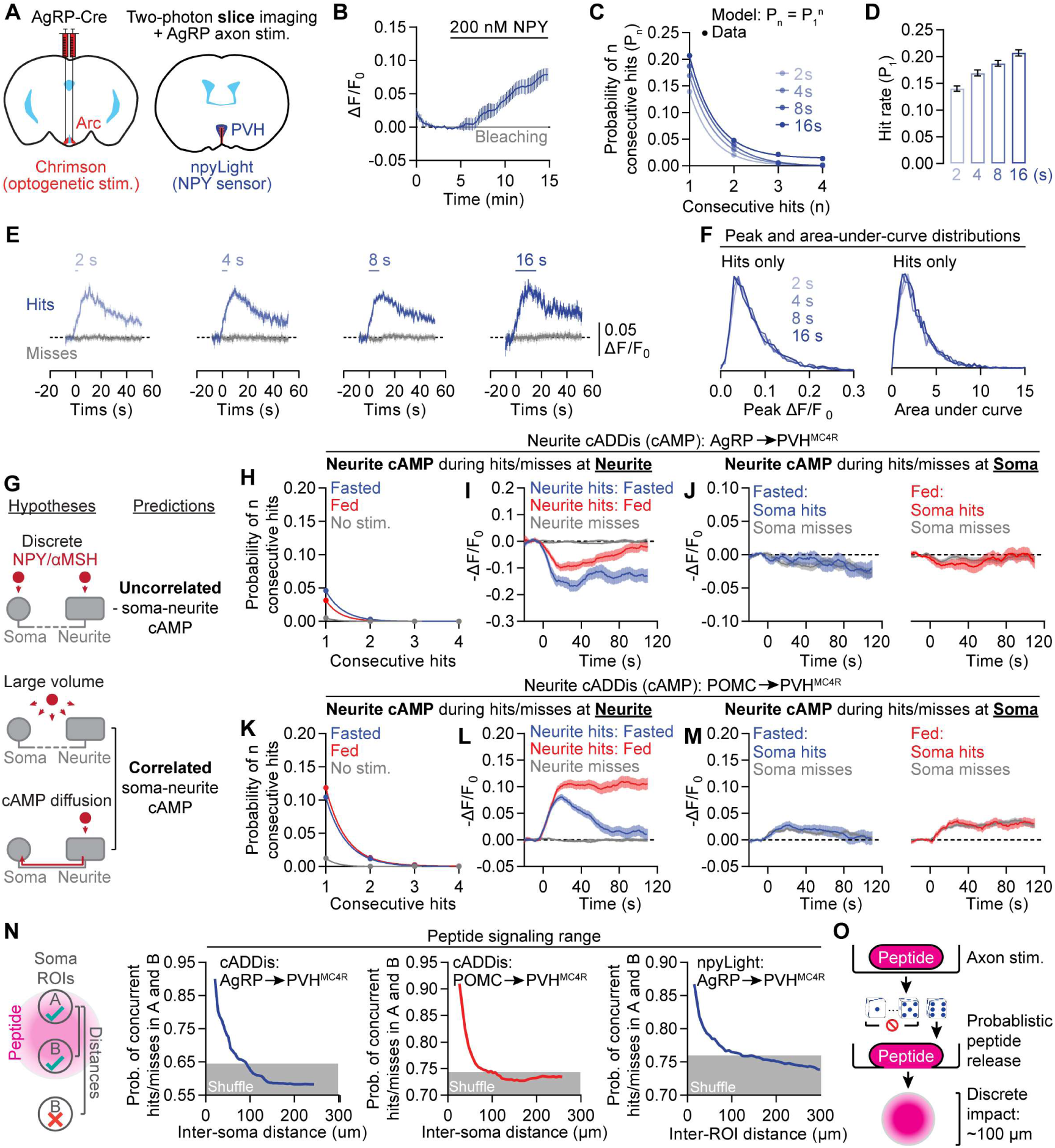
A stochastic and spatially restricted mode of neuropeptide release. **(A)** To measure NPY release in slice, Chrimson is expressed in AgRP neurons to allow axon stimulation in PVH, and npyLight is expressed in the PVH to capture NPY release. These experiments used fed mice. **(B)** The fluorescence of npyLight increases in response to NPY (200 nM; n = 10 slices from 4 mice). **(C-F)** When we increased the duration of 15.5 Hz photostimulation eight-fold, from 2 s to 16 s, the hit rate of npyLight transients increases from 14% to 21% (C-D, n = 16439-19969 trials from 7 mice), and hit amplitudes remain unchanged (E-F) (n = 2690, 2293, 2871, 2982 hits). **(G)** If neuropeptide signaling is spatially discrete (‘Discrete’), hits and misses should be uncorrelated between pairs of regions-of-interest (ROIs) from a soma and distant neurite belonging to the same cell. If hits involve neuropeptide release that diffuses across a large volume (“Large volume”), or if cAMP could diffuse intracellularly (‘cAMP diffusion’), hits and misses should be correlated for pairs of ROIs from the same cell. **(H-I)** During AgRP axon stimulation, neurite cAMP decrements are also stochastic (H) and strengthened in the fasted state (I). n = 6048-9371 trials from 4 mice. **(J)** Neurite cAMP signals averaged across trials containing *soma* hits or *soma* misses are similar and smaller in magnitude, indicating decoupling of soma and neurite signals. n = 2379 fasted and 3138 fed trials from 4 mice. **(K-M)** During POMC axon stimulation, neurite cAMP increments are also stochastic (K) and are larger and more sustained in the fed state (L). Neurite traces averaged across trials containing *soma* hits or *soma* misses were similar and smaller in magnitude, again indicating decoupling of soma and neurite signals (M). K-L: n = 4356-5122 trials from 4 mice. M: n = 1350 fasted and 1363 fed trials. **(N)** By calculating the probability of concurrent hits or concurrent misses in two somas (left model), we found that correlation decreases as the distance between the two increases. The distance beyond which the probability of concurrent hits reaches chance levels is consistently ∼100 μm for cADDis recordings during AgRP (left) and POMC stimulation (middle). The same ∼100 μm distance threshold is also seen with inter-ROI correlations in npyLight recordings during AgRP stimulation (right). See also Figure S4K. **(O)** Stochastic release leads to temporally and spatially discrete neuropeptide signaling.

**Figure 5.**
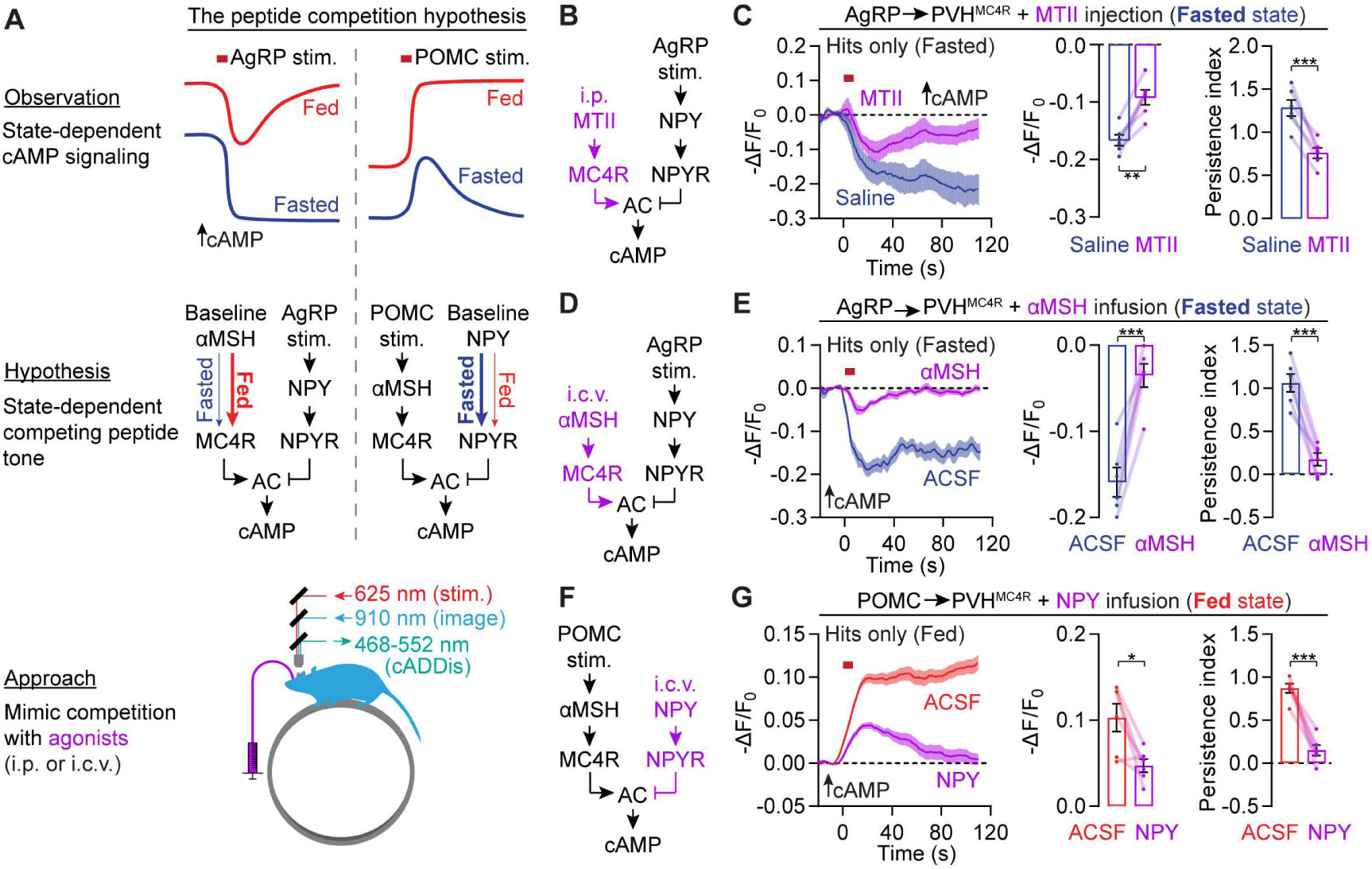
Neuropeptide competition underlies state-dependent cAMP signaling. **(A)** Hypothesis for state-dependent cAMP signaling: In the fed state, elevated basal αMSH levels curb experimentally induced cAMP decrements. Likewise, in the fasted state, increased basal NPY levels curb cAMP increments. This hypothesis is tested by mimicking the presence of competing peptides. **(B-C)** In fasted mice, MTII pre-administration (3 mg/kg, i.p.) attenuated the amplitude and persistence of AgRP stimulation–induced cAMP decrements. n = 95-114 hits and 6 FOVs from 3 mice, paired t-test. **(D-E)** In fasted mice, αMSH pre-infusion (1 nmol, i.c.v.) reduced hit amplitude and persistence of AgRP stimulation–induced cAMP decrements. n = 391-417 trials and 6 FOVs from 2 mice, paired t-test. **(F-G)** In fed mice, NPY pre-infusion (0.5 nmol, i.c.v.) reduced hit amplitude and persistence of POMC stimulation–induced cAMP increments. n = 1311-1667 trials and 6 FOVs from 2 mice, paired t-test.

**Figure 6.**
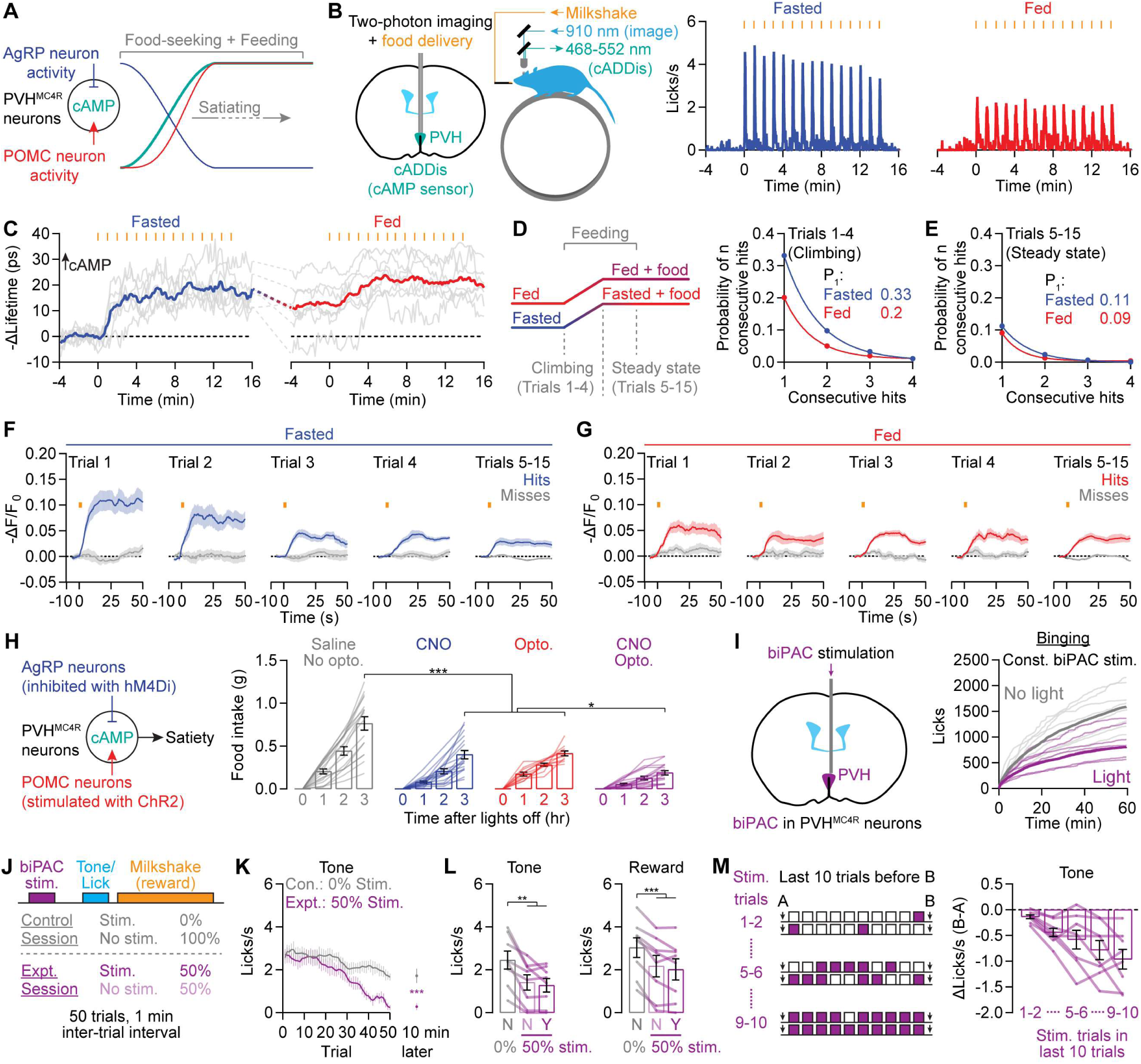
Stochastic and persistent cAMP increments during feeding set the rate of satiation. **(A)** Feeding rapidly decreases AgRP neuron activity while increasing POMC neuron activity, which should result in persistently elevated cAMP. **(B)** Periodic milkshake delivery to head-fixed mice results in transient and time-locked spikes in lick rate that are reliably observed in fasted and fed mice. Lick rates are greater in fasted than in fed mice. n = 6 mice. **(C)** cAMP in PVH^MC4R^ neurons climbs up to a steady state level within ∼4 one-minute trials in both conditions, albeit from a higher initial level in the fed state. Gray traces: mean soma cADDis lifetime changes per FOV. Blue and red traces: mean across FOVs. n = 7-11 FOVs from 6 mice. **(D-E)** In the climbing epoch where cAMP is increasing (Trials 1-4), cAMP increments are stochastic. The hit rate is lower in the steady-state epoch (Trials 5-15) (E). D: n = 561-646 trials from 6 mice, E: 1542-1759 trials. **(F-G)** Persistent cAMP increments in the fasted (F) and fed states (G). F: n = 2103 trials from 6 mice, G: 2405 trials. **(H)** Either chemogenetic inhibition of AgRP neurons (hM4Di and i.p. injection of CNO) or optogenetic stimulation of POMC neurons (ChR2) suppresses dark-phase feeding in freely-moving mice, but the suppression is strongest when the two manipulations are combined (n = 11-17 mice, one-way ANOVA), consistent with decreased peptide competition. **(I)** Continuous low-level (∼15 μW average) stimulation of biPAC in PVH^MC4R^ neurons reduces cumulative lick rate during an *ad libitum* binge assay in head-fixed, fasted mice (n = 8 mice for each condition). **(J)** Experimental design: in control sessions, fasted mice lick during tone to obtain reward (milkshake) with no biPAC stimulation. In experimental sessions, brief biPAC stimulation was delivered 5 s before 50% of the trials, while the other 50% trials did not involve stimulation. In control sessions, no trials contained stimulation. **(K)** Lick rate during tone declines faster in the experimental session and does not recover after 10 min wait (n = 8 mice, t-test). **(L)** Lick rates are lower in experimental sessions, but there is no difference between stimulation and no-stimulation trials within an experimental session (n = 8 mice, one-way ANOVA). **(M)** The decrease in lick rate during tone presentation over 10 trials scales with the average number of biPAC stimulations delivered in the intervening time (n = 8 mice).

### A stochastic and spatially restricted mode of peptide release

The probabilistic nature of the neuropeptide-mediated cAMP signaling described above is redolent of stochastic neurotransmitter release^86^, albeit with the major distinction that the probability of a cAMP transient is low despite bulk stimulation of many AgRP or POMC axons. This low probability is consistent with ultrastructural data showing that most AgRP and POMC presynaptic boutons harbor only a few peptide-containing dense-core vesicles that are sparsely distributed among hundreds of small clear vesicles^73^. To ascertain whether stochasticity in cAMP signaling reflects stochasticity in neuropeptide release, we used viral expression of a new fluorescent NPY sensor, npyLight, in PVH neurons (Figures 4A and S4A). We first confirmed this sensor was sensitive to 200 nM NPY in brain slices (Figure 4B). To induce endogenous NPY release, we used trains of Chrimson photostimulation (15.5 Hz) of varying duration (2, 4, 8, 16 s) to activate AgRP axons in the PVH in brain slices (Figures S4B-S4E). Similar to the stochastic nature of cAMP decrements, AgRP axon photostimulation only triggered npyLight transients in a subset of trials (Figures S4B-S4E). These npyLight transients were mostly well described by the dice model, with an estimated release probability of ∼15% that increased only slightly with stimulation duration (Figures 4C-4D and S4F). Moreover, the amplitude of hits remained consistent for all stimulation durations (Figures 4E-4F). Due to the lack of an αMSH sensor, we could not test these hypotheses in the POMC-to-PVH^MC4R^ cAMP signaling pathway. Nevertheless, these results are consistent with a model in which stochastic peptide release underlies the unpredictability of peptide-mediated cAMP signaling in PVH^MC4R^ neurons.

Whereas fast neurotransmission is mostly restricted to postsynaptic regions proximal to synapses, neuropeptides can diffuse across longer distances^56–58^, sometimes estimated to span entire brain areas^56,87–89^. A recent study using an oxytocin sensor indicated a spatial spread of ∼100 μm in hypothalamic slices^90^, demonstrating a local-diffusion mode of signaling^56^. Similar analyses have not been done for αMSH and NPY, or for any peptide *in vivo*, so we set out to use our imaging system to characterize the spatial impact and subcellular compartmentalization of NPY and αMSH signaling.

We considered three potential modes of peptide transmission. First, in the discrete model, somas and associated neurites function as independent peptide sensing compartments^91–95^, and show largely uncorrelated cAMP signaling (Figure 4G; although most neurite regions-of-interest (ROIs) connected to a soma are likely dendrites, we termed these neurites to be conservative). Second, in the large-volume transmission model^56,87,88^, neuropeptide signals are broadcasted widely across somas and neurites, so cAMP signals should be correlated within the same cell and across different cells (Figure 4G). Third, in the cAMP diffusion model, neuropeptide transmission is locally restricted but the resulting cAMP changes are seen in other compartments of the same cell due to intracellular diffusion (Figure 4G). In this case, trial-by-trial cAMP signals in somas and associated neurites should be correlated.

We used the above logic to re-analyze the cAMP-(Figures 2-3) and NPY-imaging datasets (Figures 4A-4F). Initial support for the discrete model comes from single-trial analyses of a PVH^MC4R^ neuron soma and two connected neurites, all three of which show local cAMP increments but on separate trials (Figure S4G). To systematically characterize cAMP signals in soma-neurite pairs of ROIs, we trained Cellpose 2.0^96^ to automatically segment neurite ROIs, followed by manual matching of neurites with associated somas (Figure S4H and Methods). Analyses of AgRP-to-PVH^MC4R^ cAMP imaging data indicated stochastic cAMP decrements in neurites similar to those observed in somas (Figures 3E and 4H). Further, neurite cAMP decrements were more persistent in the fasted state, similar to soma decrements (Figures 3F-3G and 4I). We next evaluated the presence or absence of concurrence between soma hits and neurite hits. The neurite cAMP signals averaged across *soma* hit trials were weak and similar to signals averaged across *soma* miss trials, indicating a lack of strong correlation between a cAMP decrement in a neuron’s soma and in its neurite (Figure 4J). Likewise, neurite analysis of POMC-to-PVH^MC4R^ cAMP imaging data also showed stochastic cAMP increments that were more persistent in the fed state (Figures 4K-4L). Changes in neurite cAMP were weak and of similar amplitude when averaged across *soma* hit trials or miss trials, again indicating decorrelation of soma and neurite signals (Figure 4M). These results show that NPY and αMSH transmission to PVH^MC4R^ neurites largely follow the same rules as soma signaling, albeit with a lower hit probability per surface area in neurites (Figure S4H). However, the stochastic hits and misses are decorrelated between somas and neurites of the same cell, therefore favoring the discrete compartmental model over large-volume transmission or cAMP diffusion models.

The above correlation metric treats all soma-neurite pairs equally regardless of distance, which may obscure correlations between close-range soma-neurite pairs. Such close-range correlations may occur if a neuropeptide release event has a spatially restricted impact on adjacent but not more distant targets (Figure S4I). We used XNOR (Figure S4J) as a similarity metric that measures the coincidence of simultaneous hits or misses in soma-neurite pairs. We found that for both AgRP-to-PVH^MC4R^ and POMC-to-PVH^MC4R^ transmission, concurrent cAMP changes in soma-neurite pairs were much more common for pairs less than ∼100 μm apart (Figure S4K; similar findings were also observed using other similarity metrics, Figures S4L-S4M). A similar falloff of correlated cAMP signals with distance was also seen for recordings from somas of two different cells and for NPY signals measured at different locations (Figure 4N), demonstrating that spatially adjacent somas can be impacted by the same plume of peptide release. These data further support the claim that spatially localized peptide release—but not intracellular diffusion of cAMP—defines the spatial scale of correlations in cAMP. Taken together, these results support a discretized mode of neuropeptide signaling in which stochastic neuropeptide release events evoke local cAMP changes in compartments within and across PVH^MC4R^ neurons (Figure 4O).

### Competition for cAMP signaling by opposing hunger and satiety peptides

In Figures 2-3, POMC axon stimulation drove larger and more persistent cAMP transients in the fed state, while AgRP axon stimulation drove larger and more persistent cAMP transients in the fasted state. Since the magnitude and persistence of cAMP increments and decrements are enhanced during different hunger states, it is unlikely that a global state such as arousal dictates the overall magnitude and persistence of peptidergic signaling (Figure 5A). We tested whether these hunger-state-dependent differences in signaling could be due to changes in i) axon excitability, ii) cAMP clearance, and/or iii) cAMP production across fasted and fed states. To test the first hypothesis, we asked whether photostimulation-evoked calcium transients in AgRP or POMC axons differed across hunger states. Photometry measurements of Axon-GCaMP6s^97^ *in vivo* showed that our stimulation protocol triggers short-lived calcium transients with similar amplitudes and decay kinetics in fasted and fed mice (Figures S5A-S5B), ruling out the axon-excitability hypothesis. To test the second hypothesis, we compared the clearance kinetics of cAMP produced by the blue-light-activated adenylyl cyclase, biPAC^55,98,99^. In brain slices from fasted and fed mice, cAMP clearance in PVH^MC4R^ neurons occurred within several minutes and did not differ between fasted and fed states (Figure S5C), ruling out the cAMP-clearance hypothesis.

We then tested the hypothesis that cAMP production in PVH^MC4R^ neurons was state dependent. AgRP neurons are known to exhibit elevated ongoing activity in the fasted state^8–10,100–102^, which likely results in a tonic level of NPY release that could hinder αMSH-induced cAMP production (Figure 5A). Likewise, in the fed state, steady-state levels of αMSH due to elevated tonic firing of POMC neurons^8,9^ could activate MC4R receptors to counter the suppression of cAMP production by NPY (Figure 5A).

To test this peptide competition hypothesis, we sought to artificially mimic this receptor competition by introducing neuropeptide agonists that are known to induce artificial states of hunger or satiety in behavioral experiments^42,80,81,103,104^ (Figure 5A). First, we tested whether mimicking the tonic elevation in αMSH that likely occurs in fed mice is sufficient to attenuate, in fasted mice, the amplitude and persistence of AgRP stimulation-induced cAMP decrements. As expected, intraperitoneal (i.p.) pre-injection of melanotan II (MTII, 3 mg/kg), a satiety-promoting MC4R/MC3R agonist (Figure 5B), elevated cAMP levels in PVH^MC4R^ neurons in fasted mice (Figure S5D). Although this manipulation did not affect the hit rate of AgRP stimulation-induced cAMP decrements (Figures S5E), it reduced cAMP decrement amplitude by 46% and persistence by 41% (Figure 5C) in a manner similar to our observations in fed mice (Figure 3G). To more directly simulate the effects of persistently elevated αMSH, we replaced the i.p. MTII delivery with i.c.v. pre-infusion of 1 nmol of αMSH^103^ (2 μl), a level of αMSH that was sufficient to elevate cAMP in PVH^MC4R^ neurons (Figures S2E-S2G). αMSH delivery caused an even greater reduction in the amplitude (78%) and persistence (86%) of AgRP stimulation-evoked cAMP decrements in the fasted state (Figures 5D-5E and S5F-S5G). We also tested if NPY delivery^80,81,104^ (0.5 nmol in 1 μl, i.c.v.) would weaken POMC stimulation-evoked cAMP increments in fed mice (Figure 5F). Consistent with the peptide competition hypothesis, i.c.v. pre-infusion of NPY in fed mice (Figures S3E-S3F) reduced cAMP increment amplitude by 54% and persistence by 83% (Figures 5G and S5H-S5I) in a manner similar to that observed in fasted mice (Figure 2I). While an MC4R antagonist also reduced cAMP increment amplitude, it did not attenuate persistence (Figure 2N), demonstrating that merely blocking POMC-evoked signaling at MC4Rs without introducing downstream competition at the level of cAMP is not sufficient to fully mimic state-dependent modulation.

The above experiments demonstrate that elevated levels of hunger or satiety peptides mutually blunt each other’s cAMP signaling magnitude and persistence. According to this competition model, brief elevation of a peptide due to prior axon activation should also locally enhance cAMP signaling upon additional release of the same peptide. To test this idea, we identified rare cases in which the same PVH^MC4R^ neuron exhibited cAMP transients on two consecutive trials (likely due to two peptide release events in the same spatial vicinity; Figure S5J). As predicted, cAMP transients were strengthened on the second hit as compared to the first during both AgRP and POMC stimulations (Figures S5J-S5O). Likewise, concentrating the stimulations of peptide release in time by shortening the inter-stimulation interval (from 52 s to 2 s) triggered cAMP increments and decrements that lasted more than 10 min (Figures S5P-S5Q). Together, these results show that neuropeptide competition is dose-dependent.

### Stochastic, persistent cAMP signaling during feeding defines the rate of satiation

The above findings regarding the peptide competition hypothesis indicate that cAMP levels in PVH^MC4R^ neurons are most effectively determined by the *relative* firing rates of AgRP axons as compared to POMC axons. Such a system should be able to filter out^105^ co-excitation or co-inhibition of AgRP and POMC neurons^7,9^, and be most effective in modifying cAMP levels when the activity of AgRP and POMC neurons concurrently shifts in opposite directions (Figure 6A). Such antiphasic modulation of AgRP and POMC neuron activity has been observed for virtually all regulators of energy balance (leptin^3,106^, ghrelin^8,107^, feeding^8,9^, exercise^108^). Accordingly, i.p. injection of the hunger hormone ghrelin, which stimulates AgRP neurons and inhibits POMC neurons^8,107^, readily produced a strong and sustained cAMP decrement in PVH^MC4R^ neurons (Figures S6A-S6B).

Feeding rapidly suppresses the firing of AgRP neurons and increases the firing of POMC neurons^8,9^. For fasted mice that have learned that a cue will predict a food reward, the bulk of these changes in neural activity occurs within seconds of food cue presentation and onset of food consumption^8–11^. Consequently, early in a feeding bout, the rise of αMSH release from POMC neurons should trigger cAMP increments in PVH^MC4R^ neurons, while the decrease in AgRP activity should withdraw competition and render the POMC-evoked cAMP increments persistent (Figure 6A). To experimentally characterize cAMP dynamics during natural feeding and test whether stochastic cAMP signaling also occurs in this context, we designed a feeding task in which drops of milkshake (Ensure; 15 μl/trial) were delivered to a head-fixed mouse with the same 60-s ITI as the above optogenetic experiments. After training, mice rapidly consume each milkshake delivery, albeit with higher lick rates in the fasted state than in the fed state^11^ (Figure 6B). 2p-FLIM measurements of absolute cAMP levels show that, in both fasted and fed states, the average cAMP level of PVH^MC4R^ neurons gradually rises in the first ∼4 trials and plateaus for the remainder of the session while mice are still feeding (Figures 6C and S6C). Even prior to food consumption, cAMP levels in the fed state were higher than in the fasted state (Figure 6C), consistent with elevated baseline extracellular levels of satiety peptides. These results show that the majority of elevations in cAMP occur early during feeding, when the opposing changes in AgRP and POMC neuron activity are largest^8–11^. We therefore divided each session into a climbing epoch (Trials 1-4 of the session) and a steady-state epoch (Trials 5-15) when analyzing single-trial cAMP transients (Figures 6D-6E). We found that feeding-evoked cAMP transients are also stochastic (Figures 6D-6G), indicating that stochastic neuropeptide signaling is not merely an artifact of optogenetic stimulation. As predicted by the drop in AgRP activity that coincides with the increase in POMC activity during feeding (Figure 6A; likely resulting in decreased competition from NPY), single-trial feeding-related cAMP increments were persistent (Figures 6F-6G and S6D-S6E). After the first four trials, both the rate and amplitude of hits stabilized at lower levels (Figures 6D-6G), resulting in the observed plateauing of intracellular cAMP concentration (Figure 6C).

Is the *simultaneous* drop in AgRP activity and rise in POMC activity important for feeding regulation? To answer this question, we bred POMC-Dre;AgRP-Cre mice^46^ and independently stimulated POMC neurons (with Dre-dependent ChR2) and inhibited AgRP neurons (with Cre-dependent hM4Di^21,109^) (Figure 6H). Either manipulation alone reduced feeding (see also^22,46^), but the combination of the two resulted in greater feeding suppression than either alone (Figure 6H). Together with the cAMP imaging data above, these results argue that peptide competition at the level of cAMP production functions as a biochemical filter that favors anti-phasic changes in hunger and satiety peptide levels.

### Elevating cAMP in PVH^MC4R^ gradually accelerates satiation

The above results strongly suggest that elevating cAMP in PVH^MC4R^ neurons should be sufficient to promote satiety. As an initial test, we used biPAC to bypass neuropeptide signaling and directly stimulate cAMP production in PVH^MC4R^ neurons. We found that a tonic, low level of biPAC activation reduced feeding in a refeeding paradigm involving *ad libitum* access to milkshake (Figure 6I). To understand the timescale of the satiety-promoting effect, we designed a task in which mice lick during an audible cue (“Tone”) to obtain milkshake (“Reward”) (Figure 6J). The purpose of the cue is to synchronize consumption and allow for trial-locked optogenetic stimulation of cAMP production in some trials. We used one of two types of sessions on alternating days, with each session containing 50 trials at 1 trial/min. In control sessions, no biPAC stimulation was delivered. In experimental sessions, biPAC stimulation was delivered at random on 50% of trials and consisted of 1 s of stimulation starting 5 s before cue onset (Figure 6K). This randomized biPAC stimulation design allows for within-session behavioral analysis, and the 5-s delay between stimulation and cue onset allows for cAMP to reach peak levels during cue and reward delivery (see Figure S5C). An example session is shown in Figure S6F. Over the course of 50 trials, lick rates during cue and reward windows and success rates declined faster in the experimental sessions involving biPAC stimulation (Figures 6K and S6G-S6H). The lick rates and success rates did not recover after an additional 10 min with no biPAC stimulation, indicating satiety rather than transient disengagement. Furthermore, within an experimental session, we could not detect any differences in behavioral responses between biPAC-stimulation trials and interleaved no-stimulation trials (Figures 6L and S6I), showing that elevated cAMP drives a gradual enhancement of satiety across many minutes. The drops in tone- and reward-evoked lick rates over a given ten-trial period scaled with the number of biPAC stimulations during this period (Figures 6M and S6J), arguing that repeated elevations in cAMP have an accumulating, dose-dependent effect that gradually accelerates satiation.

### cAMP sensitizes PVH^MC4R^ neurons to feeding-related excitatory inputs

How cAMP in PVH^MC4R^ neurons regulates feeding ultimately depends on how this second messenger alters the activity of these satiety-promoting neurons. A previous study found that prolonged incubation with αMSH in brain slices strengthens excitatory inputs to PVH^MC4R^ neurons^110^, but did not determine the involvement of cAMP or the dynamics of the synaptic plasticity *in vivo* during feeding.

To address these questions, we used fiber photometry to record calcium transients in PVH^MC4R^ neurons with the sensor RCaMP1a^111^ during the same conditioned feeding task as above, in which fasted mice lick during an audible cue to obtain milkshake reward at one trial/min (see Figures 6J). In contrast to the presence of cAMP responses in early trials of a session (Figure S6D-S6E), calcium responses to food consumption were absent in the first five trials, consistent with previous reports^67,112^ (Figure S7A). However, calcium responses gradually developed over the next ten trials and became clearly visible after 50 trials (Figures 7A top, 7B, and S7A). Calcium responses showed a delayed increase that peaked ∼4 s after licking onset (Figures 7A top and S7A). PVH^MC4R^ neurons were also more active at the end than at the beginning of the session in moments when no tone or reward was delivered^67^ (Figure S7B). These gradual increases in food consumption-evoked and ongoing activity in PVH^MC4R^ neurons are consistent with the key role of PVH^MC4R^ neuron activity in promoting satiety^13,28,41–^ 44.

**Figure 7.**
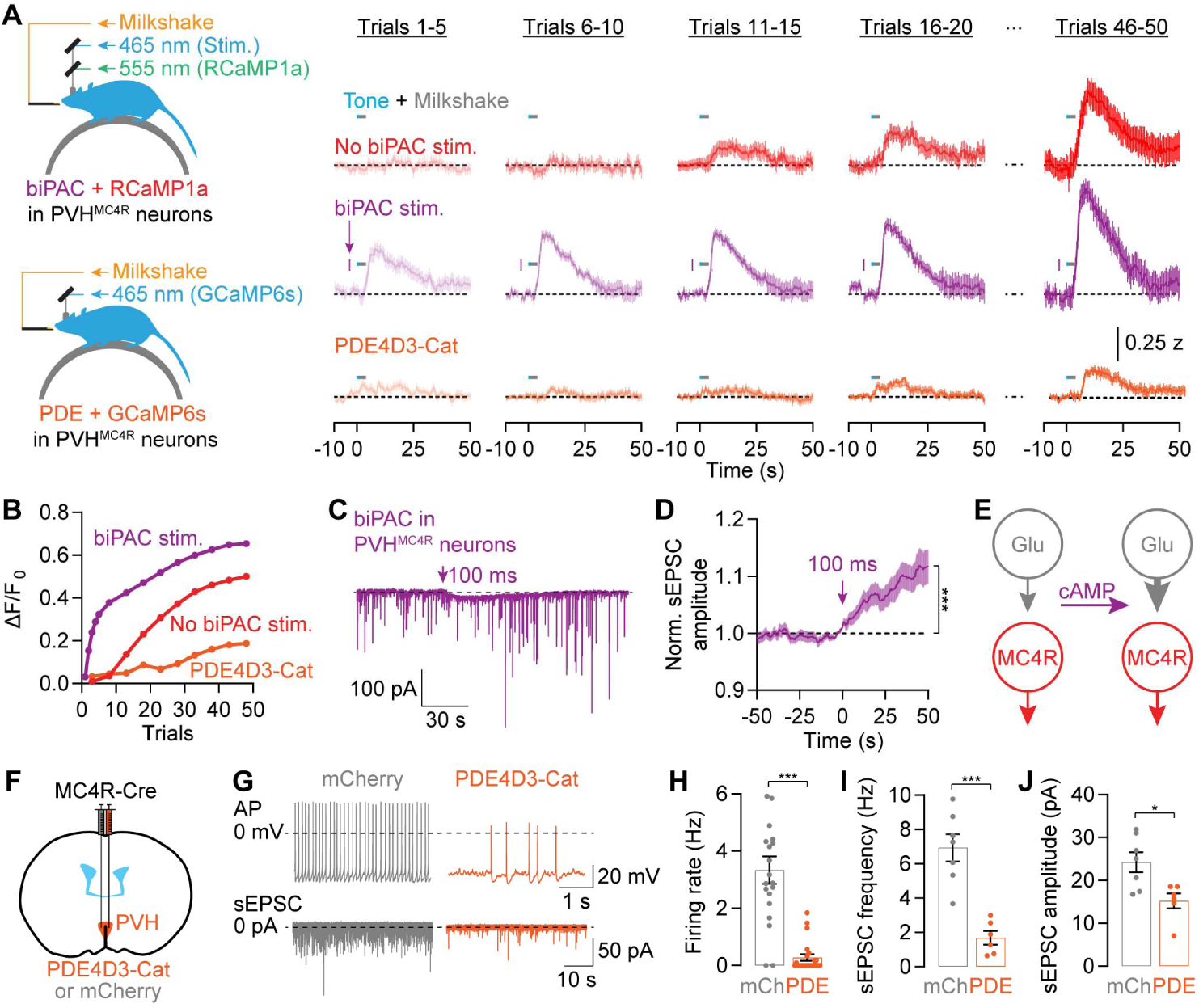
cAMP potentiates feeding-related excitatory input to PVH^MC4R^ neurons. **(A-B)** In a feeding assay where fasted mice lick during a cue to obtain a small bolus of milkshake at 1 trial/min, GCaMP6s signals in PVH^MC4R^ neurons demonstrate gradual growth of feeding-related responses (red), which likely promote satiety. When biPAC is used to produce cAMP 5 s before each trial, calcium responses grow faster and to a greater amplitude (purple). Artifacts due to bleed-through of photo-stimulation light were blanked. Reduced and delayed growth in calcium responses is seen when PDE4D3-Cat is co-expressed to blunt natural and artificial increases in cAMP in PVH^MC4R^ neurons (orange). In B, the first five trials of the biPAC stimulation group (purple) are plotted individually to capture the rapid growth in response magnitude in the presence of additional biPAC stimulation prior to each cue presentation. Peak calculations are from 5-8 s after cue onset. n = 8 mice per condition. **(C-E)** Brief 100-ms biPAC stimulation (example in C) potentiates amplitude (D) of spontaneous excitatory inputs, thereby sensitizing PVH^MC4R^ neurons to excitatory inputs (E). n = 16 cells from 3 fed mice. **(F-J)** cAMP degradation by PDE4D3-Cat (F) reduces the firing rate (G-H) and spontaneous EPSC frequency (I) as well as EPSC amplitude (J) in PVH^MC4R^ neurons in brain slices from fed mice (H-J: unpaired t-test; H: n = 19-23 cells from 4 mice each, I-J: 6-7 cells from 4 mice). mCh: mCherry control.

The gradual development of feeding responses in PVH^MC4R^ neurons across minutes (Figures 7A-7B) roughly matched the time scale of biPAC-induced decreases in feeding (Figure 6M). This led us to hypothesize that cAMP regulates the rate at which the calcium response to feeding develops. To test this, in a separate session we artificially drove cAMP production in these same PVH^MC4R^ neurons via brief biPAC photostimulation 5 s before each cue. Although the shape of each PVH^MC4R^ response to food consumption did not change, responses emerged earlier in the session, and were detectable in the first 5 trials – a time when the control traces were flat (Figures 7A top-middle and 7B). These feeding-related calcium responses during biPAC-stimulation sessions continued to grow across trials, ultimately exceeding the amplitude of control traces (Figures 7A top-middle and 7B). In the absence of feeding, biPAC stimulation did not excite PVH^MC4R^ neurons *in vivo*^98,113^ (Figures 7A and S7C-S7F). The accelerated emergence of excitation is likely caused by cAMP-mediated synaptic plasticity that strengthens excitatory inputs^114–116^, as whole-cell recordings in brain slices show that cAMP production by biPAC activation triggers a small but consistent potentiation of spontaneous EPSC (sEPSC) amplitudes (Figures 7C-7E; see Figures S7G-S7I for additional slice data).

When we blocked cAMP signaling with PDE4D3-Cat in PVH^MC4R^ neurons, feeding-related calcium responses emerged substantially later within the session and exhibited lower amplitudes even in the final trials (Figures 7A-7B). The delayed and diminished excitation is likely due to weakened synaptic inputs. Whole-cell recordings in brain slices show that PDE4D3-Cat expression decreased the spontaneous firing rate of PVH^MC4R^ neurons as well as the frequency and amplitude of excitatory inputs (Figures 7F-7J; see Figures S7J-S7M for additional slice data). These results show that PDE4D3-Cat expression causes hyperphagia and obesity (Figure 1) by diminishing the excitation of PVH^MC4R^ neurons^117^.

Taken together, these results demonstrate that the feeding-evoked and ongoing activation of the PVH^MC4R^ satiety neurons emerges gradually over the course of a meal, via potentiation of excitatory synaptic inputs at a rate that is set by the current level of cAMP (see Discussion and [^110,118^]). This cAMP level is, in turn, determined in large part by the tonic and phasic extracellular levels of competing satiety and hunger peptides, αMSH and NPY.

## Discussion

We have used imaging and manipulation of cAMP signaling to investigate spatiotemporal neuropeptide dynamics and downstream biochemical computations. These computations gradually culminate in changes in PVH^MC4R^ neuron spiking activity that then become detectable using conventional calcium imaging and electrophysiology (Figure 8). We show that photostimulation of the neuropeptide-releasing axons of AgRP and POMC neurons leads to stochastic neuropeptide release (Figures 2-4). This stochasticity, together with restricted peptide diffusion, enables a discretized mode of signaling, in which stochastic cAMP transients are detected in subcellular compartments of downstream PVH^MC4R^ neurons (Figure 4). In addition to having opposite effects on cAMP production, hunger and satiety peptides also inhibit each other’s signaling efficacy (Figures 5 and 8), rendering NPY signaling dominant in the fasted state and αMSH signaling dominant in the fed state. Feeding resolves this competition between peptides by simultaneously increasing αMSH release and decreasing NPY release, resulting in a gradual accumulation of cAMP in PVH^MC4R^ neurons (Figure 6). Finally, cAMP gradually sensitizes PVH^MC4R^ neurons to feeding-related excitatory inputs, thereby calibrating the gradual activation of these satiety-promoting neurons to the accumulation of a sufficient amount of food (Figures 6-7).

**Figure 8.**
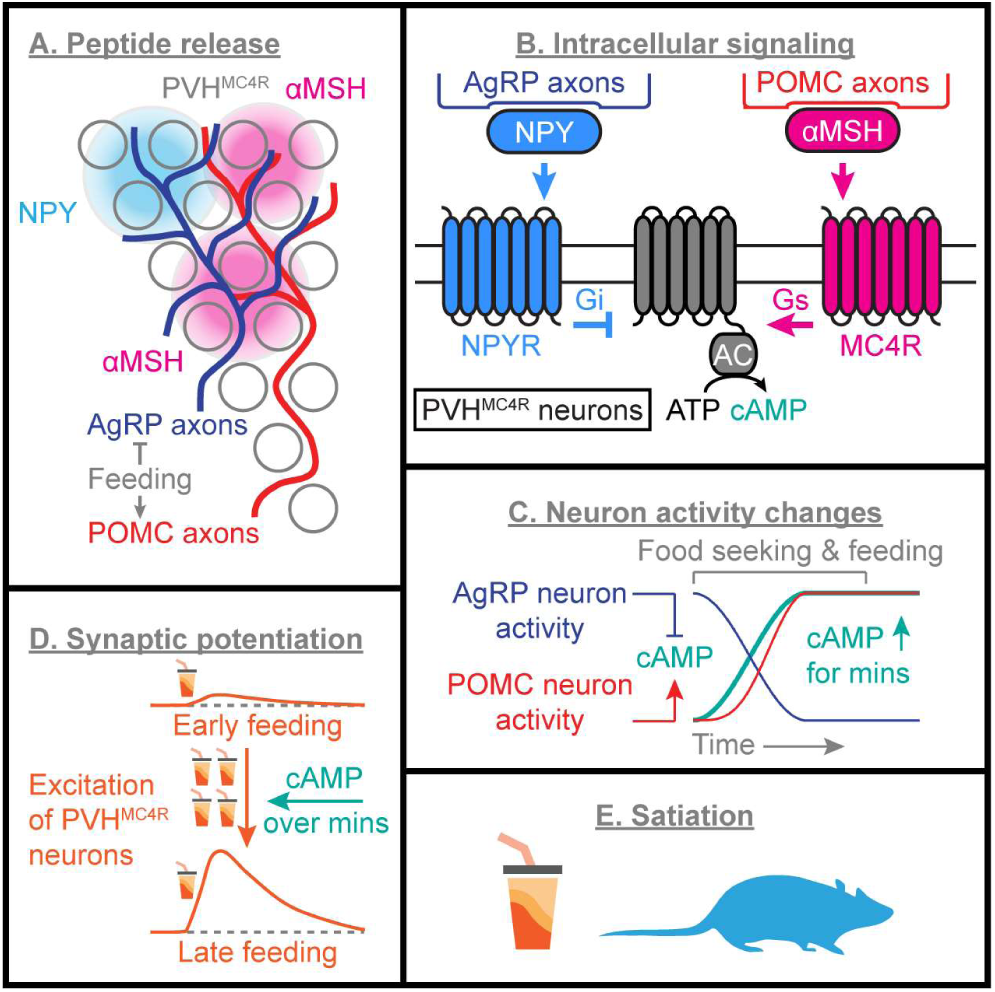
Summary of findings. Spatially localized plumes of neuropeptides NPY and αMSH are released stochastically in the PVH (A), where they compete to control cAMP in PVH^MC4R^ neurons (B). Feeding reduces AgRP neuron activity while elevating POMC neuron activity, a combination that results in sustained cAMP elevation in PVH^MC4R^ neurons for tens of minutes (C). In turn, elevated cAMP gradually potentiates feeding-related excitatory inputs to satiety-promoting PVH^MC4R^ neurons (D), thereby promoting satiation (E).

### Necessity and implications of stochastic neuropeptide signaling in biochemical computations

We show that neuropeptide release events are intrinsically stochastic in *in vivo* and brain slice experiments (Figures 2-4). Photostimulating AgRP axons triggers bimodally distributed NPY release (Figures S4B-S4E) and cAMP decrements in PVH^MC4R^ neurons (Figure 3D). The amount of NPY release (i.e., size of npyLight transients) is also insensitive to stimulation duration (Figures 4E-4F). Photostimulating POMC axons also triggers bimodally distributed cAMP increments (Figure 2D). These results lead us to propose that each cAMP event reflects the exocytosis of a single dense-core vesicle. Such stochasticity may be a consequence of the small number of dense-core vesicles in AgRP and POMC boutons^73^, similar to what is observed for most other synapses in the brain^56,119^ (but see [^120^]). Neuropeptides are replenished through axonal transport of dense-core vesicles at a rate of ∼1 μm/s^121^ – faster than typical axonal transport but perhaps still insufficient to compensate for frequent dense-core vesicle release (which typically occurs within a second^90^). These predictions can be tested using novel sensors of dense-core vesicle trafficking and fusion119,122.

In addition to preventing neuropeptide depletion, the sparsity of αMSH and NPY release, together with their intrinsically persistent impact on cAMP, enables a high-resolution mode of information accumulation during repeated food consumption events and associated changes in AgRP and POMC neuron activity (Figure 6). During feeding, the rapid changes in firing of AgRP and POMC neurons likely reflect the amount of additional calories that will be gained by imminent feeding behaviors^8,13,110^. The value of gradual cAMP accumulation is that it can integrate this estimated calorie-intake information across each bout of food consumption to properly calibrate the strengthening of excitatory inputs to PVH^MC4R^ neurons and the associated rate of satiation with calorie intake. This surprisingly slow mode of information integration via minutes-long biochemical signaling – even following a single taste of food – may underlie the effectiveness of slower meal consumption in earlier meal termination and reduced weight gain^123–125^. Due to this slow and integrative nature of biochemical signaling, it will be important to consider the activity of AgRP and POMC neurons over time scales of minutes, in addition to their moment-to-moment fluctuations (e.g., see [^126^]), when assessing their roles in energy balance.

### Peptide competition enables accurate representation of energy balance

We hypothesize that the degree of elevation in cAMP determines the *rate* of satiation. If we compare satiation rate to the speed of a vehicle, αMSH release from POMC axons and NPY release from AgRP axons may be analogous to gas and brake pedals, respectively. Feeding simultaneously steps on the gas pedal (i.e., increases POMC activity and αMSH release) and releases the brake (i.e., inhibits AgRP activity and NPY release). An elevation in cAMP does not immediately drive a switch to a sated state, but instead increases the speed of approach towards satiety. This analogy also illustrates that the *difference* between the activity of POMC and AgRP neurons is particularly important for understanding when and how these inputs to PVH^MC4R^ neurons will modify cAMP signaling. As such, concurrent increases or decreases in the activity of both AgRP and POMC neurons (e.g., during injection of cocaine, amphetamine, and nicotine^7^) may not predict changes in feeding. Indeed, our findings show that cAMP signaling in PVH^MC4R^ neurons is particularly sensitive to opposite-direction changes in extracellular αMSH and NPY, such as during feeding. Accordingly, MC4R agonists (e.g., setmelanotide^127^) may be more effective in promoting weight loss if used in combination with NPY antagonists or PDE inhibitors.

Hunger and satiety peptides inhibit each other’s signaling efficacy (Figure 5). At a molecular level, competition may take place through allosteric competition between the Gαs and Gαi binding sites on an adenylyl cyclase protein^128–131^. This mutual inhibition effectively mitigates the impact of fleeting changes in AgRP and POMC neuron activity (e.g., when a mouse finds an unexpectedly inaccessible food source^8^) on the progression towards satiety. Such friction may be well suited for maintaining a bi-stable hunger-satiety dichotomy and slows down transition from low to high cAMP during feeding, in addition to other mechanisms hypothesized to carry out similar functions in this circuit^100,101,132–134^.

We inferred the presence of baseline levels of peptides from the experiments using agonists (Figures 5B-5G), the repeated-hits analyses (Figures S5J-S5O), the grouped-stimulation experiments (Figures S5P-5Q), the state-dependent differences in basal cAMP (Figure 6C), and from prior *in vivo* experiments^8–11^. However, direct evidence for differences in the baseline levels and identity of various neuropeptides across fasted and fed states remain elusive. Given the slow diffusion of neuropeptides in the brain (∼10-20 s to cross 100 μm^58^) and the presence of peptidases^135^, a substantial duration of tonic or experimentally evoked activity (e.g., 98-s stimulation sequences in Figures S5P-S5Q) in peptide-releasing neurons may be required for substantial extracellular accumulation of peptide. Shorter bursts of activity (2-16 s) may not be able to achieve the same level of peptide accumulation (see Figure 4E). Future development of fluorescence lifetime neuropeptide sensors and other approaches^136^ will be instrumental in defining the state-dependent neuropeptidergic context surrounding each neuron, and how it affects signaling efficacy of any given neuropeptide.

### Resolving the logic of fast and slow transmission from neuropeptide-releasing neurons

Consistent with the idea that NPY transmission is responsible for the persistent hunger-promoting effects observed tens of minutes after termination of sustained AgRP stimulation^20,137^, we show that a grouped series of AgRP axon stimulations over 98 s results in a persistent decrease in cAMP that outlasts the stimulation by many minutes (Figures S5P-S5Q). This may be due to an accumulation of high levels of extracellular NPY that exceeds rates of clearance and breakdown. AgRP axons also release GABA onto PVH^MC4R^ neurons^44^, which should rapidly suppress activity in PVH^MC4R^ neurons. Unlike NPY, GABA-mediated inhibition tightly tracks the dynamics of AgRP stimulation^44^. Therefore, fast and slow transmission use biophysical and biochemical modes of signaling, respectively, to impact downstream circuitry at different timescales^13,21,138^. In contrast to AgRP neurons, stimulation of POMC neurons rarely results in fast synaptic transmission to PVH^MC4R^ neurons^110^, and instead modulates PVH^MC4R^ neuron activity gradually across tens of minutes (Figure 7). This difference likely explains why acute stimulation or inhibition of POMC neurons does not have potent effects on feeding at short timescales of a few minutes^23,110,139^. Our study also could not rule out a role for other signaling mechanisms downstream of MC4R that act in parallel with cAMP^41,47,54,62^.

WThe peptide competition model predicts that reducing AgRP neuron activity in the fasted state, when αMSH levels are low, is insufficient on its own to elevate cAMP levels in PVH^MC4R^ neurons. Early in the feeding assay, when AgRP neurons should be inhibited, we also did not observe feeding-related calcium responses in PVH^MC4R^ neurons that might be expected due to removal of GABAergic inhibition or via potential closing of GIRK channels upon removal of NPY^44,140,141^. This lack of effective disinhibition may reflect a lack of fast excitatory transmission onto PVH^MC4R^ neurons in fasted mice. These findings could explain why mimicking the suppression of AgRP neurons alone causes only a moderate suppression of feeding (Figure 6H and Krashes et al.^22^) as compared to the combined inhibition of AgRP neurons and excitation of POMC neurons that mimics natural activity patterns during feeding^8,9^ and that produces the largest satiety effects (Figure 6H; recently, the Brüning lab independently arrived at the same conclusion, personal communication).

### The 100-μm spatial impact of neuropeptide signaling

While it is proposed that most MC4Rs are found in the primary cilium near the PVH^MC4R^ soma^142–144^, POMC synapses are most commonly found on the distal dendrites of PVH neurons^73^. The ∼100 μm impact diameter during stochastic release of feeding-related neuropeptides (Figure 4N) may resolve this discrepancy due to the spread of peptidergic signals throughout much of the somatodendritic span of a given neuron. We note that this spatial scale is consistent with recent studies of neuropeptide diffusion^58,90^, suggesting that local diffusion may be a general mode of peptide action. We do not know whether NPY and αMSH release primarily occur from the synapse-forming terminals of AgRP and POMC neurons or from the axonal boutons with no postsynaptic partners^73^, nor did we record from distal dendrites of PVH^MC4R^ neurons. However, cAMP signaling downstream of POMC axon stimulation appears independent and compartmentalized in PVH^MC4R^ somas and (proximal) neurites. Moreover, cAMP signaling follows the same principles of stochasticity and competition-driven state dependency in each compartment (Figures 4 and S4). Thus, it is unlikely that the MC4R-dependent activation of cAMP via POMC axon stimulation acts solely on ciliary MCR4s. It remains to be determined whether somatic and/or dendritic cAMP signals in PVH^MC4R^ neurons drive compartment-specific potentiation of excitatory inputs (Figure 7). The same logic and challenges should also apply to other neural systems such as fly and worm brains, where neuropeptide-rich boutons often have no apparent synaptic partners^145,146^. In summary, our findings describe the biochemical mechanisms through which neuropeptidergic signals are integrated in space and in time to gradually calibrate neuronal plasticity and guide a measured transition from hunger to satiety.

## Acknowledgements

We thank Drs. J. Resch, D. Atasoy, Y. Livneh, H. Fenselau, J. Brüning, O. Yizhar, M. Frank, M. Crickmore, D. Rogulja, B. Sabatini, A. Banks, L. Tsai, J. S. Alvarado, H. Kucukdereli, M. Porniece, R. Essner, C. Massengill, and members of the Andermann and Lowell labs for useful feedback. We thank P. Sunkavalli, S. Sankar, J. Chen, P. Prasad, D. Guarino, H. Choh, and A. Sambangi for helping with animal care, behavioral experiments, and histology. C. Massengill helped test RCaMP1a. Drs. H. Fenselau and J. Brüning provided POMC-Dre mice and the unpublished Dre-dependent ChR2 AAV. Boston Children’s Hospital Viral Core (NIH P30 EY012196) and HMS Research Instrumentation Core provided viral and laser printing services, respectively. M. Cortopassi and Dr. A. Banks at the BIDMC Energy Balance Core performed the energy-balance and body-composition experiments as well as helped with analyses. Authors were supported by a Lefler Fellowship, a Charles A. King Trust Fellowship, and NIH K99 DK134853 (S.X.Z.); NIH T32 DK007529 (A.K.); NIH F32 DK112589 and T32N S007484 (A.L.); NIH T32 GM007753, T32 GM144273, T32 HL007901, and F30 DK131642 (P.N.K.); NIH U01 NS115579, and U19 NS123719 (L.T.); NIH R01 DK096010, R01 DK075632, R01 DK089044, R01 DK122976, P30 DK046200, P30 DK057521 (B.B.L.); NIH DP2 DK105570, R01 DK109930, DP1 AT010971, R01 MH12343, a McKnight Scholar Award, and grants from the Boston Nutrition and Obesity Research Center (P30 DK046200), the Klarman Family Foundation, the Pew Innovation Fund, and a Charles Robert Broderick III Phytocannabinoid Research Grant (M.L.A.).

## Author contributions

S.X.Z. and M.L.A. conceived the project and wrote the manuscript with methodological and editing inputs from all authors. S.X.Z., A.K., and A.L. conducted all surgeries. A.K. generated and maintained multi-transgene mice. A.K. performed the free-moving feeding experiments and related measurements. S.X.Z., L.F.C., and P.Z. performed photometry experiments. S.X.Z., A.K., and P.Z. performed *in vivo* two-photon imaging experiments, with technical help with i.c.v. infusion from P.N.K.. S.X.Z. and A.L. performed slice imaging experiments. J.C.M. performed electrophysiology experiments. A.K., L.F.C., and P.Z. performed all histology. Y.J., A.P., and L.T. developed npyLight. S.X.Z., A.K., J.C.M., A.L., B.B.L., and M.L.A. analyzed all data.

## Experimental Procedures

### Animals

All animal care and experimental procedures were approved by the Institutional Animal Care and Use Committee at Beth Israel Deaconess Medical Center (BIDMC). Animals were housed in a 12-hour-light/12-hour-darkness environment with standard mouse chow and water provided *ad libitum*, unless specified otherwise. Male and female mice older than 8 weeks were used in experiments, and the numbers of males and females were balanced to the degree allowed by each litter (see Table S1). All effects were seen in males and females, but male mice were typically larger in size and ate more. We used the following genotypes: B6.FVB-Tg(Pomc-cre)1Lowl/J (POMC-Cre^147^, 010714, The Jackson Laboratory), POMC-Dre^46^, Agrp^tm1(cre)Lowl^/J (AgRP-IRES-Cre^148^, 012899, The Jackson Laboratory), Mc4r^tm3.1(cre)Lowl^/J (MC4R-2A-Cre^44^, 030759, The Jackson Laboratory), and their F1 progeny. We did not intentionally backcross mice to isogenize the backgrounds, but most mice were in the C57BL/6J background. Mice with implants (e.g., fiber, GRIN lens) were single-housed to avoid damage, and all other mice were group housed. Sample sizes were chosen to reliably measure experimental parameters and keep with standards of the relevant fields^20,21,23,55,110,137,138,149^, while remaining in compliance with ethical guidelines to minimize the number of experimental animals. Experiments did not involve experimenter-blinding, but randomization was used to determine experimental order and group assignments.

### Pharmacological agents

NPY (Tocris, 1153, Batch 27A) was dissolved in ACSF (Tocris, 3525, Batch 81A) at 500 pmol/μl for i.c.v. experiments (500 pmol/mouse) and dissolved in water at 200 nM for slice experiments (200 nM). αMSH (Tocris, 2584, Batch 6A) was dissolved in ACSF at 0.5 nmol/μl and used for i.c.v. infusions (1 nmol/mouse). Melanotan II (i.e., MTII, MC3R/MC4R agonist, Tocris, 25661, Batch 6A) was dissolved in saline at 0.5 μg/μl and used for i.p. injections (3 mg/kg). SHU9119 (MC3R/MC4R antagonist, Tocris, 3420, Batch 6A) was dissolved in ACSF at 0.5 nmol/μl and used for i.c.v. infusions (1 nmol/mouse). BIBP 3226 trifluoroacetate (NPY1R antagonist, Tocris, 2707, Batch 4A/263489) was dissolved in ACSF with 10% DMSO at a concentration of 5 nmol/μl and filtered (Millipore, Millex-GV, 0.22 μm). BIBP 3226 was used in i.c.v. infusions (10 nmol/mouse). CGP 71683 hydrochloride (NPY5R antagonist, Tocris, 2199, Batch 3B/275453) was dissolved in ACSF with 30% DMSO at a concentration of 5 nmol/μl and filtered. CGP 71683 was used in i.c.v. infusions (10 nmol/mouse).

### Chronic food restriction

Chronic food restriction was performed as previously described^11,150^. In the fasted state, mice were kept at 85% of *ad libitum* weight (weighed every morning before experiments) by restricting food access during and after experiments. Weight and food intake were logged per mouse per day. To reduce the discomfort of weight changes, most mice only went through one food-restriction/refeeding cycle to test fasted and *ad libitum* fed states. For mice that were used for multiple experiments (e.g., testing different stimulation protocols), we also tried to minimize the confounding effects of time between fasted and fed states. To do so, we typically performed half of the *ad libitum* fed-state experiments first, then all the fasted-state experiments, and finally finished the ad lib fed-state experiments. We did not observe lasting effects of food restriction, with mice regaining full body weight within a day or two of *ad libitum* food access.

### Surgeries

Viral injections, fiber implantations, and GRIN lens implantations were generally performed as described in Zhang et al.^55^ with the following specifications. For experiments using 400-μm diameter fibers or 500-μm diameter GRIN lens, in order to ensure a snug fit for the fiber or lens, to reduce brain motion, and to accelerate recovery, we pre-set the insertion tracks by lowering needles with matching diameters (27 gauge or 25 gauge, respectively) to a depth of 0.1 mm above the final depth of the fiber or lens. For 22XX-gauge cannula implants, the track was set with a 22-gauge needle. In initial experiments, GRIN lens placements were ∼0.2 mm more lateral due to concerns about potential impact on animal health, and the coordinates became the values below (which enables better views of PVH) once initial health concerns were found to be unsubstantiated. All AAVs were injected at a titer of 3-15 10^13^ gc/ml (see below for volumes used). In experiments where unmixed cADDis and PDE4D3 viruses were used, both AAVs were pre-diluted 1:3 to a final titer of 3-4 10^13^ gc/ml before injections to minimize potential effects on cell heath (which we verified in *post hoc* histological analyses). In the minority of cases where viral expression was absent (<10% of surgeries), we excluded the data from subsequent analyses. All animals were allowed to recover for at least 3 weeks prior to onsets of experiments. No obvious capsid competition was seen in histology.

For experiments that test the feeding effects of expressing PDE4D3-Cat.^55^ in PVH^MC4R^ neurons, AAV2/1-EF1a-DIO-mKate2-PDE4D3-Cat (Boston Children’s Hospital Vector Core; Addgene 169128) or AAV9-Ef1a-DIO-mCherry (UNC Vector core) was injected bilaterally (50 nl) in the PVH (Bregma: AP -0.75 mm, ML ±0.25 mm, DV -4.8 mm) of MC4R-2A-Cre^44^ mice. For experiments involving two-photon imaging of cADDis^66^ in the PVH during optogenetic stimulation of POMC axons or AgRP axons in the head-fixed mice, rAAV1-hSyn-FLEX-Chrimson-tdTomato^68^ (150 nl; UNC Vector Core) was bilaterally injected into the arcuate nucleus of POMC-Cre^147^;MC4R-2A-Cre or AgRP-IRES-Cre^148^;MC4R-2A-Cre mice, respectively (Bregma: AP -1.5 mm, ML ±0.3 mm, DV -5.9 to -5.75 mm). Viral spillover to PVH was rare (see Figures S2D and S3D), and the few cells that were labeled by Chrimson-tdTomato are excluded in analysis. AAV2/1-hSyn-DIO-GreenDownwardcADDis (Boston Children’s Hospital Vector Core) was injected unilaterally in the PVH (150 nl; Bregma: AP -0.75, ML 0.2 mm, DV -4.8 mm). A larger volume of the cADDis was injected to minimize the chance of missing MC4R neurons, and any spillover to arcuate neurons should not interfere with the interpretation of results since cADDis is a sensor, not an actuator. In early experiments, we used POMC-Dre^46^;MC4R-2A-Cre mice (instead of POMC-Cre;MC4R-2A-Cre), together with an AAV expressing Cre in a Dre-dependent manner (AAV-FREX-Cre^151^), to enable Chrimson expression, but no difference was seen in the cADDis response size to stimulating POMC neurons in POMC-Dre mice or POMC-Cre mice, so the datasets were combined. A doublet GRIN lens (NEM-050-25-10-860-DM, GRINTECH; 0.5 mm diameter; 9.89 mm length; 250 μm focal distance on brain side at 860 nm [NA 0.5]; 100 μm focal distance on the air side [NA 0.19]; 2.6x magnification) was implanted in the PVH, ipsilateral to the site of expression of cADDis (Bregma: AP -0.75, ML 0.2 mm, DV -4.7 mm). We chose doublet lenses here because the 2.6x magnification provided more pixels per cell and per neurite during imaging. For imaging experiments where i.c.v. infusion was performed, we also inserted a 22XX gauge cannula (MicroGroup; 0.6 mm ID, 0.72 mm OD, 6.99 mm length) in the posterior lateral ventricle (Bregma: AP -1 mm, ML 1.8 mm, DV -2.2 mm). This coordinate was chosen to maximize the distance between cannula and GRIN lens, so as to avoid potential damage to the lens during tubing insertion and removal. A titanium head plate was centered over the GRIN lens and fixed to the skull using Metabond. A 3D-printed acrylic funnel was cemented onto the headplate to enable light-shielding during experiments. The area surrounding the lens was covered with Metabond and then dark dental cement to reduce autofluorescence. The GRIN lens was protected by a cut-off tip of an Eppendorf tube (Fisher) that was secured using Kwik-Cast (WPI).

The experiments that involve two-photon fluorescence lifetime imaging of cADDis in the PVH during feeding used the same mice as those for optogenetic stimulation of POMC and AgRP axons.

For experiments that involve two-photon imaging of npyLight in PVH during optogenetic stimulation of AgRP axons in slice, AAV1-ihSyn-tTA/TREnpyLight1.0 (150 nl; Tian lab, manuscript in preparation) was injected bilaterally in the PVH of AgRP-IRES-Cre mice (Bregma: AP -0.75 mm, ML 0.25 mm, DV -4.8 mm). AAV1-hSyn-FLEX-Chrimson-tdTomato (150 nl; UNC Vector Core) was injected bilaterally into the arcuate nucleus (Bregma: AP -1.5 mm, ML ±0.3 mm, DV -5.9 to -5.75 mm).

For experiments that test the excitability of AgRP and POMC axons, AAV1-hSyn-FLEX-Chrimson-tdTomato (UNC Vector Core) and AAV5-hSynapsin1-FLEx-axon-GCaMP6s^97^ (Addgene 112010-AAV5) were injected bilaterally in the arcuate nucleus of AgRP-IRES-Cre and POMC-Cre mice, respectively (1:1 mixture, 150 nl total; Bregma: AP -1.5, ML 0.3 mm, DV -5.9 to -5.75 mm). An optic fiber (400 μm diameter core, multimode, 6.0 mm length, NA 0.48, Doric) was implanted unilaterally in the PVH (Bregma: AP -0.75, ML 0.1 mm, DV -4.7 mm). A titanium head plate was fixed to the skull using Metabond to enable head-fixation.

For experiments that measure the clearance rate of cAMP in the PVH^MC4R^ neurons, AAV2/1-DIO-EF1α-mKate2-biPAC^55^ (Boston Children’s Hospital Vector Core; Addgene 169127) and AAV2/1-hSyn-DIO-GreenDownwardcADDis (Boston Children’s Hospital Vector Core) were bilaterally injected in the PVH of MC4R-2A-Cre mice (1:1 mixture, 150 nl total; Bregma: AP -0.75 mm, ML 0.25 mm, DV -4.8 mm).

For experiments that measure the clearance rate of cAMP in the PVH^MC4R^ neurons that also express PDE4D3-Cat, AAV2/1-DIO-EF1α-mKate2-biPAC (Boston Children’s Hospital Vector Core; Addgene 169127), AAV2/1-EF1a-DIO-mKate2-PDE4D3-Cat (Boston Children’s Hospital Vector Core; Addgene 169128), and AAV2/1-hSyn-DIO-GreenDownwardcADDis (Boston Children’s Hospital Vector Core) were bilaterally injected in the PVH of MC4R-2A-Cre mice (1:1:1 mixture, 150 nl total; Bregma: AP -0.75 mm, ML 0.25 mm, DV -4.8 mm).

For experiments that measure the impact of cAMP production in PVH^MC4R^ neurons on feeding, AAV2/1-DIO-EF1α-mKate2-biPAC (Boston Children’s Hospital Vector Core; Addgene 169127) was unilaterally injected in the PVH of MC4R-2A-Cre mice (150 nl; Bregma: AP -0.75 mm, ML 0.2 mm, DV -4.8 mm). An optic fiber (400 μm diameter core, multimode, 6.0 mm length, NA 0.48, Doric) was implanted unilaterally in the PVH (Bregma: AP -0.75, ML 0.1 mm, DV -4.7 mm). A titanium head plate was fixed to the skull using Metabond to enable head-fixation.

For experiments that measure the impact of simultaneous POMC neuron activation and AgRP neuron inhibition on feeding, AAV8-hSyn-DIO-hM4D(Gi)-mCherry^22^ (Addgene 44362-AAV8) and AAV8-FREX-ChR2-EYFP (gift from Henning Fenselau) were injected bilaterally in the arcuate nucleus of AgRP-IRES-Cre;POMC-Dre mice (1:1 mixture, 150 nl total; Bregma: AP -1.5 mm, ML ±0.3 mm, DV -5.9 to -5.75 mm). An optic fiber (400 μm diameter core, multimode, 6.0 mm length, NA 0.39, RWD) was implanted unilaterally in the PVH (Bregma: AP -0.75, ML 0.1 mm, DV -4.7 mm).

For electrophysiological recording of PVH^MC4R^ neurons expressing PDE4D3, AAV2/1-EF1a-DIO-mKate2-PDE4D3-Cat (Boston Children’s Hospital Vector Core; Addgene 169128) or AAV9-Ef1a-DIO-mCherry (UNC Vector core) was injected bilaterally (50 nl) into the PVH of MC4R-2A-Cre mice (Bregma: AP -0.75, ML ±0.25 mm, DV -4.8 mm).

For electrophysiological recording of PVH^MC4R^ neurons expressing biPAC, AAV2/1-DIO-EF1α-mKate2-biPAC (Boston Children’s Hospital Vector Core; Addgene 169127) was injected bilaterally into the PVH of MC4R-2A-Cre mice (150 nl; Bregma: AP -0.75, ML ±0.25 mm, DV -4.8 mm).

For experiments that measure the acute effect of cAMP production on neuron activity, AAV2/1-DIO-EF1α-mKate2-biPAC (Boston Children’s Hospital Vector Core; Addgene 169127), together with either AAV1-Syn-Flex-GCaMP6s-WPRE-SV40^152^ (Addgene 100845-AAV1) or AAV1-Syn-Flex-NES-jRCaMP1a-WPRE-SV40^111^ (Addgene 100848-AAV1), was bilaterally injected into the PVH of MC4R-2A-Cre mice (1:1 mixture, 150 nl total; Bregma: AP -0.75, ML ±0.25 mm, DV -4.8 mm). For the photometry version of this experiment, an optic fiber (400 μm diameter core, multimode, 6.0 mm length, NA 0.48, Doric) was implanted unilaterally in the PVH (Bregma: AP -0.75, ML 0.1 mm, DV -4.7 mm). A titanium head plate was fixed to the skull using Metabond to enable head-fixation.

For experiments that measure the effect of cAMP production on PVH^MC4R^ neuron activity during feeding, AAV2/1-DIO-EF1α-mVenus-biPAC (Boston Children’s Hospital Vector Core) and AAV1-Syn-Flex-NES-jRCaMP1a-WPRE-SV40 (Addgene 100848-AAV1) were unilaterally injected into the PVH of MC4R-2A-Cre mice (1:1 mixture, 150 nl total; Bregma: AP -0.75, ML 0.2 mm, DV -4.8 mm). An optic fiber (400 μm diameter core, multimode, 6.0 mm length, NA 0.48, Doric) was implanted unilaterally in the PVH (Bregma: AP -0.75, ML 0.1 mm, DV -4.7 mm). A titanium head plate was fixed to the skull using Metabond to enable head-fixation.

For experiments that measure the effect of cAMP degradation on PVH^MC4R^ neuron activity during feeding, AAV2/1-EF1a-DIO-mKate2-PDE4D3-Cat (Boston Children’s Hospital Vector Core; Addgene 169128) and AAV1-Syn-Flex-GCaMP6s-WPRE-SV40 (Addgene 100845-AAV1) were unilaterally injected into the PVH of MC4R-2A-Cre mice (1:1 mixture, 150 nl total; Bregma: AP -0.75, ML 0.2 mm, DV -4.8 mm). An optic fiber (400 μm diameter core, multimode, 6.0 mm length, NA 0.48, Doric) was implanted unilaterally in the PVH (Bregma: AP -0.75, ML 0.1 mm, DV -4.7 mm). A titanium head plate was fixed to the skull using Metabond to enable head-fixation.

### Freely-moving 24-hr feeding, body-length, and body-weight measurements

For freely-moving 24-hr feeding (Figure 1E), body length (Figure S1C), and body weight measurements (Figure 1C), animals were singly housed, and body weight and 24-hr food intake of individual animals were measured every week following AAV injection. Body weight was measured on day 1 of each week. 24-hr food intake was measured for 3 consecutive days at the beginning of each week (days 1-3) and these values were averaged to account for daily variations in the food intake.

For body length measurements, animals were maintained in group housing with their original littermates throughout the duration of the experiment. Body length (nose-to-anus length) was measured every week following AAV injection during brief isoflurane anesthesia.

### Indirect calorimetry and MRI

24-hour food intake (LabDiet 5008, 3.56 kCal/g), water intake, physical activity (beam breaks), oxygen consumption, carbon dioxide production, and body mass of single-housed mice were measured every 2 minutes using the Sable Systems Promethion indirect calorimeter in the BIDMC Energy Balance Core. Mice were weighed and body compositions were scanned using an EchoMRI 3-in-1 body composition analyzer (no anesthesia) before they were placed in the Promethion system for recording. All mice were allowed 12 hours of habituation in the system before recording the 72-hr experiment. From the measurements, energy intake, energy expenditure, respiratory exchange ratio, physical activity, and energy balance were calculated using CalRCalR^65^ (https://calrapp.org/).

### Head-fixation with food delivery

Head-fixation was used in photometry and two-photon microscopy experiments in order to record cAMP and calcium activity while reducing motion artifacts. The experiments were carried out using head-fixed mice that could run freely on a circular treadmill equipped with IR beam breaks to record speed^150^. For experiments involving food delivery, milkshake (Ensure Plus, 350 kCal per 8 oz bottle) was delivered through a lickspout^150^. The speed of food delivery was controlled by gravity and gated by solenoid pulses. Each pulse opened the solenoid for 150 ms, which resulted in 3 μl of milkshake delivery. For non-binge feeding experiments, we delivered 5 Ensure pulses (15 μl; e.g., Figure 6B) or 10 Ensure pulses (30 μl; e.g., Figure 6K) per trial. For all head-fixed feeding experiments, there was a 4-min waiting period before the first trial to record baseline sensor activity and bleaching rate. All dataset were collected as triplicates per condition.

Food delivery during two-photon imaging was controlled with an Arduino (https://github.com/xzhang03/Train_generator). One session was performed per day. For two-photon imaging experiments in Figures 6B-6G, 10 trials were conducted per session at a rate of 1 trial per minute, and each trial used a 5-pulse train (300 ms between pulse onsets).

Food delivery during photometry or behavioral experiments was controlled using the Nanosec photometry-behavioral system (https://github.com/xzhang03/NidaqGUI). In head-fixed binging assay, each lick triggers a pulse of Ensure, and we recorded the number of licks over an hour per mouse (Figure 6I). Low-level biPAC stimulation was performed by pulsing 465 light (50 μW) at a pulse-width of 6 ms and a frequency of 50 Hz, with a resulting power of 15 μW.

For photometry and head-fixed food intake experiments with trial structures (e.g., Figure 6J and 7A), 50 trials were conducted per session at a rate of 1 trial per minute and each trial used a 10-pulse train of Ensure deliveries (300 ms between Ensure pulse onsets, 3 μl/pulse, 30 μl total per trial). The increased number of trials and increased total Ensure delivery per trial during the photometry experiments were designed to partially satiate mice over the course of the experiment. For experiments using conditional food delivery (e.g., Figure 6J), a 2 kHz tone was played for 1 second, during which mice must lick at least once to trigger the food delivery train. If triggered, the food delivery train started as soon as the tone was over. Otherwise, mice must wait until the next trial (1 trial/min), and the current trial was omitted in photometry analysis. At 10 pulses per train and 50 trains per session, the maximum amount of milkshake a mouse could consume there is 1.5 ml, which is roughly 50% of the absolute maximum volume that a mouse would consume in previous experiments^11^. For experiments involving biPAC stimulation (e.g., Figure 6J), a 1-s light pulse (465 nm, 1 mW peak, 70% duty cycle) was delivered 5 seconds before the tone in 50% of the trials (randomly selected, in Figure 6J). The 5-s delay was chosen to allow sufficient cAMP production before the tone (see Figure S5C). To verify satiation, we also sometimes (at least once per mouse per condition) gave the mouse the choice to perform an additional 10 trials 10 minutes after the regular experiments (see Figure 6K).

Mice were trained for ∼2 weeks before each experiment, with the following sequence: 1) habituation to handling, 2) habituation to head-fixation, 3) habituation to food delivery via lick spout, and 4) operant conditioning assay (if applicable).

### Head-fixed photometry

Head-fixed photometry experiments were conducted as described previously^55^ using mice running on a circular treadmill. For experiments using a green sensor (e.g., Axon-GCaMP6s), excitation light from a 465-nm LED (for fluorescent sensor excitation, ∼100 μW peak) and a 630-nm LED (for optogenetic stimulation of Chrimson; 1 mW peak if used) were combined in a four-port fluorescence mini-cube (FMC4_E(460-490)_F(500-550)_O(580-650)_S, Doric) and transmitted to the implanted fiber via a patch cord (1 m length, NA 0.57, Doric). For biPAC stimulation during GCaMP6s recording (Figure S7D), biPAC stimulation was performed by transiently elevating the intensity of the 465-nm photometry LED light from ∼100 μW to 1 mW. This light intensity should be sufficient to photoactivate biPAC, whose half-saturation light intensity is ∼30 μW/mm^2^ (see characterization in Zhang *et al*. ^55^). For experiment using a red sensor (e.g., RCaMP1a), excitation light from a 555-nm LED (for sensor excitation, ∼100 μW peak) and a 465-nm LED (for optogenetic stimulation of biPAC; 1 mW peak if used) were combined in a five-port fluorescence mini-cube (FMC5_E(450-490)_F1(500-540)_E2(550-580)_F2(600-680), Doric) and transmitted to the implanted fiber via a patch cord (1 m length, NA 0.57, Doric). The emitted light was measured from the emission port of the mini-cube using a femtowatt photoreceiver (2151, Newport).

All photometry and optogenetic excitation lights were modulated as interleaved pulses controlled by the Nanosec photometry-behavioral system that was initially developed in a previous study^55^ (https://github.com/xzhang03/NidaqGUI). Each cycle (20 ms per cycle, or 50 cycles per second) consists of 1) turning on the photometry (465 nm or 555 nm) LED for 6 ms (Step 1) and then 2) turning off the photometry LED for 14 ms (Step 2). If Chrimson stimulation was used, once every 5 cycles, the 630-nm LED was turned on immediately after Step 2 and then turned off 10 ms later, which corresponds to 10 Hz, 10 ms pulses (duty cycle = 10%). If biPAC stimulation was used, once every cycle, the 465-nm LED was turned on immediately after Step 2 and turned off 14 ms later, which corresponds to 50 Hz, 14 ms pulses (duty cycle = 70%). The recording was conducted in darkness. Black heat shrink was placed around the fiber to help prevent the LED light from being seen by the mice. For two weeks before the photometry experiments, each mouse was habituated to handling and head-fixation.

### Two-photon imaging

Two-photon imaging was performed using a two-photon resonant-galvo scanning microscope (NeuroLabWare) controlled by Scanbox (https://scanbox.org/) as described previously^55,150^. An InSight X3 laser (Spectra-Physics) was used to excite the fluorophores (910-1050 nm), and the emission light was filtered (green: 510/84 nm; red: 607/70 nm; Semrock) before collection with photomultiplier tubes (H10770B-40; Hamamatsu). The XY scanning was performed using resonant/galvo mirrors and the Z scanning was achieved with an electrically tunable lens (Optotune).

Two-photon Fluorescence-lifetime Imaging Microscopy (FLIM) was enabled by modifying the existing two-photon microscope (NeuroLabWare). Both FLIM and the conventional two-photon imaging used the same excitation light path. In experiments involving FLIM, an extra beam-splitter (75T/25R, Semrock) was added to the emission light path such that 75% of the emission light was directed to the standard intensity-based PMT (H10770B-40; Hamamatsu) and the rest (25%) was split to the FLIM light path. In the FLIM light path, the emission light was filtered (510/84 nm; Semrock) before entering the hybrid PMT (Becker and Hickl). PMT data were digitized using a time-correlated single-photon counting board (Becker and Hickl) to estimate photon counts and arrival times. Frames were constructed from the pixel and frame clocks in the NeuroLabWare microscope. Laser clocks were recorded with an independent photodiode (Becker and Hickl) in the excitation path. In the FLIM experiments, the laser intensity was controlled (<10 mW) so that the FLIM PMT received 2.5-5 × 10^5^ photons per second (to avoid cross-pulse photon contamination, see below). Each frame was collected from 2-5 seconds of continuous scanning so that it contained at least 1 million photons/mm^2^ to ensure accurate lifetime estimates while minimizing laser power.

### Two-photon imaging of fixed brain slices

In order to count the number of PDE4D3-Cat- and mCherry-expressing cells in an automated fashion^96^, we used the two-photon microscope to scan fixed brain slices that were stained with mCherry (594 nm excitation for secondary antibody). Brain slices (60 μm thick) were prepared, and every other slice was mounted and then covered with a coverslip. The slices that contained the brain region of interest (i.e., PVH) were placed under a 16x water-immersion objective (NA 0.8, CFI75 LWD, Nikon), and were scanned at 15.5 frames/second and 796 × 512 pixels/frame. We excited the fluorophores at 1050 nm (5-15 mW). We collected 50 volumes (30 steps spanning 60 μm per volume).

### Two-photon imaging of acute brain slices

Acute brain slices were prepared as described in Lutas *et al*.^149^ and transferred to a recording chamber perfused with ACSF (oxygenated with 95% O_2_ and 5% CO_2_; flow rate: 2-5 mL/min) at room temperature. Two-photon imaging was performed using the same microscope and 16x lens as described above. The excitation wavelength used was 910 nm.

In pharmacological experiments involving the NPY sensor npyLight, each slice was imaged for 15 minutes (15.5 frames per second) and 200 nM NPY (Tocris, 1153) was applied via perfusion starting at 4 minutes into the recording. In npyLight experiments that involve photostimulating AgRP axons in the PVH, each slice was imaged across four runs, with each run corresponding to an optogenetic stimulation duration (2 s, 4 s, 8 s, 16 s). The order of the runs (i.e., stimulation durations) was randomly selected between ascending order (2 s first and 16 s last) and descending order (16 s first and 2 s last) to average out potential history-dependent effects. In each run, at time points 2 min, 3 min, 4 min, 5 min, 6 min, a 655-nm LED (1 mW/mm2, Luxeon Star LEDs) driven by an Arduino-controlled driver (Luxeon Star LEDs) was used to deliver photostimulation trains (https://github.com/xzhang03/bPACstim). The first 2 min of each run was used as baseline. To protect the PMT during optogenetic stimulation, the photostimulation pulses were time-locked to the start of each frame (immediately after galvo flyback, resulting in 15.5 photostimulation pulses per second). During frames with optogenetic pulses, the PMT was blanked for 10 ms, and the LED was turned on for the first 8 ms (and leaving 2 ms for LED to turn off before un-blanking the PMT). Using this method, the stimulation frequency is the same as frame rate (15.5 Hz) and the duration was controlled by frame number (e.g., 2 s = 31 frames of stimulation).

In experiments involving biPAC stimulation, we use longer optogenetic pulses (e.g., 100 ms) that cannot be synchronized to a single frame. Therefore, to protect the PMT, we manually turned off PMT while biPAC stimulation was triggered.

### *In vivo* intracerebroventricular infusion

Because of the close horizontal proximity of the cannula and the GRIN lens, we chose a minimal setup to perform i.c.v. infusions to not hinder the field of view or risk damaging the exposed lens. I.c.v infusion was performed via a 22XX-gauge cannula (MicroGroup, 0.6 mm ID, 0.72 mm OD, 6.99 mm length), a 1-ml syringe, an automated syringe pump (Harvard Apparatus, 70-4504), and the required tubing. When not imaged, the cannulas were plugged with small piece of tubing that is secured with a silicone gel (Kwik-Cast). Before imaging, the plug was removed, and the infusion tubing was inserted to a pre-calculated depth that does not protrude from the bottom of the cannula. The tubing was then sealed and secured with silicone gel. We regularly observe a small amount of back-flowing CSF when the plug was removed, indicating unobstructed cannula. All parts were sterilized before inserted into the cannula. The rest of the imaging setup was performed in the same way as regular imaging experiments. During infusion, the rate was kept consistent at 1 μl/min and the volume varies between 1 and 4 μl as determined by the peptide dosage. We occasionally saw increased running when infusing 4 μl, but no agitation or vocalization was noted. For peptides that are dissolved in ACSF/DMSO mixtures, same-ratio mixtures were used for the corresponding control experiments. We did not observe a noticeable impact of DMSO on cAMP.

### *In vivo* two-photon imaging experiments

*In vivo* two-photon imaging of the PVH via a GRIN lens in head-fixed mice was generally performed as described^55,149^. The excitation wavelength was 910 nm. The experiments were carried out using head-fixed mice that could run freely on a circular treadmill^150^. Imaging was performed with a 4x 0.2 NA air objective (Nikon) in mice implanted with doublet GRIN lens (see above; 2.6 magnification, NA 0.19 on the air side). Light shield material was used to protect the lens and the objective from external light. Imaging fields-of-view were at a depth of 100–300 μm below the face of the GRIN lens. The laser and frame parameters were described above. Because cAMP changes relatively slowly^55,99^, we opted to simultaneously image 2-3 FOVs that were at least 60 μm apart (using Optotune) to increase cell sampling and FOV counts at the expense of frame rate (effective frame rate = 10.3 for a 3-FOV session and 15.5 for a 2-FOV session). The FOVs were chosen to avoid repeated imaging of the same cells.

For the optogenetic experiments, we imaged for 15 min per session. Excitation light from a 617-nm LED (ThorLabs; M617L3) was focused into the GRIN lens using the same 4x objective. LED power was measured to be 2 mW below the 4x objective, and based on prior experiments, ∼50 of the power passes through the GRIN lens. Photostimulation trains were delivered at time points from 2 min through 11 min into the recording (10 trains, 1 min between train onsets). To protect the PMT during optogenetic stimulation, the photostimulation pulses were time-locked to the start of each frame (31 fps). During frames with optogenetic pulses, the PMT was blanked for 10 ms, and the LED was turned on for the first 8 ms (leaving 2 ms for the LED to turn off before un-blanking the PMT). Using this method, the stimulation frequency is the same as frame rate (31 Hz) and the duration was controlled by frame number (i.e., 250 frames = ∼8 s). In grouped stimulation experiments (e.g., Figures S5P-S5Q), the time between train onsets was shortened from 1 min to 10 s while keeping all other stimulation parameters the same (including 8 s train lengths). This effectively resulted in a single prolonged train per session lasting 98 s in duration.

For optogenetic experiments that also used i.c.v. infusion of neuropeptide agonists or antagonists, the drugs were pre-infused through the cannula 10 min before imaging started. For optogenetic experiments that also used i.p. injections (e.g., MTII), the injection was performed 15 min before imaging.

For i.c.v. infusion experiments (e.g., Figure S2F) or i.p. injection experiments (e.g., Figure S5D) without optogenetic stimulation, we imaged for 15 min per session, and the drugs were introduced at 3 min into the recording. In these experiments, because cAMP changes are slow, FLIM was also used, and the FLIM data were acquired at 1 frame (5-sec acquisition) every 10 seconds throughout the session.

For feeding experiments, simultaneous FLIM and non-FLIM imaging datasets (20 min per session) were acquired as described above. FLIM data were used to determine the overall changes in absolute cAMP levels, while non-FLIM (i.e. intensity-based imaging) data were used to visualize single-trial cAMP transients. FLIM data were acquired at 1 frame (5-sec duration) every 10 seconds throughout the session. Non-FLIM imaging data were acquired at 15.5 fps and only 1 FOV was imaged per session. Uncued food deliveries started at 4 min, and 10 trials were presented (1 min between trial onsets) using the methods described above.

### Electrophysiology in acute slices

To prepare *ex vivo* brain slices, 6-10 week-old mice were deeply anesthetized with isoflurane before decapitation and removal of the entire brain. Brains were immediately submerged in ice-cold, carbogen-saturated (95% O_2_, 5% CO_2_) choline-based cutting solution consisting of (in mM): 92 choline chloride, 10 HEPES, 2.5 KCl, 1.25 NaH_2_PO_4_, 30 NaHCO_3_, 25 glucose, 10 MgSO_4_, 0.5 CaCl_2_, 2 thiourea, 5 sodium ascorbate, 3 sodium pyruvate, oxygenated with 95% O_2_/5% CO_2_, measured osmolarity 310 – 320 mOsm/L, pH= 7.4. Then, 275-300 μm-thick coronal sections were cut with a vibratome (Campden 7000smz-2) and incubated in oxygenated cutting solution at 34°C for 10 min. Next, slices were transferred to oxygenated ACSF (126 mM NaCl, 21.4 mM NaHCO_3_, 2.5 mM KCl, 1.2 mM NaH_2_PO_4_, 1.2 mM MgCl_2_, 2.4 mM CaCl_2_, 10 mM glucose) at 34°C for an additional 15 min. Slices were then kept at room temperature (20–24°C) for ≥45 min until use. A single slice was placed in the recording chamber where it was continuously superfused at a rate of 3–4 mL per min with oxygenated ACSF. Neurons were visualized with an upright microscope (SliceScope Pro 1000, Scientifica) equipped with infrared-differential interference contrast and fluorescence optics.

Whole-cell patch clamp recordings were performed with an intracellular solution containing (in mM): 128 potassium gluconate, 10 KCl, 10 HEPES, 1 EGTA, 1 MgCl_2_, 0.3 CaCl_2_, 5 Na_2_-ATP, 0.3 Na-GTP (pH 7.3). Action potential frequency was assessed for a period of 2 min immediately after breakthrough. Assessment of spontaneous EPSCs (sEPSCs) was performed in voltage-clamp mode with recordings lasting at least 3 min after establishment of a stable baseline.

Cell-attached recordings (seal resistance 20-50 MΩ) were made in voltage-clamp mode with the recording pipette filled with ACSF and holding current maintained at *V*_h_ = 0 mV. To photoactivate biPAC, a LED light source (470 nm) was used. The blue light was focused onto the back aperture of the microscope objective (40x), producing wide-field illumination around the recorded cell of 10-15 mW per mm^2^ as measured using an optical power meter (PM100D, Thorlabs). A programmable pulse stimulator, Master-8 (A.M.P.I.), and pClamp 10.2 software (Molecular Devices, Axon Instruments) controlled the photostimulation output.

All recordings were made using a Multiclamp 700B amplifier, and data were filtered at 2 kHz and digitized at 20 Hz. Access resistance (<30 MΩ) was continuously monitored by a voltage step and recordings were accepted for analysis if changes were <15%.

### Histology

Perfusion and histology were performed as described in Garfield *et al*.^153^. Brain slices (60 μm thick) were collected and one of every three consecutive slices was scanned. The primary antibodies used in this paper are: chicken anti-GFP (1:1000, Invitrogen; used for GFP and GCaMP6s) and rat anti-mCherry (1:1000, ThermoFisher; used for mCherry and tdTomato). The secondary antibodies are Donkey anti-chicken 488 (1:1000, Jackson) and Donkey anti-rat 594 (1:1000, Jackson).

## Analysis Procedures

All data analyses were performed using custom scripts in MATLAB (MathWorks), Python, ImageJ (NIH), SPCImage (Becker and Hickl), and Prism (GraphPad).

### Freely-moving and head-fixed feeding assays

For food intake assays involving optogenetic and chemogenetic manipulations (Figure 6H), mice were singly housed immediately following the surgery and acclimated to the handling and experimental setting for >1 hour/day for 5 consecutive days prior to the experiment. Food intake studies were performed at the onset of the dark-cycle (Zeitgeber time 12-15 when mice engage in normal dark-cycle feeding). Food intake was measured at 0, 1, 2, and 3 hr time points. On the day of the experiment, mice were attached to the patch cord (1 m length, NA 0.57, Doric Lenses) and injected with either saline or CNO (1 mg/kg) 30 min prior to the onset of the experiment. The light stimulation was delivered via LEDs (Plexbright) at a peak power of 7-9 mW at the fiber tip. Optogenetic trains (20Hz, 10ms pulses, 1 s on 3 s off) were generated using a custom-made TTL pulse generator. The light stimulation protocol began immediately before the onset of the first food measurement and continued for the 3 h duration of each food intake session.

For the head-fixed binging assay (Figure 6I), we simply plotted mean cumulative lick number per mouse. Analysis of conditional head-fixed feeding assays were based on the lick rates of the mice during the tone window (1 s after tone onset) and reward window (10 s after reward delivery onset), regardless of trial success. A trial was marked as a success if a mouse licked at least once during the tone window. See Figure S6F for ethogram of an example experiment. These parameters were averaged per mouse (i.e., average of 3 sessions per condition) and per condition (e.g., Figure 6K). If biPAC stimulation was used, means of biPAC-stimulation trials and no-stimulation trials in the experimental session were calculated separately before comparing them to the control session (e.g., Figure 6L). For correlation analysis (e.g., Figure 6M), we analyzed sequences of 10 consecutive trials (e.g., Trials 23-32), calculated the change in the feeding metric (e.g. lick rate during cue), and correlated that value to the number of stimulation trials within that set of 10 trials.

### Analyses of head-fixed photometry experiments

Head-fixed photometry data were analyzed as described previously^55^ (https://github.com/xzhang03/Photometry_analysis). Pulses that trigger the photometry LED (50 Hz, see above) were used to determine when the LED was on. For each pulse (6 ms), we took the median of the corresponding data points in the photodetector trace to quantify the photometry signal. This results in a 50 Hz trace which we then filtered with a 10 Hz low-pass filter.

To estimate bleaching, we fitted the pre-stimulation data (first 4 min of each experiment) with a mono-exponential function and subtracted the estimated values from each timepoint in the photometry trace across the entire recording. Since all photometry experiments have a trial structure, we used the baseline fluorescence (e.g., the 10 s pre-stimulation window, see e.g., Figure S5A) to estimate ΔF/F_0_. The ΔF/F traces were then pooled across days and z-scored per mouse. To show single-trial photometry traces, we triggered photometry traces with cue onsets with a 10-s pre-cue baseline and a 50-s post-cue recording (see Figure S7A for an example experiment). For Figure 7A, we grouped the calcium transients in bins of 5 trials to show the gradual progression of excitation.

### Pre-processing of two-photon imaging experiments

Image registration was performed in MATLAB using a two-step process: 1) a rigid xy-translation step that is repeated 3 times (https://github.com/xzhang03/Tiff_preprocess), and 2) a non-rigid registration step using the Demonsreg function (https://github.com/xzhang03/Demonsreg-Oneshot). For experiments that involve simultaneous imaging of multiple z-planes, each z-plane was registered independently. To extract cell bodies, we down-sampled images spatially by a factor of two. Then, we used Cellpose 2.0^96^ to segment soma ROIs. Repeating somas from multiple FOVs were excluded during analysis. For fixed-slide imaging, the number of ROIs was reported as the number of somas expressing the markers. We note that this number likely only reflects 50% of total cell counts, as we imaged every other slice.

To extract neurite ROIs, we retrained three built-in Cellpose models – cyto, nuclei, and the base network – to manually labeled neurites in three steps on an NVIDIA RTX3080 GPU: 1) 100 iterations of a 31-image set, 2) 500 additional iterations of the same set, and 3), 2000 iterations of a 119-image set. We eventually decided to use the nuclei-based neurite model for subsequent analyses. Training progress, segmentation results, and the models are all deposited to a public repository (https://github.com/xzhang03/cellpose_GRIN_dendrite_models). After segmenting neurites, neurite ROIs were then manually matched to somas of the same field of view for future analysis.

For cross-day experiments (e.g., Figures 6B-6G), soma-ROIs in the same FOV were also matched manually.

After segmentation, we calculated the fluorescence traces from soma and neurite ROIs, and subtracted from them the fluorescence traces of the surrounding neuropil rings. Neuropil rings were calculated by first dilating the ROI by 14 pixels and subtracting all ROIs from the dilated mask. If the resulting neuropil ring has fewer than 2500 pixels, the process is repeated with incrementing dilating size. Mean neuropil traces were subtracted from the mean ROI trace at 1:1 scaling. The underlying assumption for the no-scaling subtraction is that if a given pixel contains photons from both the intended ROI (e.g., soma) and the neuropil, the neuropil photon contribution should be similar at the soma and the surrounding area. Therefore, the subtraction was done to remove global influences from the estimation of neurons’ responses to stimulation. The traces were then triggered relative to the onset of optogenetic stimulation or food delivery, and the relative changes (ΔF/F_0_) were calculated. We used a 20-s pre-stimulation and 110-s post-stimulation window to display most cAMP traces (e.g., Figure 2I). The 110-s post-stimulation window was chosen to capture the entirety of non-persistent cAMP transients.

Because cADDis shows decreased fluorescence intensity when cAMP concentration increases, we plot cADDis intensity changes on a flipped Y-axis (-ΔF/F_0_) for presentation purposes. For estimates of persistence indicies, trial-average traces with more than one hit event were excluded to prevent sustaining of cAMP signaling due to repeated hits.

### Pre-processing two-photon FLIM data

Pre-processing of FLIM data was performed as in Zhang et al.^55^ and briefly described below. Images were first pre-processed using SPCImage (Becker and Hickl). We used the first-moment method of estimating lifetimes^55,76^. The advantage of this method is its high signal-to-noise ratio (due to the lack of fit uncertainty) and robustness when the photon number is low. Moreover, to increase the accuracy of lifetime estimates, the first moment of each pixel was calculated after local, spatial binning (bin size: 5×5 pixels, ∼25 μm^2^; the size of the spatial binning was small compared to the typical size of a cell [1500-2000 pixels]). Further, if the peak of the resulting decay histogram (256 bins in time, 48.86 fs per bin) had fewer than 5 photons, then that pixel was excluded from subsequent analyses. In this way, we obtained an image of photon counts and an image of estimated lifetimes at the resolution of the original frame (650 × 550 pixels for *in vivo* imaging through a GRIN lens and 900 × 550 pixels for slice imaging).

The remainder of the data processing for FLIM experiments was performed in MATLAB (https://github.com/xzhang03/SPC_analysis). Both intensity and lifetime images were downsampled by a factor of two. Image registration was performed in the same manner as the non-FLIM (intensity-based) datasets above, and the shifts were applied to lifetime images. Segmentation was performed using Cellpose 2.0^96^. Since a typical cell had a mean pixel intensity of 10-15 photons/pixel, its lifetime was estimated from ∼20000-26000 photons. Neuropil-ring fluorescence intensity and lifetime traces were calculated as described previously^55^ and subtracted from the soma data. Lifetimes are usually plotted as changes from baseline, with the exception of Figure 6C, where changes are relative to the fasted baseline.

Because cADDis traces show decreased fluorescence lifetime when cAMP concentration activity increases, we plot their lifetime changes on flipped Y-axis (-Δlifetime) for presentation purposes.

### Classifier of cAMP increments and decrements

We found that, for any given cell, neuropeptide-mediated cAMP increments and decrements were unpredictable on a trial-by-trial basis, even when the same photostimulation protocol was used across trials. Therefore, we used a classifier to apply to determine hit trials (increments or decrements) or miss trials (no change in cAMP). Previous studies showed slow cAMP dynamics across many different brain areas^55,95,98,99^, so we assumed that cAMP also operates on a slow timescale in the PVH and pre-smoothed the single-trial cADDis trials before classification. The classifier is based on the area under receiver operating characteristic curve (auROC) applied to ΔF/F_0_ data from the pre-stimulation baseline and the post-stimulation period. The unit of classification is a single trial of a single ROI. The pre-stimulation window is defined as the last 20 sec before stimulation or food delivery. The post-stimulation window is defined as a 30-sec window after stimulation or food delivery, and the window onset is different for cADDis and npyLight experiments. Longer windows were used for the grouped stimulation experiment (Figures S5P-S5Q). The classifier is two-tailed. An auROC value greater than 0.995 or less than 0.005 indicates that there is at most 1% chance that the pre-stimulation and the post-stimulation data points are drawn from the same distribution (i.e., the null hypothesis). Trials with auROC>0.995 or auROC<0.005 were treated as hits, and then further differentiated into increments (post-stim. cAMP > pre-stim. cAMP) or decrements (post-stim. cAMP < pre-stim. cAMP). Within each group, the trials were sorted by their agonist response. Consistent with previous biochemical characterizations^29,30,41,142^, POMC stimulation drove mostly cAMP increments (>12% of the trials) while AgRP stimulation drove mostly cAMP decrements (>20%). The opposite-direction hits (i.e., cAMP decrements during POMC stimulation and increments during AgRP stimulation) occur in less than 1% of the trials, lending credibility to the classification scheme. Following the classification, we show means of hits and misses. Whenever appropriate, we also show means across all trials as well (e.g., Figure 2H).

Due to the serendipitously large estimated cAMP changes in PVH^MC4R^ neurons when compared to previous experiments^55,99^ as well as the bimodal nature of the amplitude of these changes (e.g., Figure 2D), classifier outcomes were not sensitive to classifier thresholds in non-pharmacological experiments. For experiments involving the application of neuropeptide agonists and antagonists, however, the amplitudes of cAMP increments and decrements did shrink closer to 0, and the hit-amplitude distribution (usually Gaussian) could at times become truncated (e.g., Figure S3H). We suspect that, in these experiments, cAMP changes were closer to the detection threshold of cADDis (Kd = ∼20 μM)^66^. We nevertheless kept the classification parameters the same for consistency. In these experiments, adjusting the classification thresholds did not qualitatively affect the results.

To calculate log distribution of peak ΔF/F_0_ values (e.g., Figure 2D inset), ΔF/F_0_ values were first offset by the max (for decrement events) or the min values (for increment events). An additional small offset (0.001 or -0.001) was then added to remove zeros, which would otherwise give infinity in a log function. The natural log function was then applied to the resulting values, from which the distribution was generated. For hits-only distribution of ΔF/F_0_ and area-under-curve values in Figure 4F, no offsets or log functions were used to enable more accurate cross-condition comparisons.

### Dice model of hits and misses

Stochasticity analysis was inspired by a previous study of coin-toss-like behavioral choices in *Drosophila*^154^ and studies of quantal release^86^. The key to the dice-roll or coin-toss model is that the probability of obtaining an outcome (e.g., cAMP increment) is the same from trial to trial. One ramification of this assumption is that, given the unitary event probability (P_1_), the chance of seeing 2, 3,…, n outcomes in a row must follow the power function P_n_ = P_1n_. We used this mathematical function to construct dice plots such as in Figure 2F. One disadvantage of this plot is that the chance of seeing 3 or more events in a row is close to zero (while still following the power function), so we also constructed linearized dice plots using the function Ln(P_n_) = n x Ln(P_1_), where Ln is the natural logarithm function (and could be a log function of any base). The downside of this linearized model is that it amplifies noise in small probabilities.

### Correlation of cAMP and NPY changes between ROIs

Correlation of binary events (hits or misses) between different ROIs were generally calculated using the XNOR (exclusive not-or) function (see Figure S4J for truth table), because this function does not give invalid values to sequences that lack a single hit. The disadvantage is that the estimation of the chance level of correlation needs to be calculated on a case-by-case basis, which we did through bootstrapping. We also verified the main conclusions with an independent method, F1 score^155^ (also referred to as Dice correlation), as well as Pearson correlation of ΔF/F_0_ values.

For analyses that also involve distance between ROIs, we used inter-centroid distances and adjust for GRIN lens magnification (2.6x magnification for doublets) as needed for *in vivo* experiments. For slice experiments (npyLight), no GRIN lens was used so we did not apply the same magnification factor.

### Kymograph of soma and neurite hits

In Figure S4G, the presence of a bipolar cell morphology with two clear, continuous neurites affords us a rare chance to use a kymograph to show the decorrelation between soma-and neurite-hits. We label the neurites A and B and the soma S, and show single-pixel changes in fluorescence intensity. Due to the low signal-to-noise ratio of this method, we used additional low-pass filtering on the traces (0.2 Hz filter).

### Electrophysiological analysis

Cell-attached spike recordings and whole-cell sEPSC recordings were analyzed using MATLAB by finding local peaks with pre-determined thresholds on current, charge (time-domain integral of current), and widths (https://github.com/xzhang03/ephys_analysis). The results were spot-checked manually to ensure fidelity.

### Statistics

Two-tailed t-tests and ANOVA were performed using Prism. For ANOVA, multiple comparisons were done using the Sidak *posthoc* correction. For behavioral, metabolic, and photometry results, each mouse (typically averaged between 3 tests) was treated as an independent sample. For two-photon experiments, each FOV was treated as an independent sample (2-5 FOVs per mouse), as done previously^55^. For non-fractional data, error bars show standard error of the mean (S.E.M.). Fractional analysis of hits and misses are done in MATLAB and uses the total number of trials multiplied by the number of ROIs as the denominators. Hit rates were estimated through fitting using the linearized dice model (e.g., Figure 2F) For bar plots of hit rates, error bars show 95% confidence interval of 100,000 iterations of bootstrapping the data (with replacements) and fitting the dice model per iteration. Hypothesis testing between fractional data (e.g., between hit rates) was performed using bootstrapping with the null-hypothesis being data from different conditions were drawn from the same pool. For all tests, non-significance means that the null hypothesis cannot be rejected (i.e., p ≥ 0.05), and significance in each figure is indicated as follows: *p<0.05, **p<0.01, ***p<0.001.

**Figure S1.**
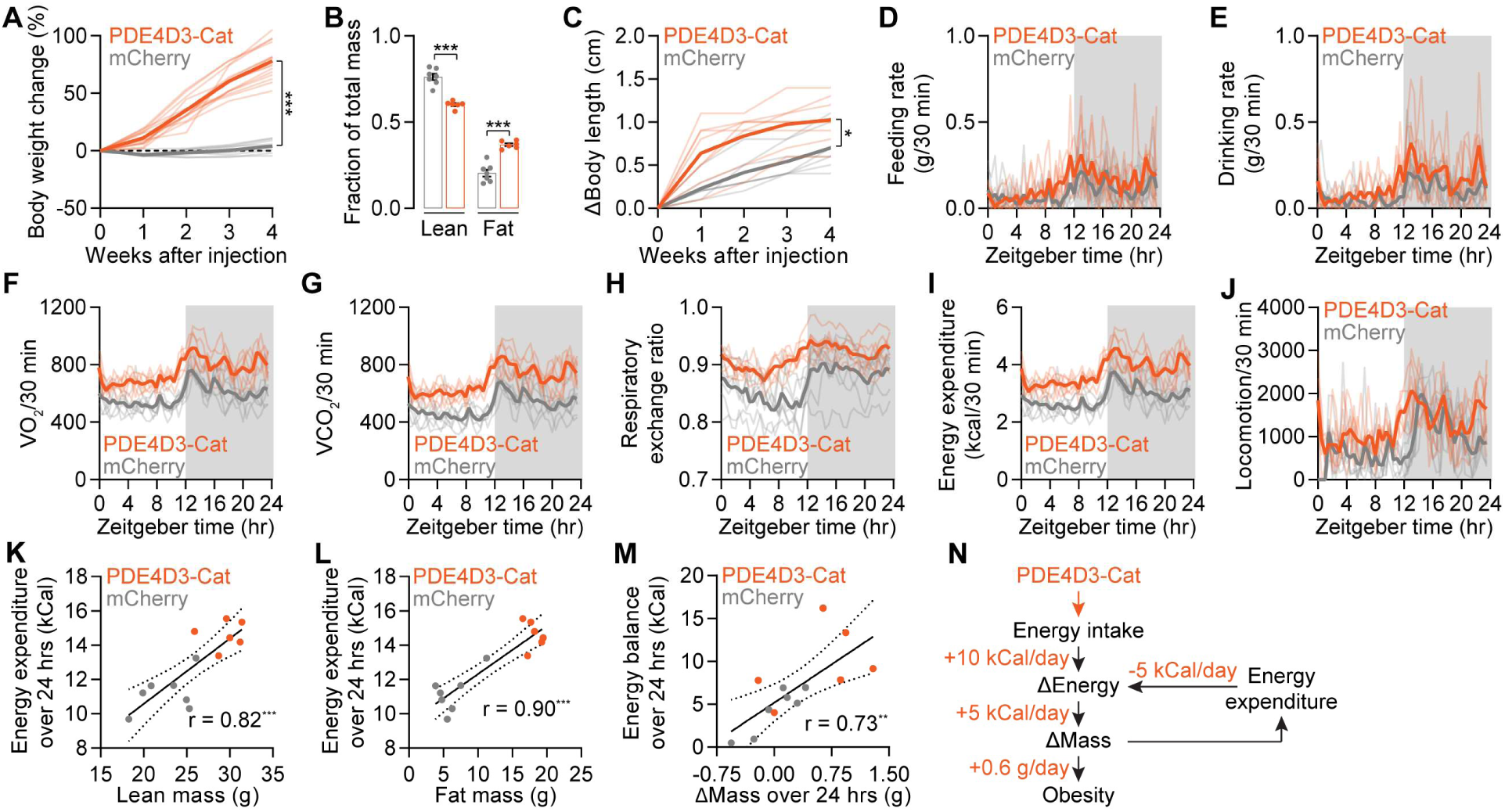
Metabolic changes in mice expressing PDE4D3-Cat in PVH^MC4R^ neurons. **(A)** AAV expression of PDE4D3-Cat in PVH^MC4R^ neurons in adult mice results in ∼80% weight gain in the 4 weeks following surgery (n = 9-15 mice, t-test). **(B)** Mice that express PDE4D3-Cat have an elevated contribution of fat mass to their body composition (n = 6-7 mice, One-Way ANOVA). **(C)** PDE4D3-Cat expression in PVH^MC4R^ neurons in adult mice results in ∼0.5 cm increase in axial length in the 4 weeks following surgery (n = 7-8 mice, t-test). **(D-J)** A panel of 24-hr recordings of metabolic parameters in 30-min bins: feeding rate (D), drinking rate (E), VO_2_ (F), VCO_2_ (G), respiratory exchange ratio (H), energy expenditure (I), locomotor activity (J). n = 6-7 mice. Gray shading: dark cycle. **(K-M)** Across individual mice, energy expenditure is well correlated with lean mass (K), fat mass (L), and change in mass (M), in both the presence or absence of PDE4D3-Cat expression. n = 6-7 mice. **(N)** Diagram of energy gain: PDE4D3-Cat expression results in increased energy intake (10 kCal/day) that is only partially offset by elevated energy expenditure (5 kCal/day), resulting in a net 5 kCal/day surplus which translates to 0.6 g/day of weight gain. Hence, mice become obese over time.

**Figure S2.**
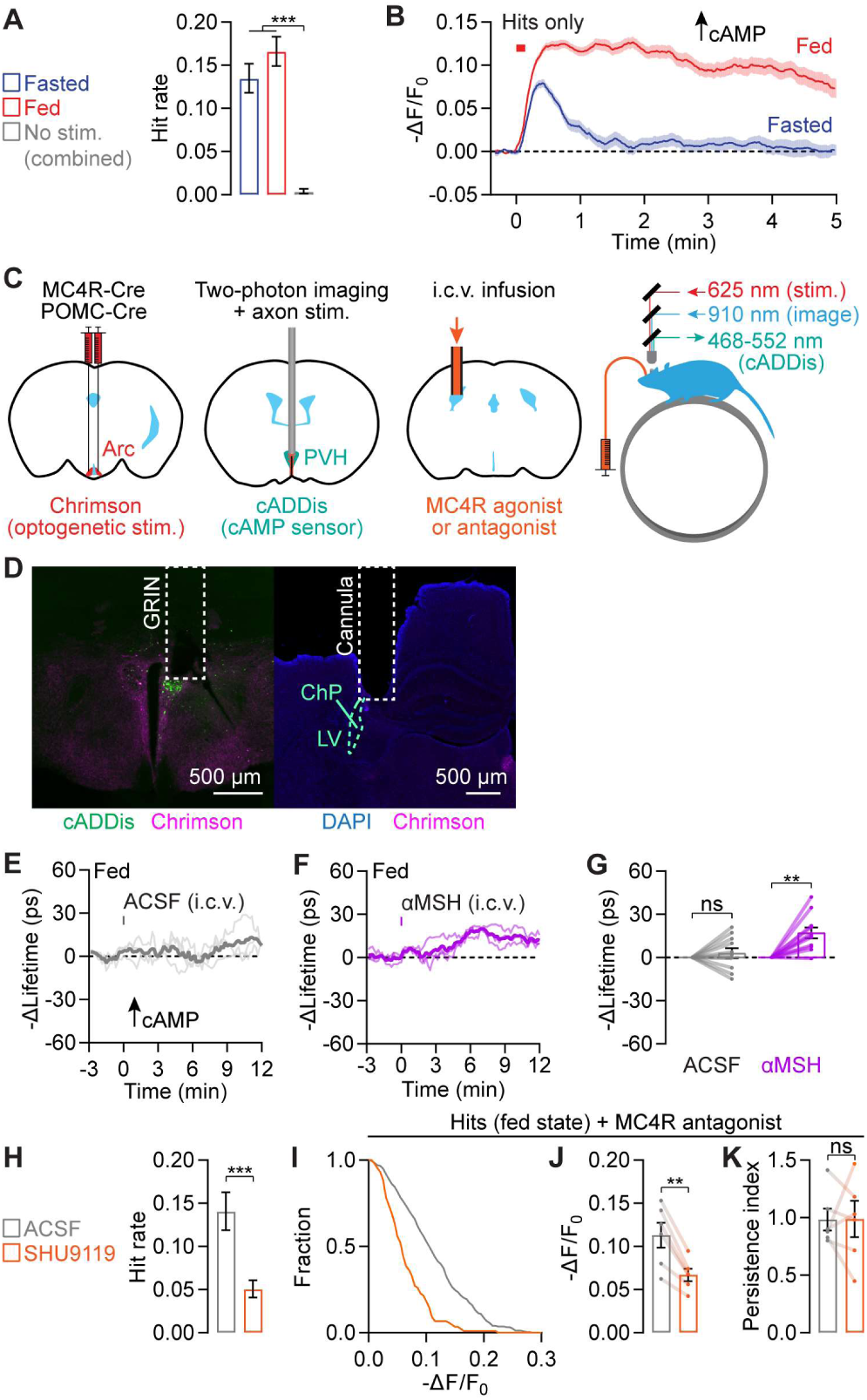
Characterizing αMSH signaling to PVH^MC4R^ neurons. **(A)** Mean hit rates of POMC axon stimulation–induced cAMP increments in PVH^MC4R^ neurons are not different between fasted and fed states. No-stimulation condition in both fasted and fed states shows a low hit rate. n = 1471-3176 trials from 4 mice. Bootstrap comparison of hit rates. Mean ± 95% C.I. **(B)** cAMP increments last more than 5 min in fed mice and 1-2 min in fasted mice. n = 211-304 hits from 4 mice. **(C)** Complete surgical setup for combining i.c.v. infusion, optogenetic axon stimulation, and cAMP recording through a GRIN lens. Chrimson is expressed in POMC neurons in the arcuate nucleus of the hypothalamus (Arc), and cADDis is virally expressed in PVH^MC4R^ neurons. A GRIN lens is placed above PVH, and a cannula is placed in the posterior lateral ventricle. We chose i.c.v. infusion of peptide antagonist/agonists here and below because many of these drugs do not cross the blood-brain barrier. **(D)** Histology of cADDis expression (green) in PVH^MC4R^ neurons and Chrimson-tdTomato expression (red) in POMC axons. In the left panel, dotted lines delineate GRIN lens track. In the right panel, white dotted line delineates the anterior side of the infusion cannula inserted in the lateral ventricle (LV; cyan dotted line) which contains choroid plexus (ChP). **(E-F)** Single-mouse fluorescence lifetime traces of cADDis in PVH^MC4R^ neurons in response to infusion of ACSF (E) or 1 nmol αMSH (F). n = 3 mice. **(G)** Single field-of-view summary of cAMP increase in PVH^MC4R^ neurons in response to αMSH infusion (n = 11 FOVs from 3 mice). **(H)** Pre-infusing MC4R antagonist SHU9119 (1 nmol) reduced the hit rate of POMC stimulation–induced cAMP increments in PVH^MC4R^ neurons (n = 931-1757 trials from 2 mice). **(I)** Cumulative distribution of the magnitude of single-trial cAMP increments (measured with cADDis) indicates a reduction in magnitude when SHU9119 was pre-infused before optogenetic stimulation of POMC axons (n = 88-130 hits from 2 mice). **(J-K)** Pre-infusing SHU9119 reduced the magnitude of POMC stimulation–induced cAMP increments on hit trials without affecting cAMP persistence (n = 6 FOVs from 2 mice, t-test).

**Figure S3.**
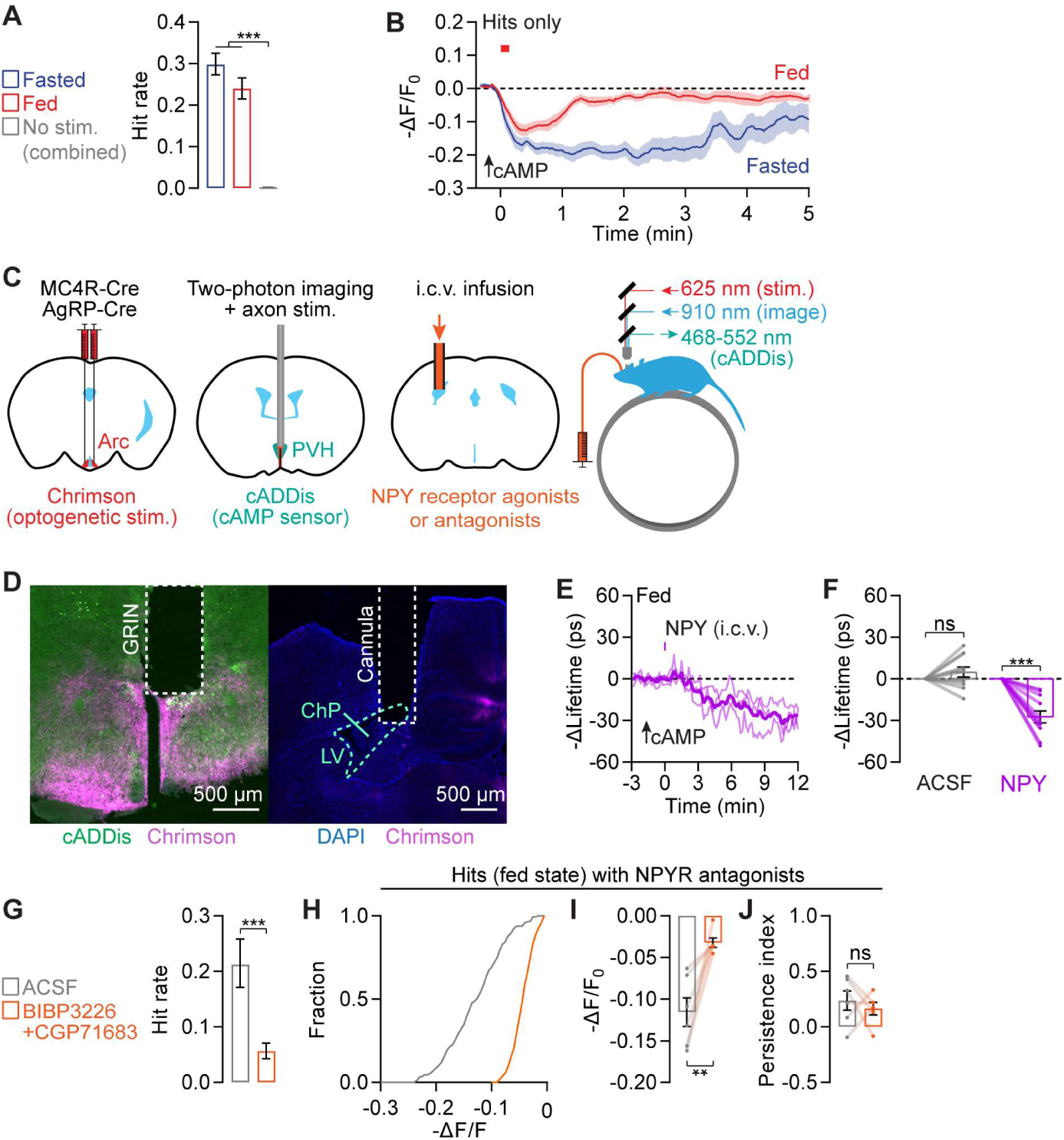
Characterizing NPY signaling to PVH^MC4R^ neurons. **(A)** Mean hit rates of AgRP axon stimulation–induced cAMP decrements in PVH^MC4R^ neurons are not different between fasted and fed states. No-stimulation condition in both fasted and fed states shows a low hit rate. n = 1093-1928 trials from 4 mice. Bootstrap comparison of hit rates. Mean ± 95% C.I. **(B)** cAMP decrements last more than 5 min in fasted mice and 1-2 min in fed mice. n = 262-355 hits from 4 mice. **(C)** Complete surgical setup for combining i.c.v. infusion, optogenetic axon stimulation, and cAMP recording through the GRIN lens. Chrimson is virally expressed in AgRP neurons in the arcuate, and cADDis is virally expressed in PVH^MC4R^ neurons. A GRIN lens is placed above PVH, and a cannula is placed in the posterior lateral ventricle. **(D)** Histology of cADDis expression (green) in PVH^MC4R^ neurons and Chrimson-tdTomato expression (red) in AgRP axons. In the left panel, dotted lines delineate the GRIN lens track. In the right panel, white dotted line delineates the anterior side of the infusion cannula inserted in the lateral ventricle (LV; cyan dotted line) which contains choroid plexus (ChP). **(E)** Single-mouse fluorescence lifetime traces of cADDis in PVH^MC4R^ neurons in response to infusion of 0.5 nmol NPY. n = 3 mice. **(F)** Single field-of-view summary of decrease in cAMP in PVH^MC4R^ neurons in response to infusion of 0.5 nmol NPY (n = 11 FOVs from 3 mice). **(G)** Pre-infusing NPY1R antagonist BIBP3226 (10 nmol) together with NPY5R antagonist CGP71683 (10 nmol) reduced the hit rate of AgRP stimulation–induced cAMP decrements in PVH^MC4R^ neurons (n = 680-909 trials from 2 mice). **(H)** Cumulative distribution of magnitudes of single-trial cAMP decrements (measured with cADDis) indicates a decrease in magnitudes when BIBP3226 and CGP71683 were pre-infused before optogenetic stimulations (n = 55-144 hits from 2 mice). **(I-J)** Pre-infusing BIBP3226 and CGP71683 reduced the magnitude of AgRP stimulation–induced cAMP decrements without affecting the persistence of these cAMP decrements (n = 6 FOVs from 2 mice, t-test).

**Figure S4.**
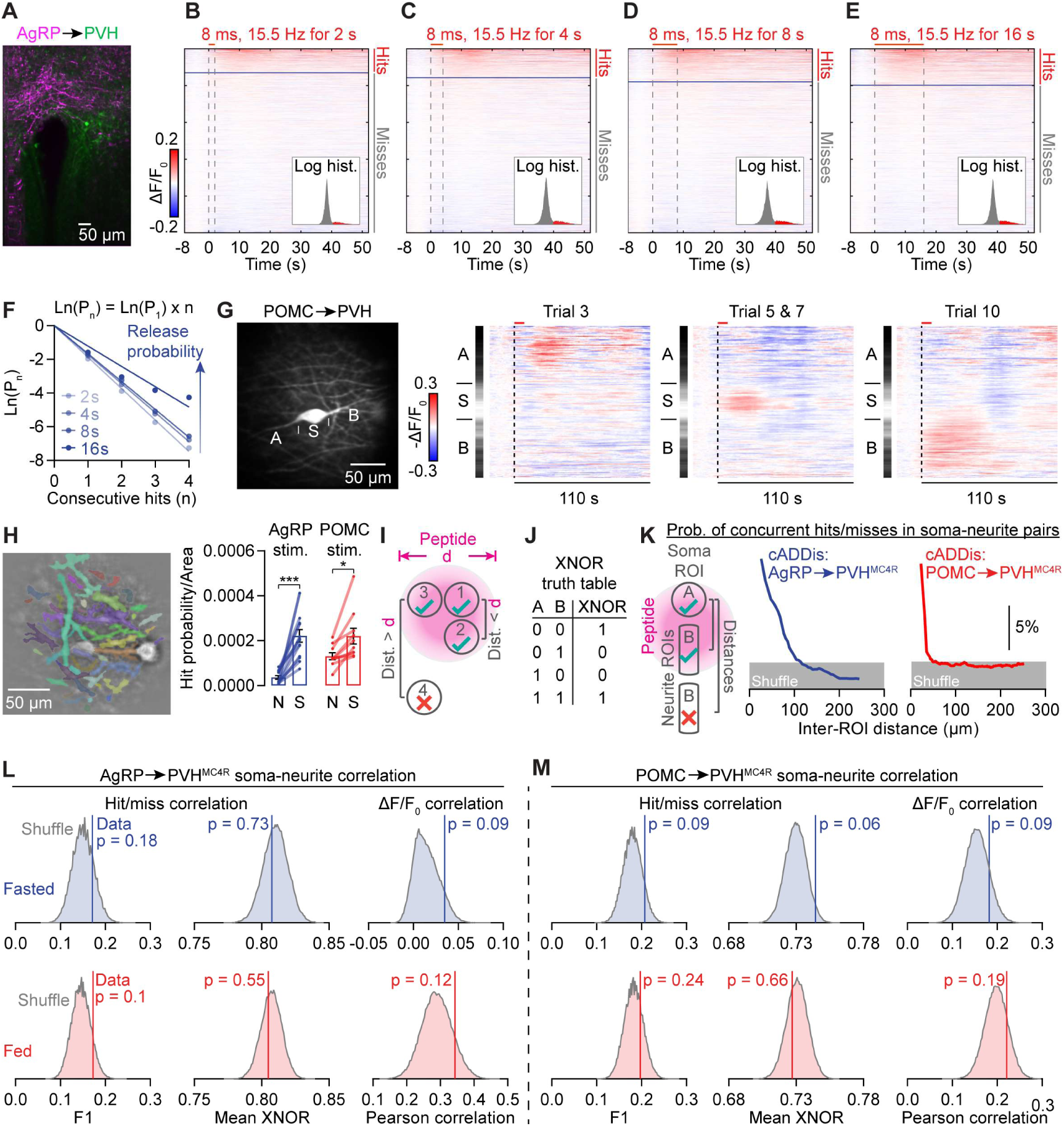
Stochastic neuropeptide release triggers spatially discrete cAMP signals. **(A)** Example field view of npyLight expression in PVH (green) and Chrimson-tdTomato expression in AgRP axons (red). **(B-E)** Summary heatmaps of single-trial npyLight signals in response to 2 s (B), 4 s (C), 8 s (D), or 16 s (E) of Chrimson photostimulation (n = 16439-19969, trials from 7 mice). Trials are sorted by peak intensity. Inset: distribution of peak intensities, color-coded red for hits and gray for misses, with x-axis on a log scale. **(F)** The hit rates of AgRP stimulation–induced NPY signals (measured with npyLight) are mostly well described using the dice model, with a modest, sublinear increase in probability of release with increasing stimulation duration (n = 16439-19969 trials from 7 mice). **(G)** In an example neuron with a clearly visible soma (S) and two associated neurites (A and B), POMC stimulation–induced cAMP increments are spatially localized and occur on different trials for the soma and for each neurite. **(H)** Left: example neurite segmentation by Cellpose 2.0 nuclear model that is retrained by manual neurite segmentation (see Methods). Right: During both AgRP axon stimulation and POMC axon stimulation, the hit rate per area is non-zero in neurites (N) but is lower in neurites than soma (S; n = 10-13 FOVs from 8 mice, one-way ANOVA). **(I)** Model: during spatially restricted peptide release, ROIs that are closer to each other than the impact diameter are more likely to receive the same peptide signal than ROIs that are further apart. **(J)** The truth table of exclusive not-or (XNOR), a metric of concurrence between pairs of binary events, A and B. When cAMP responses in two regions-of-interest concur on a given trial (e.g. both exhibit a hit or both exhibit a miss), the XNOR value equals one. **(K)** Probability of concurrent hits or misses between soma-neurite pairs drops as the distance between the two ROIs increases. The distance beyond which concurrence of cAMP responses drops to chance levels is ∼100 μm during both AgRP stimulation (left) and POMC stimulation (right). **(L)** During AgRP axon stimulation in both fasted (blue) and fed (red) states, the soma-neurite signals are consistently decorrelated from each other. This is evident using F1-score analysis of binary hit/miss data, XNOR analysis of binary hit/miss data, or Pearson correlation of continuous ΔF/F_0_ data. Vertical lines are experimental data, and shaded area shows the bootstrapped distribution (100,000 iterations, with replacement). P-value indicates two-tailed probability that actual concurrence estimate (F1, XNOR or Pearson correlation) falls outside the mean of shuffled values. **(M)** Same as L, but for POMC axon stimulation.

**Figure S5.**
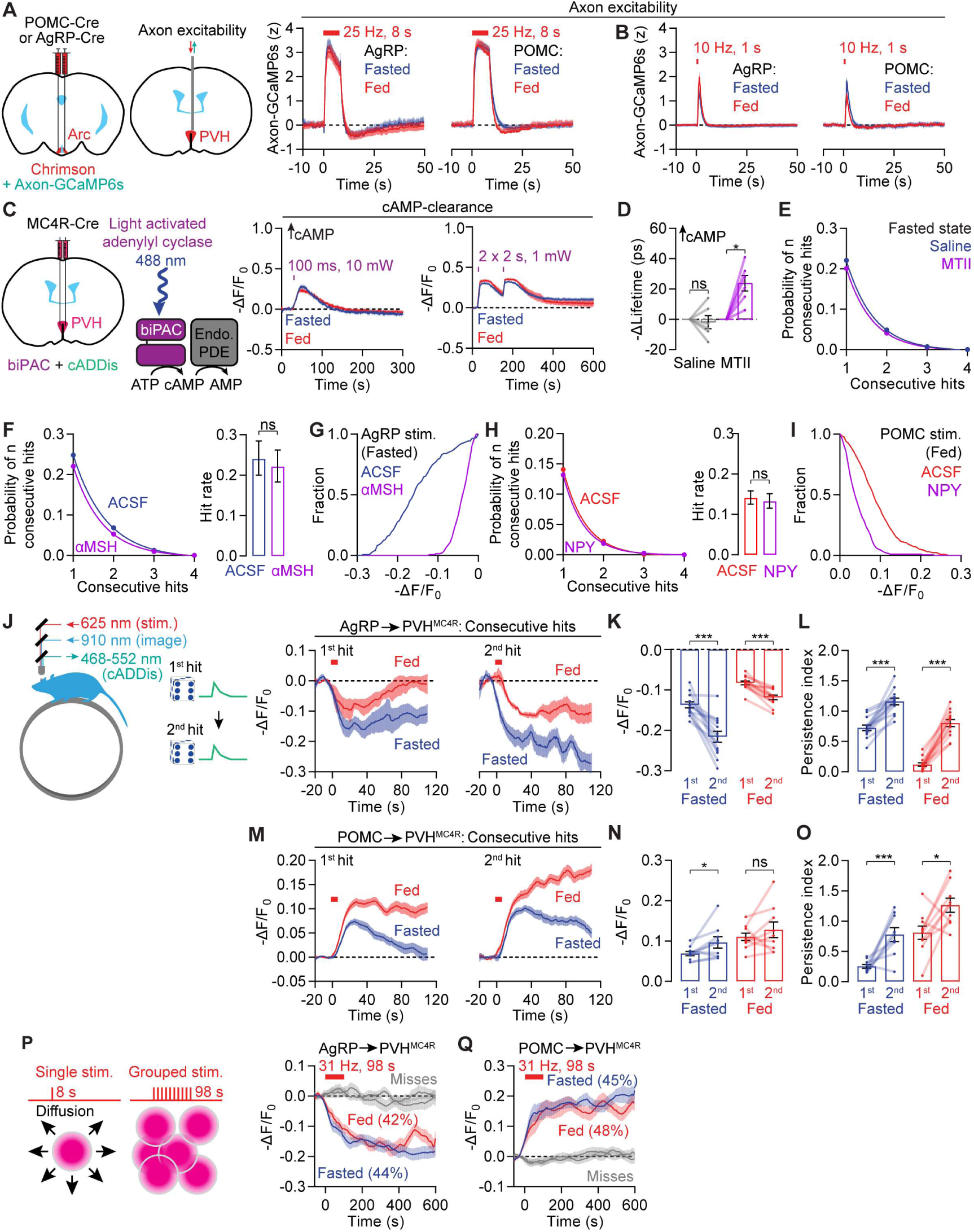
Additional evidence for the neuropeptide competition hypothesis. **(A)** To measure potential axon excitability differences across states, which could lead to differences in peptide release, we first co-expressed Chrimson and Axon-GCaMP6s in AgRP neurons and, in separate experiments, in POMC neurons. In both cases, we implanted an optic fiber in PVH to photostimulate Chrimson-expressing axons and to record the resulting calcium transients in these same axons. Axon calcium transients evoked by 25-Hz, 8-s Chrimson stimulation in AgRP (left) and POMC axons (right) were similar in fasted and fed states (n = 9 mice). **(B)** Brief 10 Hz, 1 s Chrimson stimulation of AgRP axons triggers stronger calcium transients in the fed state (when NPY signaling is weaker). The same photostimulation of POMC axons triggers stronger calcium transients in the fasted state (when αMSH signaling is weaker). These differences are presumably due to lower baseline activity of AgRP axons and POMC axons in the fed and fasted state, respectively. n = 9 mice. Note that these differences argue against a major contribution of presynaptic excitability to the state-dependent differences in cAMP responses: AgRP stimulation drives weaker PVH^MC4R^ cAMP responses in the fed state, despite the slightly *stronger* 1-s stimulation-evoked AgRP axon calcium signals in the fed state. Similarly, POMC stimulation drives weaker PVH^MC4R^ cAMP responses in the fasted state, despite the slightly stronger 1-s stimulation-evoked POMC axon calcium signals in the fasted state. **(C-D)** We used blue-light activation of the optogenetic adenylyl cyclase, biPAC, to bypass endogenous peptide receptor signaling and directly produce cAMP in PVH^MC4R^ neurons in slices, while monitoring cAMP dynamics with the sensor cADDis. cAMP produced by biPAC photostimulation (1x 100 ms or 2x 2 s) in PVH^MC4R^ neurons is cleared by endogenous PDEs at similar rates in fasted and fed states, arguing against state-dependent cAMP degradation (n = 8 slices from 4 mice). **(D)** MTII injection (3 mg/kg, i.p.) elevates cAMP in PVH^MC4R^ neurons using two-photon fluorescence lifetime imaging *in vivo* (n = 6 FOVs from 3 mice, one-way ANOVA). **(E)** MTII pre-injection does not change the hit rate of AgRP stimulation–induced cAMP decrements (n = 473-517 trials from 3 mice). **(F-G)** αMSH pre-injection does not change the hit rate of AgRP stimulation–induced cAMP decrements (F) but reduces hit magnitudes (G). n = 391-473 trials from 2 mice, bootstrap comparison of hit rates. **(H-I)** NPY pre-infusion does not change the hit rate of POMC stimulation–induced cAMP increments (H) but reduces hit magnitudes (I). n = 1311-1667 trials from 2 mice, bootstrap comparison of hit rates. **(J-L)** When analyzing two consecutive hits of AgRP stimulation–induced cAMP decrements, the second decrement is larger than the first (n = 13 FOVs from 4 mice). Amplitudes are calculated as –ΔF/F_0_ means in the 20-40 s window following stimulation onset. Baselines are calculated separately for first and second hits to prevent lingering elevation from the first hit from contributing to the calculations of the second. **(M-O)** The same as J-L but for POMC stimulation–induced cAMP increments (n = 10 FOVs from 4 mice). **(P-Q)** To more directly manipulate the total amount of peptide released in a local region of PVH, we presented groups of ten 8-s photostimulations of AgRP and POMC axons with a much shorter inter-stimulation interval (2 s instead of 52 s; other experimental parameters were not modified). The shorter inter-stimulation intervals within each 98-s stimulation sequence should decrease the degree to which peptides released during each 8-s stimulation diffuse away or are broken down by peptidases (Xiong *et al*., 2022; Turner *et al*., 1985) between trials, resulting in greater accumulation of extracellular neuropeptide levels that could overcome endogenous competition from opposing neuropeptides (illustrated in P). Consistent with this prediction, these 98-s groups of AgRP or POMC axon stimulations drove cAMP decrements (P; 42-44% hit rate per group) and increments (Q; 45-48% hit rate) that were long-lasting (>8 min after the last pulse) and insensitive to hunger state. P: n = 112-167 trials from 4 mice, Q: n = 97-139 trials from 4 mice.

**Figure S6.**
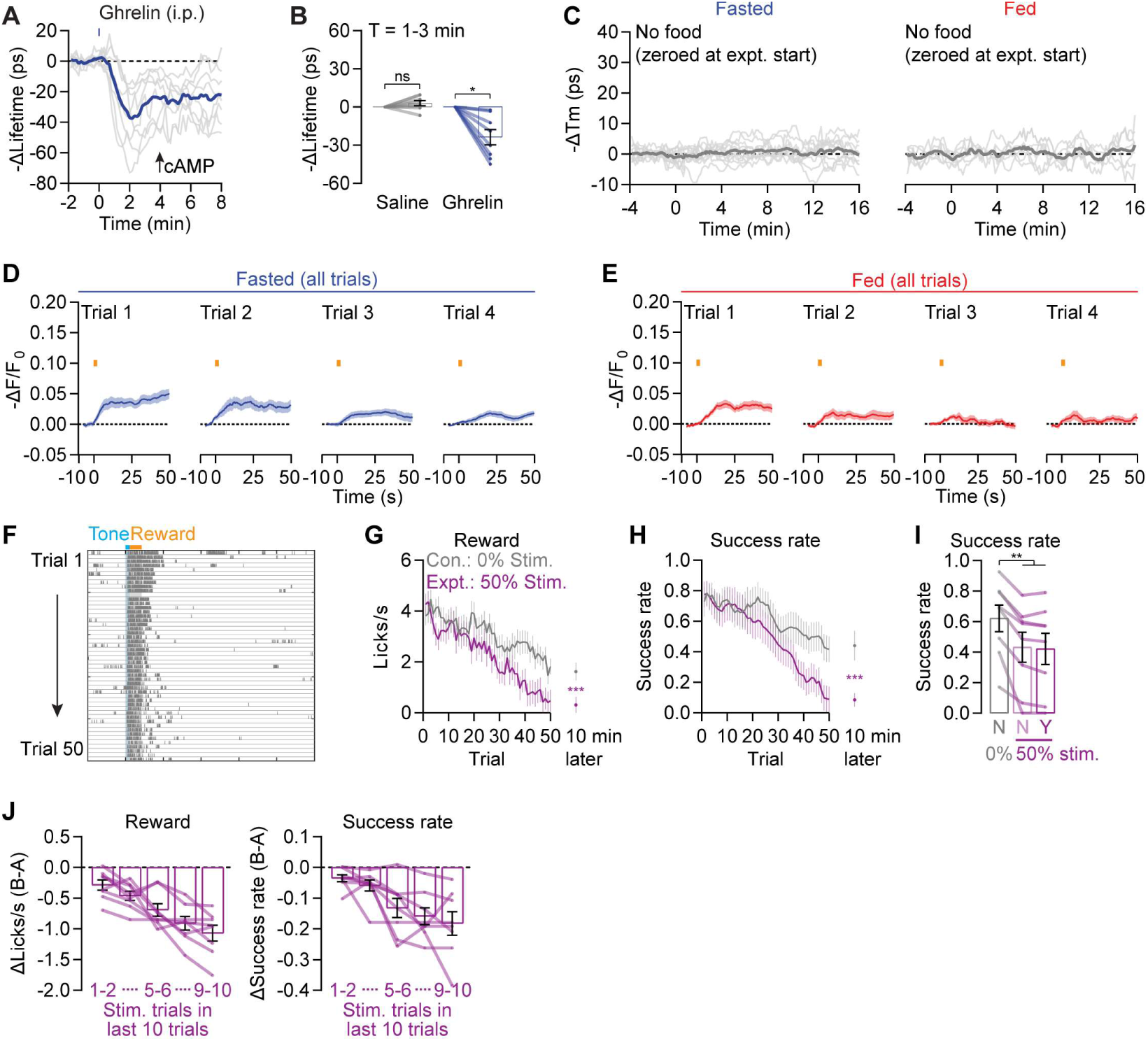
cAMP in PVH^MC4R^ neurons gradually promotes satiation. **(A-B)** Injection of ghrelin (2.5 mg/kg, i.p.), which stimulates AgRP neurons and inhibits POMC neurons, induces a robust decrease in cAMP in PVH^MC4R^ neurons in fed mice. **(C)** cAMP levels are stable in PVH^MC4R^ neurons in the absence of feeding (n = 8 FOVs from 3 mice), arguing against any non-stationarities due to elapsed time within a session. **(D-E)** Trial-average cAMP increments in the fasted (D) and fed states (E) in the first four trials. D: n = 561 trials from 6 mice, E: 646 trials. **(F)** In an assay where fasted mice lick during a tone (blue) to obtain reward (orange, milkshake), well-trained food-restricted mice start licking during the tone but the lick rate gradually decreases over 50 trials. **(G-H)** In experimental sessions (purple), in which biPAC stimulation was delivered in 50% of the trials, lick rates (G) and success rates (H) drop off faster than in control sessions (gray), and do not recover after 10 min without additional cues or reward deliveries. n = 8 mice, t-test. **(I)** Within the experimental session, no difference in success rate (rate at which the mouse exhibited licking following the tone but prior to the reward delivery) was observed between stimulation trials (purple, ‘Y’) and no-stimulation trials (purple, ‘N’). n = 8 mice, one-way ANOVA. **(J)** Over a window of 10 trials, the decrease in lick rate during reward (left), and success rate of correct licking following the cue (right) scale with the total number of biPAC stimulations delivered across the last ten trials (n = 8 mice). Together with panel I, this suggests that the effects of cAMP increments are gradual (i.e., they do not affect same-trial performance) and cumulative across minutes.

**Figure S7.**
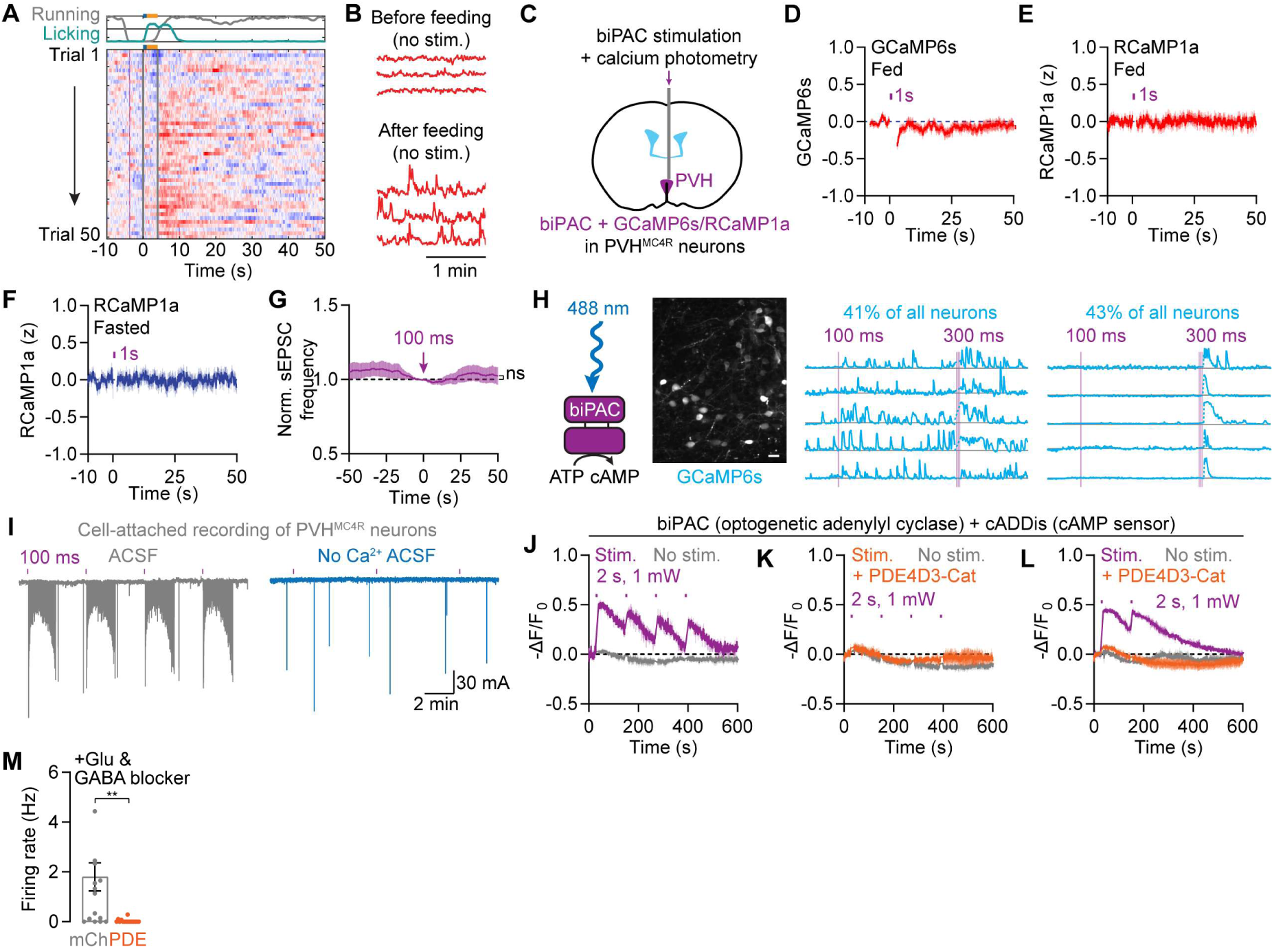
cAMP potentiates excitatory inputs to PVH^MC4R^ neurons. **(A)** An example session from a fasted mouse that licks during the cue (blue) to obtain reward (orange). Panel shows a heatmap of RCaMP1a photometry signals (each row is a trial). There is a delayed increase in calcium activity during each trial (which peaks ∼4s after cue onset, and several seconds after consumption onset), potentially reflecting ingestion-related gastrointestinal signals. This increase in activity takes ∼10 trials to develop. **(B)** Photometry recording of spontaneous bulk calcium activity after a session suggests increases in ongoing PVH^MC4R^ calcium activity after feeding. **(C-F)** In mice co-expressing biPAC and a calcium sensor (GCaMP6s or RCamp1a) in PVH^MC4R^ neurons, briefly stimulating biPAC through a fiber does not result in noticeable calcium transients in either fed mice (D and E) or fasted mice (F). We tested RCaMP1a, a green-light sensitive calcium sensor, to avoid biPAC activation by photometry light. Blanking (2-4 s) in traces is done to remove temporary photobleaching due to optogenetic stimulation. n = 8 mice per panel. **(G)** Brief 100-ms biPAC stimulation did not increase the frequency of spontaneous excitatory inputs, thereby sensitizing PVH^MC4R^ neurons to excitatory inputs. n = 16 cells from 3 fed mice. **(H)** In acute brain slices, brief biPAC activation (100 ms or 300 ms, 10 mW) resulted in changes in calcium activity that can be described in two categories: 41% of PVH^MC4R^ neurons showed a persistent elevation in calcium activity, while a different 43% showed relatively transient (∼100 s) increases in calcium activity (n = 5 slices from 2 fed mice). Because the transient activation was not seen *in vivo* (see Figure S7C-S7F), we did not pursue it further. **(I)** Example cell-attached recording shows elevated firing rate (negative deflections) for ∼2 min after each brief biPAC stimulation pulse (100 ms). The acute neuronal activation by biPAC stimulation was not seen *in vivo* (see Figure S7C-S7F). Such a difference between slice and *in vivo* results could be due to lower extracellular calcium concentration *in vivo*, as the excitability effects in slice depended on extracellular calcium (right). **(J-L)** In acute brain slices, brief biPAC activation (2 s, 1 mW) induces cAMP increments in PVH^MC4R^ neurons that gradually decay back to baseline. Co-expressing PDE4D3-Cat completely blocks biPAC-induced cAMP transients and therefore should also reduce feeding-related cAMP increments. Traces in L allow for longer time to visualize cAMP decay. n = 3-5 slices from 3 mice total. **(M)** In slices with both glutamate and GABA blockers, PDE4D3-Cat-expressing cells almost never show spontaneous spikes (n = 6-7 cells from 4 fed mice total).

