## Supplementary material for "Competition between stochastic neuropeptide signals calibrates the rate of satiation": Table S1

**Table S1. Statistical analysis and sex comparisons**

| Figure | Group | N_total | Male mice | Female mice | Mean_total | Mean_males | Mean_females | P values | F and DF | Statistical notes |
|---|---|---|---|---|---|---|---|---|---|---|
| 1C | PDE4D3-Cat | 15 mice | 12 males | 3 females | 50.8 g at Week 4 | 52.6 g | 43.4 g | p < 0.0001 | DF = 22 | Two-tailed, unpaired t-test using Week 4 data |
| | mCherry | 9 mice | 6 males | 3 females | 26.6 g at Week 4 | 30.1 g | 19.6 g | | | |
| 1D | Lean mCherry | 7 mice | 5 males | 2 females | 22.7 g | 24.1 g | 19.1 g | | | |
| | Lean PDE4D3-Cat | 6 mice | 3 males | 3 females | 29.5 g | 29.9 g | 29.1 g | Lean: p < 0.0001 | F = 116.3 | Two-tailed, unpaired one-way ANOVA with sidak post hoc |
| | Fat mCherry | 7 mice | 5 males | 2 females | 6.3 g | 6.7 g | 5.1 g | Fat: p < 0.0001 | DF = 3 | test between PDE4D3-Cat and mCherry |
| | Fat PDE4D3-Cat | 6 mice | 3 males | 3 females | 18.1 g | 17.1 g | 18.9 g | | | |
| 1E | mCherry Week 1 | 8 mice | 5 males | 3 females | 3.96 g | 4.07 g | 3.84 g | | | |
| | PDE4D3-Cat Week 1 | 6 mice | 3 males | 3 females | 5.79 g | 6.01 g | 5.42 g | | | |
| | mCherry Week 2 | 8 mice | 5 males | 3 females | 4.08 g | 4.2 g | 3.96 g | Week 1: p = 0.0006 | | |
| | PDE4D3-Cat Week 2 | 6 mice | 3 males | 3 females | 6.56 g | 6.72 g | 6.29 g | Week 2: p < 0.0001 | F = 14.2 | Two-tailed, unpaired one-way ANOVA with sidak post hoc |
| | mCherry Week 3 | 8 mice | 5 males | 3 females | 3.95 g | 4.24 g | 3.66 g | Week 3: p < 0.0001 | DF = 7 | test between PDE4D3-Cat and mCherry |
| | PDE4D3-Cat Week 3 | 6 mice | 3 males | 3 females | 6.54 g | 6.98 g | 5.8 g | Week 4: p = 0.0021 | | |
| | mCherry Week 4 | 8 mice | 5 males | 3 females | 4.24 g | 4.11 g | 4.36 g | | | |
| | PDE4D3-Cat Week 4 | 6 mice | 3 males | 3 females | 5.88 g | 5.71 g | 6.18 g | | | |
| 1F | PDE4D3-Cat | 6 mice | 3 males | 3 females | 9.72 kCal at 24 hrs | 10.11 kCal | 9.31 kCal | p = 0.02 | DF = 11 | Two-tailed, unpaired t-test using 24 hr data |
| | mCherry | 7 mice | 5 males | 2 females | 4.36 kCal at 24 hrs | 4.76 kCal | 3.36 kCal | | | |
| 1G | PDE4D3-Cat | 6 mice | 3 males | 3 females | 24.35 kCal at 24 hrs | 24.89 kCal | 23.81 kCal | p = 0.002 | DF = 11 | Two-tailed, unpaired t-test using 24 hr data |
| | mCherry | 7 mice | 5 males | 2 females | 15.59 kCal at 24 hrs | 16.29 kCal | 13.82 kCal | | | |
| 1H | PDE4D3-Cat | 6 mice | 3 males | 3 females | 14.63 kCal at 24 hrs | 14.77 kCal | 14.48 kCal | p < 0.0001 | DF = 11 | Two-tailed, unpaired t-test using 24 hr data |
| | mCherry | 7 mice | 5 males | 2 females | 11.23 kCal at 24 hrs | 11.53 kCal | 10.46 kCal | | | |
| 1I | PDE4D3-Cat | 6 mice | 3 males | 3 females | - | - | - | p < 0.0001 from the no-correlation null hypothesis | NA | Pearson correlation |
| | mCherry | 7 mice | 5 males | 2 females | - | - | - | | | |
| 1J | PDE4D3-Cat | 6 mice | 3 males | 3 females | 549 cells | 585 cells | 513 cells | p = 0.843 | DF = 11 | Two-tailed, unpaired t-test |
| | mCherry | 7 mice | 5 males | 2 females | 568 cells | 565 cells | 574 cells | | | |
| 2C | Example session | - | - | - | - | - | - | NA | NA | |
| 2D | All sessions | 3407 trials | 2 males | 2 females | - | - | - | NA | NA | |
| 2E | Fasted state | 1571 trials | 2 males | 2 females | - | - | - | NA | NA | |
| | Fed state | 1836 trials | | | - | - | - | | | |
| | Combined no-stim. | 3176 trials | | | - | - | - | | | |
| 2F | Fasted state | 1571 trials | 2 males | 2 females | - | - | - | NA | NA | |
| | Fed state | 1836 trials | | | - | - | - | | | |
| 2H | Fasted state | 1571 trials | 2 males | 2 females | - | - | - | NA | NA | |
| | Fed state | 1836 trials | | | - | - | - | | | |
| 2I | Fasted state | 211 hits | 2 males | 2 females | - | - | - | NA | NA | |
| | Fed state | 304 hits | | | - | - | - | | | |
| 2J | Fasted state | 10 FOVs | 2 males | 2 females | 0.07 | - | - | p = 0.0005 | DF = 9 | Two-tailed, paired t-test. |
| | Fed state | 10 FOVs | | | 0.12 | - | - | | | |
| 2K | Fasted state | 10 FOVs | 2 males | 2 females | 0.17 | - | - | p < 0.0001 | DF = 9 | Two-tailed, paired t-test. |
| | Fed state | 10 FOVs | | | 1 | - | - | | | |
| 2M | ACSF | 931 trials | 1 male | 1 female | - | - | - | NA | NA | |
| | SHU9119 | 1757 trials | | | - | - | - | | | |
| 2N | ACSF | 130 hits | 1 male | 1 female | - | - | - | NA | NA | |
| | SHU9119 | 88 hits | | | - | - | - | | | |
| 3C | Example session | - | - | - | - | - | - | NA | NA | |
| 3D | All sessions | 2283 trials | 2 males | 2 females | - | - | - | NA | NA | |
| 3E | Fasted state | 1190 trials | 2 males | 2 females | - | - | - | NA | NA | |
| | Fed state | 1093 trials | | | - | - | - | | | |
| | Combined no-stim. | 1928 trials | | | - | - | - | | | |
| 3F | Fasted state | 1190 trials | 2 males | 2 females | - | - | - | NA | NA | |
| | Fed state | 1093 trials | | | - | - | - | | | |
| 3G | Fasted state | 355 hits | 2 males | 2 females | - | - | - | NA | NA | |
| | Fed state | 262 hits | | | - | - | - | | | |
| 3H | Fasted state | 13 FOVs | 2 males | 2 females | -0.165 | - | - | p < 0.0001 | DF = 12 | Two-tailed, paired t-test. |
| | Fed state | 13 FOVs | | | -0.113 | - | - | | | |
| 3I | Fasted state | 13 FOVs | 2 males | 2 females | 1.064 | - | - | p < 0.0001 | DF = 12 | Two-tailed, paired t-test. |
| | Fed state | 13 FOVs | | | 0.327 | - | - | | | |
| 3J | ACSF | 680 trials | 1 male | 1 female | - | - | - | NA | NA | |
| | BIBP3226 + CGP71683 | 909 trials | | | - | - | - | | | |
| 3K | ACSF | 144 hits | 1 male | 1 female | - | - | - | NA | NA | |
| | BIBP3226 + CGP71683 | 55 hits | | | - | - | - | | | |
| 4B | NPY wash on | 10 slices | 1 male | 3 females | - | - | - | NA | NA | |
| 4C | 2s stim. | 18864 trials | 4 males | 3 females | - | - | - | NA | NA | |
| | 4s stim. | 16439 trials | | | - | - | - | | | |
| | 8s stim. | 19969 trials | | | - | - | - | | | |
| | 16s stim. | 17379 trials | | | - | - | - | | | |
| 4D | 2s stim. | 18864 trials | 4 males | 3 females | 0.14 | - | - | NA | NA | |
| | 4s stim. | 16439 trials | | | 0.169 | - | - | | | |
| | 8s stim. | 19969 trials | | | 0.187 | - | - | | | |
| | 16s stim. | 17379 trials | | | 0.207 | - | - | | | |
| 4E | 2s stim. | 18864 trials | 4 males | 3 females | - | - | - | NA | NA | |
| | 4s stim. | 16439 trials | | | - | - | - | | | |
| | 8s stim. | 19969 trials | | | - | - | - | | | |
| | 16s stim. | 17379 trials | | | - | - | - | | | |
| 4F | 2s stim. | 2632 hits | 4 males | 3 females | - | - | - | NA | NA | |
| | 4s stim. | 2782 hits | | | - | - | - | | | |
| | 8s stim. | 3736 hits | | | - | - | - | | | |
| | 16s stim. | 3595 hits | | | - | - | - | | | |
| 4H | Fasted state | 6048 trials | 2 males | 2 females | - | - | - | NA | NA | |
| | Fed state | 8053 trials | | | - | - | - | | | |
| | Combined no-stim. | 9371 trials | | | - | - | - | | | |
| 4I | Fasted state | 280 hits | 2 males | 2 females | - | - | - | NA | NA | |
| | Fed state | 255 hits | | | - | - | - | | | |
| | Combined misses | 13566 misses | | | - | - | - | | | |
| 4J | Fasted hits | 146 hits | 2 males | 2 females | - | - | - | NA | NA | Hits/misses from classification of paired soma trials. |
| | Fasted misses | 2233 misses | | | - | - | - | | | |
| | Fasted trials | 2379 trials | | | - | - | - | | | |
| | Fed hits | 191 hits | | | - | - | - | | | |
| | Fed misses | 2947 misses | | | - | - | - | | | |
| | Fed trials | 3138 trials | | | - | - | - | | | |
| 4K | Fasted state | 4356 trials | 2 males | 2 females | - | - | - | NA | NA | |
| | Fed state | 4998 trials | | | - | - | - | | | |
| | Combined no-stim. | 5122 trials | | | - | - | - | | | |
| 4L | Fasted state | 455 hits | 2 males | 2 females | - | - | - | NA | NA | |
| | Fed state | 592 hits | | | - | - | - | | | |
| | Combined misses | 8307 misses | | | - | - | - | | | |
| 4M | Fasted hits | 188 hits | 2 males | 2 females | - | - | - | NA | NA | Hits/misses from classification of paired soma trials. |
| | Fasted misses | 1162 misses | | | - | - | - | | | |
| | Fasted trials | 1350 trials | | | - | - | - | | | |
| | Fed hits | 271 hits | | | - | - | - | | | |
| | Fed misses | 1092 misses | | | - | - | - | | | |
| | Fed trials | 1363 trials | | | - | - | - | | | |
| 4N | npyLight | 72651 trials | 4 males | 3 females | - | - | - | NA | NA | |
| | AgRP stim. (soma) | 2283 trials | 2 males | 2 females | - | - | - | | | |
| | POMC stim. (soma) | 3407 trials | | | - | - | - | | | |
| 5C left | Saline | 114 hits | 2 males | 1 female | - | - | - | NA | NA | |
| | MTII | 95 hits | | | - | - | - | | | |
| 5C middle | Saline | 6 FOVs | 2 males | 1 female | -0.166 | - | - | p = 0.0013 | DF = 5 | Two-tailed, paired t-test. |
| | MTII | 6 FOVs | | | -0.092 | - | - | | | |
| 5C right | Saline | 6 FOVs | 2 males | 1 female | 1.28 | - | - | p = 0.0007 | DF = 5 | Two-tailed, paired t-test. |
| | MTII | 6 FOVs | | | 0.76 | - | - | | | |
| 5E left | ACSF | 97 hits | 1 male | 1 female | - | - | - | NA | NA | |
| | αMSH | 92 hits | | | - | - | - | | | |
| 5E middle | ACSF | 6 FOVs | 1 male | 1 female | -0.158 | - | - | p = 0.0005 | DF = 5 | Two-tailed, paired t-test. |
| | αMSH | 6 FOVs | | | -0.035 | - | - | | | |
| 5E right | ACSF | 6 FOVs | 1 male | 1 female | 1.06 | - | - | p = 0.0004 | DF = 5 | Two-tailed, paired t-test |

| Figure | Group | N_total | Male mice | Female mice | Mean_total | Mean_males | Mean_females | P values | F and DF | Statistical notes |
|---|---|---|---|---|---|---|---|---|---|---|
| [illegible] | αMSH | 6 FOVs | 1 male | 1 female | 0.18 | - | - | p = [illegible] | [illegible] | Two-tailed, paired t-test. |
| 5G left | ACSF | 235 hits | 1 male | 1 female | - | - | - | NA | NA | |
| | NPY | 173 hits | | | - | - | - | | | |
| 5G middle | ACSF | 6 FOVs | 1 male | 1 female | 0.103 | - | - | p = 0.026 | DF = 5 | Two-tailed, paired t-test. |
| | NPY | 6 FOVs | | | 0.047 | - | - | | | |
| 5G right | ACSF | 6 FOVs | 1 male | 1 female | 0.87 | - | - | p = 0.0005 | DF = 5 | Two-tailed, paired t-test. |
| | NPY | 6 FOVs | | | 0.15 | - | - | | | |
| 6B | Fasted state | 6 mice | 3 males | 3 females | - | - | - | NA | NA | |
| | Fed state | 6 mice | | | - | - | - | | | |
| 6C | Fasted state | 11 FOVs | 3 males | 3 females | - | - | - | NA | NA | |
| | Fed state | 7 FOVs | | | - | - | - | | | |
| 6D | Fasted state | 561 trials | 3 males | 3 females | - | - | - | NA | NA | |
| | Fed state | 646 trials | | | - | - | - | | | |
| 6E | Fasted state | 1542 trials | 3 males | 3 females | - | - | - | NA | NA | |
| | Fed state | 1759 trials | | | - | - | - | | | |
| 6F | Fasted state (hits) | 359 hits | 3 males | 3 females | - | - | - | NA | NA | |
| | Fasted state (misses) | 1744 misses | | | - | - | - | | | |
| | Fasted state (trials) | 2103 trials | | | - | - | - | | | |
| 6G | Fasted state (hits) | 290 hits | 3 males | 3 females | - | - | - | NA | NA | |
| | Fasted state (misses) | 2115 misses | | | - | - | - | | | |
| | Fasted state (trials) | 2405 trials | | | - | - | - | | | |
| 6H | Saline + No opto. (A) | 17 mice | 9 males | 8 females | 0.77 g at 3 hrs | 0.9 g | 0.63 g | p < 0.0001 (A - B) | F = 22.4 | Two-tailed, unpaired one-way ANOVA with sidak post hoc test between pairs of data points at 3 hrs |
| | CNO only (B) | 17 mice | 9 males | 8 females | 0.4 g at 3 hrs | 0.42 g | 0.38 g | p = 0.0004 (A - C) | DF = 3 | |
| | Opto. Only (C) | 11 mice | 6 males | 5 females | 0.42 g at 3 hrs | 0.45 g | 0.39 g | p < 0.0001 (A - D) | | |
| | Saline + Opto. (D) | 17 mice | 9 males | 8 females | 0.18 g at 3 hrs | 0.21 g | 0.16 g | p = 0.998 (B - C) | | |
| | | | | | | | | p = 0.018 (B - D) | | |
| | | | | | | | | p = 0.032 (C - D) | | |
| 6I | No light | 8 mice | 4 males | 4 females | - | - | - | NA | NA | |
| | Light | 8 mice | | | - | - | - | | | |
| 6K | Control session | 8 mice | 4 males | 4 females | 1.72 licks/s at the end | 1.88 licks/s | 1.56 licks/s | p < 0.0001 | DF = 7 | Two-tailed, paired t-test with data after 10 min wait (10 trials) |
| | Expt. Session | 8 mice | | | 0.29 licks/s at the end | 0.33 licks/s | 0.25 licks/s | | | |
| 6L left | Control session | 8 mice | 4 males | 4 females | 2.46 licks/s | 2.67 licks/s | 2.24 licks/s | p = 0.0037 (control - no stim.) | F = 19.3 | Two-tailed, unpaired one-way ANOVA with sidak post hoc test |
| | Expt. (no stim.) | 8 mice | | | 1.41 licks/s | 1.71 licks/s | 1.10 licks/s | p = 0.0065 (control - stim.) | DF = 2 | |
| | Expt. (stim.) | 8 mice | | | 1.28 licks/s | 1.38 licks/s | 1.18 licks/s | p = 0.646 (no stim. - stim.) | | |
| 6L right | Control session | 8 mice | 4 males | 4 females | 3.03 licks/s | 3.23 licks/s | 2.84 licks/s | p = 0.0002 (control - no stim.) | F = 53.3 | Two-tailed, unpaired one-way ANOVA with sidak post hoc test |
| | Expt. (no stim.) | 8 mice | | | 2.18 licks/s | 2.49 licks/s | 1.85 licks/s | p < 0.0001 (control - stim.) | DF = 2 | |
| | Expt. (stim.) | 8 mice | | | 2.02 licks/s | 2.38 licks/s | 1.65 licks/s | p = 0.389 (no stim. - stim.) | | |
| 6M | Expt. Session | 8 mice | 4 males | 4 females | - | - | - | NA | NA | |
| 7A | RCaMP1a (no stim.) | 8 mice | 4 males | 4 females | - | - | - | NA | NA | |
| | RCaMP1a (biPAC stim.) | 8 mice | 4 males | 4 females | - | - | - | | | |
| | GCaMP6s (PDE4D3-Cat) | 8 mice | 4 males | 4 females | - | - | - | | | |
| 7B | RCaMP1a (no stim.) | 8 mice | 4 males | 4 females | - | - | - | NA | NA | |
| | RCaMP1a (biPAC stim.) | 8 mice | 4 males | 4 females | - | - | - | | | |
| | GCaMP6s (PDE4D3-Cat) | 8 mice | 4 males | 4 females | - | - | - | | | |
| 7C | Example cell | - | - | - | - | - | - | NA | NA | |
| 7D | All cells | 16 cells | 3 mice | | - | - | - | NA | NA | |
| 7H | mCherry | 19 cells | 4 mice | | 3.33 Hz | - | - | p < 0.0001 | DF = 40 | Two-tailed, unpaired t-test |
| | PDE4D3-Cat | 23 cells | 4 mice | | 0.28 Hz | - | - | | | |
| 7I | mCherry | 7 cells | 4 mice | | 6.93 Hz | - | - | p = 0.0001 | DF = 11 | Two-tailed, unpaired t-test |
| | PDE4D3-Cat | 6 cells | 4 mice | | 1.69 Hz | - | - | | | |
| 7J | mCherry | 7 cells | 4 mice | | 24.2 pA | - | - | p = 0.0116 | DF = 11 | Two-tailed, unpaired t-test |
| | PDE4D3-Cat | 6 cells | 4 mice | | 15.2 pA | - | - | | | |
| S1A | PDE4D3-Cat | 15 mice | 12 males | 3 females | 178% at Week 4 | 177% | 182% | p < 0.0001 | DF = 22 | Two-tailed, unpaired t-test using Week 4 data |
| | mCherry | 9 mice | 6 males | 3 females | 104% at Week 4 | 107% | 98% | | | |
| S1B | Lean mCherry | 7 mice | 5 males | 2 females | 0.76 | 0.76 | 0.76 | Lean: p < 0.0001 | F = 264 | Two-tailed, unpaired one-way ANOVA with sidak post hoc test between PDE4D3-Cat and mCherry |
| | Lean PDE4D3-Cat | 6 mice | 3 males | 3 females | 0.6 | 0.61 | 0.59 | Fat: p < 0.0001 | DF = 3 | |
| | Fat mCherry | 7 mice | 5 males | 2 females | 0.2 | 0.21 | 0.2 | | | |
| | Fat PDE4D3-Cat | 6 mice | 3 males | 3 females | 0.37 | 0.35 | 0.39 | | | |
| S1C | PDE4D3-Cat | 8 mice | 3 males | 5 females | 1.03 cm at Week 4 | 1.3 cm | 0.86 cm | p = 0.0325 | DF = 13 | |
| | mCherry | 7 mice | 5 males | 2 females | 0.7 cm at Week 4 | 0.78 cm | 0.5 cm | | | Two-tailed, unpaired t-test using Week 4 data |
| S1D | PDE4D3-Cat | 6 mice | 3 males | 3 females | - | - | - | NA | NA | NA |
| | mCherry | 7 mice | 5 males | 2 females | - | - | - | | | |
| S1E | PDE4D3-Cat | 6 mice | 3 males | 3 females | - | - | - | NA | NA | NA |
| | mCherry | 7 mice | 5 males | 2 females | - | - | - | | | |
| S1F | PDE4D3-Cat | 6 mice | 3 males | 3 females | - | - | - | NA | NA | NA |
| | mCherry | 7 mice | 5 males | 2 females | - | - | - | | | |
| S1G | PDE4D3-Cat | 6 mice | 3 males | 3 females | - | - | - | NA | NA | NA |
| | mCherry | 7 mice | 5 males | 2 females | - | - | - | | | |
| S1H | PDE4D3-Cat | 6 mice | 3 males | 3 females | - | - | - | NA | NA | NA |
| | mCherry | 7 mice | 5 males | 2 females | - | - | - | | | |
| S1I | PDE4D3-Cat | 6 mice | 3 males | 3 females | - | - | - | NA | NA | NA |
| | mCherry | 7 mice | 5 males | 2 females | - | - | - | | | |
| S1J | PDE4D3-Cat | 6 mice | 3 males | 3 females | - | - | - | NA | NA | NA |
| | mCherry | 7 mice | 5 males | 2 females | - | - | - | | | |
| S1K | PDE4D3-Cat | 6 mice | 3 males | 3 females | - | - | - | p = 0.0006 from the no-correlation null hypothesis | NA | Pearson correlation |
| | mCherry | 7 mice | 5 males | 2 females | - | - | - | | | |
| S1L | PDE4D3-Cat | 6 mice | 3 males | 3 females | - | - | - | p < 0.0001 from the no-correlation null hypothesis | NA | Pearson correlation |
| | mCherry | 7 mice | 5 males | 2 females | - | - | - | | | |
| S1M | PDE4D3-Cat | 6 mice | 3 males | 3 females | - | - | - | p = 0.0062 from the no-correlation null hypothesis | NA | Pearson correlation |
| | mCherry | 7 mice | 5 males | 2 females | - | - | - | | | |
| S2A | Fasted state | 1571 trials | 2 males | 2 females | 0.134 | - | - | p < 0.0001 (fasted - no stim.) | NA | Two-tailed mean comparison of bootstrapped hit rates, with Bonferroni correction. |
| | Fed state | 1836 trials | | | 0.166 | - | - | p < 0.0001 (fed - no stim.) | | |
| | Combined no-stim. | 3176 trials | | | 0.003 | - | - | p = 0.22 (fasted - fed) | | |
| S2B | Fasted state | 211 hits | 2 males | 2 females | - | - | - | NA | NA | |
| | Fed state | 304 hits | | | - | - | - | | | |
| S2E | ACSF | 3 mice | 1 male | 2 females | - | - | - | NA | NA | |
| S2F | αMSH | 3 mice | 1 male | 2 females | - | - | - | NA | NA | |
| S2G | ACSF | 11 FOVs | 1 male | 2 females | 3 ps | - | - | p = 0.654 (ACSF) | F = 10.4 | Two-tailed, paired one-way ANOVA with sidak post hoc test between -3-0 min and 9-12 min means. |
| | αMSH | 11 FOVs | | | 17 ps | - | - | p = 0.002 (αMSH) | DF = 3 | |
| S2H | ACSF | 931 trials | 1 male | 1 female | - | - | - | p < 0.0001 | NA | Two-tailed mean comparison of bootstrapped hit rates. |
| | SHU9119 | 1757 trials | | | - | - | - | | | |
| S2I | ACSF | 130 hits | 1 male | 1 female | - | - | - | NA | NA | |
| | SHU9119 | 88 hits | | | - | - | - | | | |
| S2J | ACSF | 6 FOVs | 1 male | 1 female | 0.113 | - | - | p = 0.002 | DF = 5 | Two-tailed, paired t-test. |
| | SHU9119 | 6 FOVs | | | 0.067 | - | - | | | |
| S2K | ACSF | 6 FOVs | 1 male | 1 female | 0.98 | - | - | p = 0.977 | DF = 5 | Two-tailed, paired t-test. |
| | SHU9119 | 6 FOVs | | | 0.99 | - | - | | | |
| S3A | Fasted state | 1190 trials | 2 males | 2 females | 0.298 | - | - | p < 0.0001 (fasted - no stim.) | NA | Two-tailed mean comparison of bootstrapped hit rates, with Bonferroni correction. |
| | Fed state | 1093 trials | | | 0.24 | - | - | p < 0.0001 (fed - no stim.) | | |
| | Combined no-stim. | 1928 trials | | | 0 | - | - | p = 0.17 (fasted - fed) | | |
| S3B | Fasted state | 355 hits | 2 males | 2 females | - | - | - | NA | NA | |
| | Fed state | 262 hits | | | - | - | - | | | |
| S3E | ACSF | 3 mice | 1 male | 2 females | - | - | - | NA | NA | |
| S3F | ACSF | 11 FOVs | 1 male | 2 females | 5 ps | - | - | p = 0.351 (ACSF) | F = 24.7 | Two-tailed, paired one-way ANOVA with sidak post hoc test between -3-0 min and 9-12 min means. |
| | αMSH | 11 FOVs | | | -27.5 ps | - | - | p = 0.0002 (αMSH) | DF = 3 | |
| S3G | ACSF | 680 trials | 1 male | 1 female | 0.212 | - | - | NA | NA | Two-tailed mean comparison of bootstrapped hit rates. |
| | BIBP+CGP | 909 trials | | | 0.056 | - | - | | | |
| S3H | ACSF | 144 hits | 1 male | 1 female | - | - | - | NA | NA | |
| | BIBP+CGP | 55 hits | | | - | - | - | | | |
| S3I | ACSF | 6 FOVs | 1 male | 1 female | -0.116 | - | - | p = 0.0085 | DF = 5 | Two-tailed, paired t-test. |
| | BIBP+CGP | 6 FOVs | | | 0.032 | - | - | | | |
| S3J | ACSF | 6 FOVs | 1 male | 1 female | 0.98 | - | - | p = 0.571 | DF = 5 | Two-tailed, paired t-test. |
| | BIBP+CGP | 6 FOVs | | | 0.99 | - | - | | | |

| Figure | Group | N_total | Male mice | Female mice | Mean_total | Mean_males | Mean_females | P values | F and DF | Statistical notes |
|---|---|---|---|---|---|---|---|---|---|---|
| S4B | 2s stim. | 18864 trials | 4 males | 3 females | - | - | - | NA | NA | |
| S4C | 4s stim. | 16439 trials | 4 males | 3 females | - | - | - | NA | NA | |
| S4D | 8s stim. | 19969 trials | 4 males | 3 females | - | - | - | NA | NA | |
| S4E | 16s stim. | 17379 trials | 4 males | 3 females | - | - | - | NA | NA | |
| | 2s stim. | 18864 trials | | | - | - | - | | | |
| S4F | 4s stim. | 16439 trials | 4 males | 3 females | - | - | - | NA | NA | |
| | 8s stim. | 19969 trials | | | - | - | - | | | |
| | 16s stim. | 17379 trials | | | - | - | - | | | |
| S4G | Example session | - | - | - | - | - | - | NA | NA | |
| | AgRP stim. (neurite) | 13 FOVs | 2 males | 2 females | 0.00005 | - | - | | | |
| S4H | AgRP stim. (soma) | 13 FOVs | | | 0.00022 | - | - | p = 0.011 (POMC) | DF = 3 | Two-tailed, unpaired one-way ANOVA with sidak post hoc |
| | POMC stim. (neurite) | 10 FOVs | 2 males | 2 females | 0.00013 | - | - | p < 0.0001 (AgRP) | F = 15.82 | test between soma and neurites |
| | POMC stim. (soma) | 10 FOVs | | | 0.00021 | - | - | | | |
| S4K | AgRP stim. (neurite) | 14101 trials | 2 males | 2 females | - | - | - | NA | NA | |
| | POMC stim. (neurite) | 9354 trials | 2 males | 2 females | - | - | - | | | |
| | Fasted state (neurite) | 6048 trials | | | - | - | - | | | |
| S4L | Fed state (neurite) | 8053 trials | 2 males | 2 females | - | - | - | NA | NA | Two-tailed mean comparison of bootstrapped data. From 2 males and 2 females |
| | Fasted state (soma) | 1190 trials | | | - | - | - | | | |
| | Fed state (soma) | 1093 trials | | | - | - | - | | | |
| | Fasted state (neurite) | 4356 trials | | | - | - | - | | | |
| S4M | Fed state (neurite) | 4998 trials | 2 males | 2 females | - | - | - | NA | NA | Two-tailed mean comparison of bootstrapped data. From 2 males and 2 females |
| | Fasted state (soma) | 1190 trials | | | - | - | - | | | |
| | Fed state (soma) | 1093 trials | | | - | - | - | | | |
| S5A | Fasted state | 9 mice | 5 males | 4 females | - | - | - | NA | NA | |
| | Fed state | 9 mice | 5 males | 4 females | - | - | - | | | |
| S5B | Fasted state | 9 mice | 5 males | 4 females | - | - | - | NA | NA | |
| | Fed state | 9 mice | 5 males | 4 females | - | - | - | | | |
| | Fasted state | 4 slices | 2 male | 2 female | - | - | - | NA | NA | |
| S5C | Fed state | 4 slices | | | - | - | - | | | |
| | Fasted state | 8 slices | 3 males | 3 females | - | - | - | NA | NA | |
| | Fed state | 8 slices | | | - | - | - | | | |
| S5D | Saline | 6 FOVs | 2 males | 1 female | -1.8 ps | - | - | p = 0.903 (Saline) | F = 13.5 | Two-tailed, paired one-way ANOVA with sidak post hoc test |
| | MTII | 6 FOVs | | | 23.8 ps | - | - | p = 0.01 (MTII) | DF = 3 | between -3-0 min and 9-12 min means. |
| S5E | Saline | 517 trials | 2 males | 1 females | - | - | - | NA | NA | |
| | MTII | 473 trials | | | - | - | - | | | |
| S5F left | ACSF | 391 trials | 1 male | 1 female | - | - | - | NA | NA | |
| | αMSH | 417 trials | | | - | - | - | | | |
| S5F right | ACSF | 391 trials | 1 male | 1 female | - | - | - | p = 0.891 | NA | Two-tailed mean comparison of bootstrapped hit rates. |
| | αMSH | 417 trials | | | - | - | - | | | |
| S5G | ACSF | 97 hits | 1 male | 1 female | - | - | - | NA | NA | |
| | αMSH | 92 hits | | | - | - | - | | | |
| S5H left | ACSF | 1667 trials | 1 male | 1 female | - | - | - | NA | NA | |
| | NPY | 1311 trials | | | - | - | - | | | |
| S5H right | ACSF | 1667 trials | 1 male | 1 female | - | - | - | p = 0.937 | NA | Two-tailed mean comparison of bootstrapped hit rates. |
| | NPY | 1311 trials | | | - | - | - | | | |
| S5I | ACSF | 235 hits | 1 male | 1 female | - | - | - | NA | NA | |
| | NPY | 173 hits | | | - | - | - | | | |
| S5J | Fasted state | 87hit-hits | 2 males | 2 females | - | - | - | NA | NA | |
| | Fed state | 57 hit-hits | | | - | - | - | | | |
| | Fasted state (1st) | 13 FOVs | | | 0.069 | - | - | | | |
| S5K | Fasted state (2nd) | 13 FOVs | 2 males | 2 females | 0.11 | - | - | p < 0.0001 (Fasted) | F = 41.85 | Two-tailed, paired one-way ANOVA with sidak post hoc test |
| | Fed state (1st) | 13 FOVs | | | 0.096 | - | - | p = 0.0003 (Fed) | DF = 3 | between 1st and 2nd hits. |
| | Fed state (2nd) | 13 FOVs | | | 0.13 | - | - | | | |
| | Fasted state (1st) | 13 FOVs | | | 0.25 | - | - | | | |
| S5L | Fasted state (2nd) | 13 FOVs | 2 males | 2 females | 0.78 | - | - | p < 0.0001 (Fasted) | F = 92.93 | Two-tailed, paired one-way ANOVA with sidak post hoc test |
| | Fed state (1st) | 13 FOVs | | | 0.81 | - | - | p < 0.0001 (Fed) | DF = 3 | between 1st and 2nd hits. |
| | Fed state (2nd) | 13 FOVs | | | 1.27 | - | - | | | |
| S5M | Fasted state | 24 hit-hits | 2 males | 2 females | - | - | - | NA | NA | |
| | Fed state | 39 hit-hits | | | - | - | - | | | |
| | Fasted state (1st) | 10 FOVs | | | 0.069 | - | - | | | |
| S5N | Fasted state (2nd) | 10 FOVs | 2 males | 2 females | 0.11 | - | - | p = 0.04 (Fasted) | F = 4.65 | Two-tailed, paired one-way ANOVA with sidak post hoc test |
| | Fed state (1st) | 10 FOVs | | | 0.096 | - | - | p = 0.449 (Fed) | DF = 3 | between 1st and 2nd hits. |
| | Fed state (2nd) | 10 FOVs | | | 0.13 | - | - | | | |
| | Fasted state (1st) | 10 FOVs | | | 0.25 | - | - | | | |
| S5O | Fasted state (2nd) | 10 FOVs | 2 males | 2 females | 0.78 | - | - | p = 0.0009 (Fasted) | F = 18.87 | Two-tailed, paired one-way ANOVA with sidak post hoc test |
| | Fed state (1st) | 10 FOVs | | | 0.81 | - | - | p = 0.012 (Fed) | DF = 3 | between 1st and 2nd hits. |
| | Fed state (2nd) | 10 FOVs | | | 1.27 | - | - | | | |
| | Fasted state (hits) | 74 hits | | | - | - | - | | | |
| | Fasted state (misses) | 93 misses | | | - | - | - | | | |
| S5P | Fasted state (trials) | 167 trials | 2 males | 2 females | - | - | - | NA | NA | |
| | Fed state (hits) | 46 hits | | | - | - | - | | | |
| | Fed state (misses) | 66 misses | | | - | - | - | | | |
| | Fed state (trials) | 112 trials | | | - | - | - | | | |
| | Fasted state (hits) | 43 hits | | | - | - | - | | | |
| | Fasted state (misses) | 54 misses | | | - | - | - | | | |
| S5Q | Fasted state (trials) | 97 trials | 2 males | 2 females | | | | NA | NA | |
| | Fed state (hits) | 67 hits | | | - | - | - | | | |
| | Fed state (misses) | 72 misses | | | | | | | | |
| | Fed state (trials) | 139 trials | | | - | - | - | | | |
| S6A | Ghrelin | 8 FOVs | 2 males | 1 female | - | - | - | | | |
| S6B | ACSF | 8 FOVs | 2 males | 1 female | 3.1 ps | - | - | p = 0.281 (Saline) | F = 14.4 | Two-tailed, paired one-way ANOVA with sidak post hoc test |
| | Ghrelin | 8 FOVs | | | -23.7 ps | - | - | p = 0.011 (Ghrelin) | DF = 3 | between -2-0 min and 6-8 min means. |
| S6C | Fasted state | 11 FOVs | 3 males | 3 females | - | - | - | NA | NA | |
| | Fed state | 7 FOVs | | | - | - | - | | | |
| S6D | Fasted state | 561 trials | 3 males | 3 females | - | - | - | NA | NA | |
| S6E | Fed state | 646 trials | | | - | - | - | NA | NA | |
| S6F | Example session | - | - | - | - | - | - | NA | NA | |
| S6G | Control session | 8 mice | 4 males | 4 females | 1.63 licks/s at the end | 1.81 licks/s | 1.44 licks/s | p = 0.0002 | DF = 7 | Two-tailed, paired t-test with data after 10 min wait (10 trials) |
| | Expt. Session | 8 mice | | | 0.32 licks/s at the end | 0.29 licks/s | 0.35 licks/s | | | |
| S6H | Control session | 8 mice | 4 males | 4 females | 0.44 | 0.49 | 0.39 | p = 0.0007 | DF = 7 | Two-tailed, paired t-test with data after 10 min wait (10 trials) |
| | Expt. Session | 8 mice | | | 0.085 | 0.064 | 0.106 | | | |
| | Control session | 8 mice | | | 0.62 | 0.65 | 0.6 | p = 0.0016 (control - no stim.) | | |
| S6I | Expt. (no stim.) | 8 mice | 4 males | 4 females | 0.43 | 0.48 | 0.38 | p = 0.0013 (control - stim.) | F = 33.9 DF = 2 | Two-tailed, unpaired one-way ANOVA with sidak post hoc test |
| | Expt. (stim.) | 8 mice | | | 0.42 | 0.47 | 0.38 | p = 0.568 (no stim. - stim.) | | |
| S6J left | Expt. Session | 8 mice | 4 males | 4 females | - | - | - | NA | NA | |
| S6J right | Expt. Session | 8 mice | 4 males | 4 females | - | - | - | NA | NA | |
| S7A | Example session | - | - | - | - | - | - | NA | NA | |
| S7B | Example session | - | - | - | - | - | - | NA | NA | |
| S7D | GCaMP6s - Fed | 8 mice | 4 males | 4 females | - | - | - | NA | NA | |
| S7E | RCaMP1a - Fed | 8 mice | 4 males | 4 females | - | - | - | NA | NA | |
| S7F | RCaMP1a - Fasted | 8 mice | 4 males | 4 females | - | - | - | NA | NA | |
| S7G | All cells | 16 cells | 3 mice | | - | - | - | NA | NA | |
| S7H | All cells | 471 cells | 2 males | - | - | - | - | NA | NA | |
| S7I | With calcium | 35 cells | 4 mice | | - | - | - | NA | NA | |
| | Without calcium | 7 cells | 2 mice | | - | - | - | | | |
| S7J | No light | 5 slices | 1 male | 1 female | - | - | - | NA | NA | |
| | Light | 5 slices | | | - | - | - | | | |
| S7K | No light | 4 slices | 1 male | 1 female | - | - | - | NA | NA | |
| | Light | 4 slices | | | - | - | - | | | |
| | No light | 3 slices | | | - | - | - | | | |
| S7L | Light | 3 slices | 1 male | 1 female | - | - | - | NA | NA | |
| | Light + PDE4D3-Cat | 3 slices | | | - | - | - | | | |
| S7M | mCherry | 7 cells | 4 mice | | 1.80 Hz | - | - | p = 0.0038 | DF = 30 | Two-wailed, unpaired t-test |
| | PDE4D3-Cat | 6 cells | 4 mice | | 0.03 Hz | - | - | | | |
